# Resolving the “Ontogeny Problem” in Vertebrate Paleontology

**DOI:** 10.1101/2024.10.25.620216

**Authors:** James G. Napoli

## Abstract

Ontogenetic change is a major source of phenotypic variation among members of a species and is often of greater magnitude than the anatomical differences that distinguish closely related species. Ontogeny has therefore become a problematic confounding variable in vertebrate paleontology, especially in study systems distant from extant crown clades, rendering taxonomic hypothesis testing (a fundamental process in evolutionary biology) rife with difficulty. Paleontologists have adopted quantitative methods to compensate for the perception that juvenile specimens lack diagnostic apomorphies seen in their adult conspecifics. Here, I critically evaluate these methods and the assumptions that guide their interpretation using a µCT dataset comprising growth series of American and Chinese alligator. I find that several widespread assumptions are scientifically unjustifiable, and that two popular methods – geometric morphometrics and cladistic analysis of ontogeny – have unacceptably high rates of type II error and present numerous procedural difficulties. However, I also identify a suite of ontogenetically invariant characters that differentiate the living species of *Alligator* throughout ontogeny. These characters overwhelmingly correspond to anatomical systems that develop prior to (and play a signaling role in) the development of the cranial skeleton itself, suggesting that their ontogenetic invariance is a consequence of the widely conserved vertebrate developmental program. These observations suggest that the architecture of the cranium is fixed early in embryonic development, and that ontogenetic remodeling does not alter the topological relationships of the cranial bones or the soft tissue structures they house. I propose a general model for future taxonomic hypothesis tests in the fossil record, in which the hypothesis that two specimens different ontogenetic stages of a single species can be falsified by the discovery of character differences that cannot be attributed plausibly to ontogenetic variation.

## Introduction

Ontogeny has long been recognized as a key lens for establishing homology, reconstructing phylogeny, and elucidating the mechanisms by which major evolutionary transformations occurred. While ontogeny does not “recapitulate phylogeny” in the Haeckelian sense, it remains the case that heritable trait differences within and between species ultimately arise from modifications to the developmental program, and therefore that the manner in which a trait develops during ontogeny conveys valuable information about its evolutionary origin and the phylogenetic relationships of different species. Interest in ontogeny among evolutionary biologists was re-invigorated in the 1970s by Gould (1977), who demonstrated the importance of ontogeny in evolutionary theory and articulated distinct mechanisms (such as heterochrony) that could produce evolutionary change. The ensuing decades have seen the rise of evolutionary developmental biology as an interdisciplinary field, and the establishment of labs that couple data from living animals and the fossil record to study the “great transformations” in vertebrate evolution, including the evolution of jaws, the origin of tetrapods, and the mechanisms underlying anatomical novelties such as the turtle shell. The origin of birds, and with it dinosaur evolution more broadly, has been the subject of a series of landmark integrative studies clarifying (for example) the role of paedomorphosis in the generation of the avian skull (Bhullar et al. 2012, Bhullar et al. 2016), distinct patterns of developmental modularity among birds, relative to other archosaurs (Felice et al. 2019), developmental linkage between the brain and overlying dermal roof bones (Fabbri et al. 2017), the identity of the carpal ossifications in living birds and their theropod ancestors (Botelho et al. 2014), and the evolutionary transformations that ultimately created unique avian pelvic and hindlimb anatomy (Egawa et al. 2018, Egawa et al. 2022, Griffin et al. 2022).

In that time, growth and development have also become leading subjects of interest within vertebrate paleontology *sensu stricto* (viz., in studies focusing almost exclusively or entirely on the fossil record). However, in paleontology these quantities of interest have also caused longstanding confusion arising from taxonomic and systematic uncertainty. Trueman (1924) articulated the “species problem” in paleontology as difficulty in accounting for variation within populations and microevolutionary change in populations through time. The third axis of the species problem was not recognized until somewhat later. Unambiguous juvenile animals were not well-documented in the fossil record until juvenile non-avian dinosaurs were first discovered by the American Museum of Natural History Central Asiatic Expeditions at the Gobi desert localities Iren Dabasu, Öösh (=Ashile, Oshii, Oshih), and Bayn Dzak (=Flaming Cliffs), including juvenile material of the ornithischian dinosaurs *Bactrosaurus johnsoni* (Gilmore 1933), *Psittacosaurus mongoliensis* (Coombs 1980), and *Protoceratops andrewsi* (Brown and Schlaikjer 1940), which was the first nearly complete growth series of a non-avian dinosaur. As collection priorities for fossil reptiles expanded beyond the recovery of “showpiece” specimens for major universities and museums, there seems to have been a general growth in both the number of and scientific interest in juvenile specimens (Johnson 1977, Horner and Makela 1979, Coombs 1980, Galton 1982, Coombs 1986, Brinkman 1988, Horner and Weishampel 1988, Horner et al. 1992). Perhaps unsurprisingly, this period corresponds closely with resurgent interest in development as a component of evolutionary theory (e.g., Gould 1977), which is reflected in a substantial body of literature focusing on allometric scaling of skeletal proportions (Rozhdestvensky 1965, Russell 1970, Dodson 1975b, Dodson 1976, Currie and Carroll 1984) and early efforts at ontogenetic staging from fossil material (Johnson 1977, Bakker 1982, Callison and Quimby 1984, Hutchison 1984, Brinkman 1988).

The growing body of data on the ontogeny of extinct groups revealed a troubling pattern: ontogenetic changes such as allometric scaling, development of ornamental features, and change in overall body sizes meant that juvenile specimens differ significantly from putative adult conspecifics^1^. Perhaps due to the abundance of well-understood osteological maturity indicators in mammals (e.g., fusion of epiphyses, closure of cranial sutures, and dental eruption sequence), this pattern has overwhelmingly affected non-mammalian paleontology. Mook (1921) was among the first to recognize that ontogenetic variation could obscure species boundaries, and while pronounced ontogenetic differences in non-avian dinosaurs were noted by Gilmore (1933) and Brown and Schlaikjer (1940), the systematic implications of such differences were first treated extensively by Rozhdestvensky (1965). In light of prior research documenting significant ontogenetic changes and putative body size differences between species, Rozhdestvensky (1965) synonymized five sauropodomorph species in three genera, and four tyrannosaurid species in three genera, into a single taxon each – *Lufengosaurus huenei* and *Tarbosaurus bataar*, respectively. Rozhdestvensky’s work was followed by applications to the North American fossil record, which similarly reinterpreted small specimens that had been erected as holotypes as juvenile individuals pertaining to other species (Russell 1970, Dodson 1975b). This trend has largely continued as methods for determining the ontogenetic stage of fossil material have advanced (see Griffin et al. [2020] for review and commentary). As more and more specimens have been reinterpreted as skeletally immature, many species have been reinterpreted as morphologically distinct growth stages (sometimes referred to as semaphoronts or alternatively as “ontogimorphs”, a term primarily used in the dinosaur-specific literature) of other taxa.

Notable examples include the interpretation of the centrosaurine ceratopsids *Brachyceratops montanensis* and *Monoclonius crassus* as *nomina dubia*, based on juvenile materials that may pertain to other centrosaurine taxa (Sampson et al. 1997); of the chasmosaurine ceratopsians *Nedoceratops hatcheri* and *Torosaurus latus* as growth stages of *Triceratops* (Scannella and Horner 2010, Scannella and Horner 2011); of the identification of the pachycephalosaurs *Dracorex hogwartsia* and *Stygimoloch spinifer* as successively mature stages *Pachycephalosaurus wyomingensis* (Horner and Goodwin 2009); and reinterpretation of *Thespesius edmontoni*, *Edmontosaurus saskatechwanensis*, and *Anatotitan copei* as immature and fully mature stages referable to *Edmontosaurus regalis* and *E. annectens* (Campione and Evans 2011). While many of these specific conclusions are debated, these and other cases have been influential in raising awareness that ontogenetic variation may be so significant as to be mistaken for phylogenetic variation between species. Thus, the paleontological “species problem” has a third axis. Not only do populations change over time, and do individuals within populations vary, but individuals themselves also vary during their own ontogeny. This principle prompted Hennig to develop his concept of the ‘character-bearing semaphoront’ and is well-known to systematists, but continues to cause so much difficulty within vertebrate paleontology that I suggest it be referred to as the “ontogeny problem”.

The ontogeny problem is perhaps best exemplified by the dispute that has proven most professionally acrimonious and attracted the most popular attention – the status of the tyrannosauroid taxon *Nanotyrannus lancensis*, which was initially interpreted as an adult of a small-bodied species of tyrannosauroid (Gilmore 1946, Bakker et al. 1988, Schmerge and Rothschild 2016b, Schmerge and Rothschild 2016a, Longrich and Saitta 2024) or a juvenile representing an adolescent of *Tyrannosaurus rex* (Carr 1999, Carr and Williamson 2004, Brusatte et al. 2010, Brusatte et al. 2012, Bever et al. 2013, Gold and Norell 2013, Brusatte and Carr 2016, Brusatte et al. 2016, Carr et al. 2017, Voris et al. 2019, Carr 2020, McKeown et al. 2020, Woodward et al. 2020, Schroeder et al. 2021, Voris et al. 2022). The synonymy of *Nanotyrannus* with *Tyrannosaurus rex* was first suggested by Rozhdestvensky (1965), based on similarity he noted between the type specimen and the specimens he identified as juvenile *Tarbosaurus bataar* (a close relative of *Tyrannosaurus rex*), but this was not formally proposed until Carr (1999) published the first exhaustive study of tyrannosaurid cranial ontogeny, and proposed that all putatively distinct features of *Nanotyrannus* were artefacts of the immaturity of the holotype specimen. Currently, the widespread consensus is that *Nanotyrannus lancensis* is a junior synonym of *Tyrannosaurus rex* and represents an immature growth stage of that taxon.

This implies that *Tyrannosaurus rex* ontogeny was characterized by extreme morphological change, including loss of several tooth positions in both the maxilla and the dentary, development of serrations on its premaxillary teeth, loss of a ‘premaxillariform’ first maxillary tooth, resorption of ornamental crests on the lacrimals, development of ornamental bosses on the postorbitals, closure of a pneumatic recess in the quadratojugal, opening of a medial pneumatic recess in the lacrimals, and a suite of other craniodental changes that led Carr (2020) to describe its ontogeny as featuring a “secondary metamorphosis”. However, proponents of a distinct *Nanotyrannus* have been a vocal minority, and the debate has at times grown quite contentious and continues to the present, often with significant attention from the popular press.

Regardless of whether *Nanotyrannus* is or is not a juvenile *Tyrannosaurus rex*, its status matters beyond the confines of what may appear to be a simple taxonomic dispute. Such questions cast long shadows, with implications for character construction and scoring, phylogenetic analysis, and all downstream evolutionary questions that depend upon resolution of the phylogenetic relationships of the taxa of interest and the ontogenetic trajectories thereof.

*Nanotyrannus* alone is a fulcrum for many outstanding questions in dinosaur paleontology, such as the ontogenetic mechanisms underlying gigantism in tyrannosauroids (Voris et al. 2022), the dynamics of Mesozoic ecosystems (Schroeder et al. 2021, Therrien et al. 2023), and whether dinosaurs were in terminal decline prior to the K-Pg bolide impact (e.g., Brusatte et al. 2015).

Systematics underlies every question in evolutionary biology, and cases such as *Nanotyrannus* have clear broader relevance as well. Hennig articulated the principle that every individual organism should be seen as a character-bearing semaphoront of a particular ontogenetic stage of the species to which it belongs. Juveniles, by definition, have not completed development and will therefore lack apomorphies that appear late in ontogeny. Perhaps unsurprisingly, “stemward slippage” of juvenile individuals in morphological phylogenetic analyses is a widespread phenomenon (e.g., Sereno et al. 2009, Kammerer 2011, Tsuihiji et al. 2011, Campione et al. 2012, Carballido and Sander 2013, Choiniere et al. 2013, Moore et al. 2018). Integrating ontogenetic information in phylogenetic analyses has been a topic of rich discussion in its own right; suffice it to say here that morphological systematics, as employed in vertebrate paleontology, depends upon both knowledge of which individuals are juveniles of which taxa and what traits are phylogenetically variable in each study system to avoid being biased by stemward slippage of juveniles. If *Tyrannosaurus rex* experienced dramatic posthatching ontogenetic transformations, it is unlikely that it was the only species to do so. Resolution of the status of *Nanotyrannus* is therefore important for establishing expectations (and thus null hypotheses) for testing the taxonomic affinities of other specimens. Indeed, general acceptance of *Nanotyrannus* as a juvenile morph of *Tyrannosaurus* was likely itself influenced by the above-discussed proposals of extreme ontogenetic changes among ornithischians. It is important to note that the *very same* characters that some teams use as evidence in favor of *Nanotyrannus* being a juvenile *Tyrannosaurus rex* have been identified by other teams of workers as evidence that *Nanotyrannus* is a valid species, suggesting that the persistence of the ontogeny problem reflects analytical assumptions and methods to a greater degree than they reflect the body of available data.

Paleontologists agree on one point – the status of putative taxa based on juvenile holotypes can only be definitively resolved by the discovery of clearly mature specimens pertaining to the same species that are clearly not their putative synonym (e.g., an osteologically mature *Nanotyrannus* that is clearly not a *Tyrannosaurus rex*), or multi-individual bonebeds that clearly show the ontogenetic trajectory of the synonymous taxon. While these ‘Rosetta Stone’ specimens have tremendous importance and may be considered the gold standard, this is not a tenable general solution because there is no guarantee that such specimens, if they exist, will ever be found. Rather, we must scrutinize the assumptions and methods used by paleontologists to test these hypotheses from the available data, to determine what deficiencies in them allow competing groups of researchers to draw diametrically opposed conclusions from the exact same suite of character evidence. One pervasive assumption is that a given rock unit will have only one species per ecological guild (sometimes relaxed to allow multiple species per guild, but none from the same clade), under the presumption that coexistence of two or more species with an overlapping niche would be impossible. Both Rozhdestvensky (1965) and Dodson (1975b) explicitly list ecological considerations (viz., the coexistence of multiple large-bodied and closely related species) as part of their rationale for synonymizing multiple species as growth stages of others, and in my experience this assumption is now often implicit despite guiding both specimen referral and the perceived “reasonability” of taxonomic hypotheses. Implicit or explicit, this assumption is unjustifiable. Many extinct ecosystems preserve a wide range of closely related (and often large-bodied) species at even high trophic levels. Among dinosaurs, the Jurassic Morrison Formation appears to have hosted multiple species of large carnivores and a diverse assemblage of sauropods, and the Cretaceous Dinosaur Park Formation attests to a remarkably diverse fauna of coexisting multi-ton hadrosaurs, ceratopsians, ankylosaurs, and tyrannosaurs (Currie 2003b, Brown et al. 2013, Paulina Carabajal et al. 2021). Cretaceous localities in the Gobi desert of Mongolia also often preserve coexisting members of the same clade. At Ukhaa Tolgod, there is evidence of at least two dromaeosaurid species (Norell et al. 2006), two oviraptorosaur species (Clark et al. 2001), and four troodontid species (Norell et al. 2000, Norell and Hwang 2004, Pei et al. 2017); two dromaeosaurids are also present at Khulsan (Napoli et al. 2021, Turner et al. 2021), and at least two tyrannosaurids are known from the Nemegt Formation (Brusatte et al. 2009, Brusatte et al. 2012). The mammalian paleobiota of Rancho La Brea is no exception, including at least 5 large felids (*Panthera atrox*, *Panthera onca*, *Puma concolor*, *Smilodon fatalis*, and *Homotherium serum*), three ursids (*Ursus americanus*, *Ursus arctos*, and *Arctodus simus*), and 3 canids (*Aenocyon dirus*, *Canis latrans*, and *Canis lupus*), to discuss only the highest trophic levels, at which we might expect competitive exclusion to be most common.

This is to say nothing of the abundance of modern, easily observable examples co-occurrence of modern species. A full survey would be impossible to present here, but crocodilians present an instructive case study. Extant crocodilians are ecologically similar semiaquatic predators, and are known to change prey choice as they grow – a phenomenon that Dodson (1975b) specifically posited would preclude coexistence of multiple species with different body sizes (under the reasoning that juveniles of the larger species would fully overlap with the niche of the smaller). Nevertheless, coexistence of multiple, differently sized crocodilian species is the rule, rather than the exception; *Crocodylus acutus* (American crocodile) and *Crocodylus porosus* (saltwater crocodile) both share portions of their range with approximately a half-dozen other species. Nile crocodiles (*Crocodylus niloticus*) coexist with the sacred crocodile (*Crocodylus suchus*), to which are so morphologically similar that the two species were mistakenly synonymized until recently (Schmitz et al. 2003, Hekkala et al. 2011), and coexisted with the extinct *Voay robustus* in Madagascar well into historical times (Hekkala et al. 2021). A particularly sobering example is that reported by Marioni et al. (2013), who caught individuals of four caiman species (*Caiman crocodilus*, *Melanosuchus niger*, *Paleosuchus trigonatus*, and *Paleosuchus palpebrosus*) at a single locality on the Purus River.

Each was of a different ontogenetic stage, and when placed next to each other it is easy to imagine a future paleontologist mistaking their remains as constituting a growth series of one taxon. A familiar mammalian corollary example are lions and tigers, which historically shared a large overlapping range from Anatolia to the Indian subcontinent, and still coexist in the Gir Forest (despite extirpation of both taxa from much of their formerly shared range). Lions and tigers are reliably differentiated by only one cranial character (Williams et al. 2015), despite not even being sister taxa among extant pantherines (Davis et al. 2010, Mazák et al. 2011). Put simply, sympatry and similarity do not equal synonymy.

The assumption that only one species per loosely defined “clade” or “guild” should be present in any extinct fauna is not justifiable, but has led to a corollary epistemological principle: establishing that a specimen is a juvenile is often taken as equivalent to establishing that it is a juvenile *of a previously named taxon*. Clearly, this is a problematic position. Most studies that synonymize taxa as growth stages of others (e.g., Horner and Goodwin 2009, Scannella and Horner 2010, Woodward et al. 2020) lack any detailed treatment of diagnostic traits or autapomorphies present in both the adult taxon and its putative juveniles; morphological evidence, if presented at all, mostly describes the *differences* between the hypothesized growth stages, rather than derived characters justifying the inclusion of the specimens under consideration within one species. This approach is obviously problematic in that it provides no positive evidence for the taxonomic identity of the specimen(s) of interest. When the possibility that juveniles pertain to distinct taxa is addressed, it is often dismissed with an appeal to parsimony (e.g., “the simplest explanation is that these are all members of the same species”), which still fails to provide positive evidence in favor of the conclusion. The maturity of a specimen is a question separate from its identity, and these questions must be answered separately. Apomorphy-based identification of fossil material, in which the synapomorphies present in a specimen are used to justify referral to increasingly exclusive phylogenetic groups, may be employed in conjunction with osteohistological maturity assessment and stratigraphic placement of specimen to test taxonomic hypotheses. While this approach does not always allow referral to the species level, it minimizes the possibility of inadvertently creating chimeric assemblages and leverages the greatest available positive evidence (Norell 1989, Nesbitt et al. 2007, Nesbitt and Stocker 2008, Bell et al. 2010, Napoli et al. 2021). However, it remains true that juvenile animals lack the full suite of characters present in adult conspecifics, so it is understandable that many workers have placed less of a premium on morphological data and accordingly increased their emphasis on alternative lines of evidence. It is also important to note that monospecific assemblages have corroborated expectations of extreme ontogenetic change in some taxa, including the theropods *Coelophysis bauri*, which shows highly variable ontogenetic trajectories (Griffin and Nesbitt 2016), and *Limusaurus inextricabilis*, which experiences a series of pronounced transformations including a complete loss of teeth (Wang et al. 2017). Therefore, apomorphy-based identification alone is clearly insufficient to resolve this problem without some *a priori* knowledge of which characters are ontogenetically variable.

Quantitative morphological analyses offer a potential solution. When large datasets are available, they can be studied with either linear or landmark-based morphometric methods (e.g., Dodson 1975b, Dodson 1976, Forster 1996, Currie 2003a, Campione and Evans 2011, Hedrick and Dodson 2013, Maiorino et al. 2015, Evans et al. 2017, Wosik et al. 2018, Mallon et al. 2020, Powers et al. 2020). Species identity can be assessed by either the presence of multiple allometric trendlines (i.e., distinct ontogenetic trajectories) or, as is now more common, the presence or absence of multivariate morphospace clusters. Other workers prefer to emphasize discrete characters, analyzing them quantitatively under phylogenetic algorithms. This approach was developed by Brochu (1996) for developmental staging of crocodilians, and has been used to describe ontogenetic stages in an wide array of taxa (Carr and Williamson 2004, Tumarkin- Deratzian et al. 2006, Carr 2010, Longrich and Field 2012, Frederickson and Tumarkin- Deratzian 2014, Ezcurra and Butler 2015, Carr et al. 2017, Carr 2020, Foster et al. 2022), including an explicit use by Zietlow (2020) to simultaneously test taxonomic hypotheses, search for sexual dimorphism, and describe ontogenetic transformations in mosasaurs. An extension of this method, Ontogenetic Sequence Analysis (Colbert and Rowe 2008), incorporates sequence polymorphism rather than only considering a consensus tree, and while rarely applied to systematic questions it has yielded important insights into developmental variability in fossil taxa (Griffin and Nesbitt 2016).

Despite the increasing adoption of quantitative approaches to taxonomic hypothesis testing, there have been few attempts to validate how such methods perform when applied to datasets in which all specimens have a known taxonomic identity (i.e., extant datasets). This is a major gap in the literature. It is intuitively appealing to suggest that every species should form its own unique allometric trendline, morphospace cluster, or clade in a cladistic analysis, but without empirical validation of these expectations we cannot be sure that our quantitative methods are fair tests of the hypotheses of interest. Furthermore, it is rare for taxonomic decisions to be made solely on the basis of quantitative optimality criteria (e.g., diagnosing the number of clusters in morphospace using clustering analysis, rather than visual examination) – and even when such methods are used, misapplication can bias results in favor of erroneous conclusions (Carr et al. 2022, Paul et al. 2022). More commonly, results of quantitative analyses are interpreted visually to guide final taxonomic decisions, which re-introduces subjectivity and at worst may render their use, as the saying goes, using “statistics like a drunk man uses a lamp post; more for support than illumination”.

The goal of this manuscript is twofold. First, I critically evaluate morphometric and cladistic approaches that have been employed to test taxonomic hypotheses incorporating ontogenetic series, to determine whether they allow confident and repeatable quantitative species delimitation among species of known taxonomic identity. Second, I seek ontogenetically invariant characters that differ between closely related extant species, and couple them with data from developmental biology to articulate a simple model to predict whether character differences are compatible with ontogenetic variation, or are more likely to suggest that a pair of specimens pertain to different species.

## Methods

### Sampling and Study System

I use the genus *Alligator* as a study system. The genus *Alligator* contains two extant species – the American alligator (*Alligator mississippiensis*) and the Chinese alligator (*Alligator sinensis*). The American alligator is a “model archosaur” that is used frequently in paleobiological studies of non-avian dinosaurs due to its easy availability; live animals can be studied *in vivo* in zoos, and eggs and fresh animals are easily obtained commercially or from sources such as the Rockefeller National Wildlife Refuge. No comprehensive anatomical description of the American alligator yet exists, but thorough documentation of its cranial osteology, pneumatic sinus systems, and vasculature allows for detailed anatomical study (Witmer 1995, Witmer 1997, Dufeau and Witmer 2015, Porter et al. 2016, Schwab et al. 2022). For these reasons, the American alligator forms a natural choice of model system. Its closest living relative is the Chinese alligator, a smaller and critically endangered species. The skull of this animal was described briefly by Mook (1923) and described extensively by Cong et al. (1998). Only a summary of this work is presently available in English, and so many of my anatomical interpretations are based upon comparison with the American alligator. This pair of sister species is an ideal model for stress-testing paleontological approaches to species delimitation. The two taxa have never been suggested to be synonymous, and cryptic species have been identified in neither – establishing their monophyly with reasonable confidence. Chinese alligators are small relative to American alligators and retain more juvenile-like proportions (e.g., larger eyes) into adulthood, and so allow critical appraisal of the ability of these approaches to accommodate ontogenetic trajectories with different endpoints. Finally, a 70^th^ specimen was added to the analysis. This individual was a spectacled caiman (*Caiman crocodilus*). Any paleontological analysis runs the risk of including singletons or small samples of a taxon that is not recognized, and it is possible that an overriding signal of the ontogeny of better-sampled species would obscure the distinctiveness of these “interloper” specimens. Including a single spectacled caiman allows for an explicit test of our ability to detect and identify such interlopers.

### Data Acquisition

A total of 70 specimens (54 *Alligator mississippiensis*, 15 *Alligator sinensis*, and 1 *Caiman crocodilus*) were µCT scanned at the American Museum of Natural History Microscopy and Imaging Facility. This sample included all scannable individuals of *A. sinensis* and the maximal growth series of both taxa; only the largest (presumably male) *A. mississippiensis* could not be scanned because they exceeded the size of the scanning chamber. Scanning parameters (including voltage, current, exposure time, and resolution) varied depending on the size and density of the specimen, and were optimized for maximal X-ray penetration at the smallest feasible voxel size. Scans were reconstructed and stitched (when necessary) into single 16-bit TIFF image stacks before being imported into VG Studio Max for segmentation and export as surface meshes for landmarking and character scoring. Four specimens, representing terminal end-members of the ontogeny of both species, were selected for full segmentation, during which every bony element of the cranium was isolated and exported as a separate 3D mesh to facilitate detailed anatomical study and character contruction. These specimens included AMNH R 8011 (juvenile *A. mississippiensis*), AMNH R 8058 (adult *A. mississippiensis*), AMNH R 175172 (juvenile *A. sinensis*) and AMNH R 23899 (adult *Alligator sinensis*). Several scans were discarded *a posteriori* because the specimens moved during scan acquisition (making the final scan dataset blurry or distorted) or due to then-unknown challenges within the specimens (e.g., the presence of metal wire within a skull that had presumably once been a teaching or display specimen).

### Geometric Morphometrics

Geometric morphometrics (GM) is a widely used method for quantifying the shape of anatomical structures, which has been leveraged for the study of myriad evolutionary and systematic question in numerous study systems. Summarized briefly, the method uses Procrustes-aligned coordinates of anatomical landmarks in 2D or 3D space as variables for multivariate ordination analyses, which reduce the dimensionality of the dataset to reveal a morphospace. Many recent paleontological studies have used landmark data to test taxonomic hypotheses, and have gradually replaced univariate regression approaches. I therefore sought to test whether closely related modern species reliably form distinct morphometric clusters during ontogeny. My landmark sampling protocol includes 54 landmarks, consisting of type I, type II, and type III landmarks. Semilandmark curves or patches were not used. Landmark numbers and definitions are listed in Table 1, and shown graphically in Figure 1. Landmarks 1- 12 were placed in dorsal view, 13-23 in lateral view, 24-32 in posterior view, 33-46 in ventral view, and 47-54 in nonstandard views; all landmarks were placed on only the left side (except those that lie on the midline). Palatal landmarks were placed on the left anterior element to allow landmarking in young juveniles with incompletely ossified palatal bones. Specimens were landmarked in Stratovan Checkpoint. In total, 56 specimens (43 *Alligator mississippiensis*, 12 *Alligator sinensis*, 1 *Caiman crocodilus*) were landmarked. Analyses were conducted in R using the packages ‘geomorph’ (Baken et al. 2021, Adams et al. 2024) and ‘Morpho’ (Schlager 2017), following a standard pipeline in which landmark data were Procrustes-transformed before analysis using principal components analysis (PCA). Missing landmarks were estimated using the estimate.missing() function.

**Figure 1.**
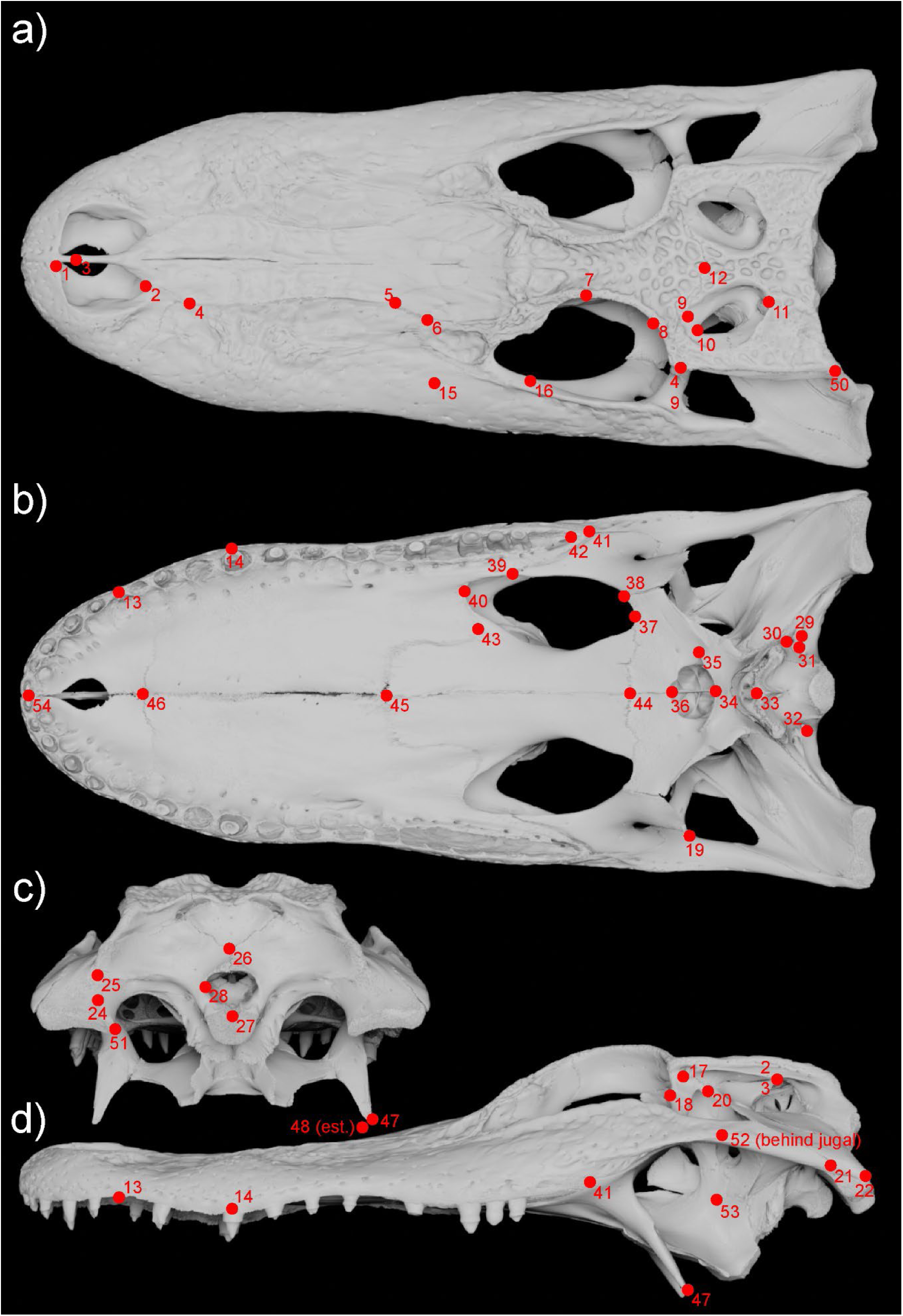
Skull of *Alligator mississippiensis* (AMNH R 8058) in a) dorsal, b) ventral, c) posterior, and d) left lateral views. Numbers denote landmark identities, defined in Table 1. Landmark 19 was digitized on the left side, but is shown on the right in ventral view for visibility. Landmark 48 is denoted as “est.” because this specimen lacks a complete pterygoid ala, and the landmark is placed in its approximate position. Landmark 52 is obscured by the jugal in lateral view.

**TABLE 1.**

| NUMBER | Definition | Type |
| --- | --- | --- |
| 1 | Anteromedial extent of external naris in dorsal view | III |
| 2 | Posterior extent of external naris in dorsal view | III |
| 3 | Anterior extent of external naris in internarial septum | II |
| 4 | Intersection of maxilla, nasal, and premaxilla (posterior extent of premaxillary posterior process) | I |
| 5 | Intersection of prefrontal, nasal, and maxilla | I |
| 6 | Intersection of prefrontal, lacrimal, and maxilla | I |
| 7 | Posterior extent of prefrontal | II |
| 8 | Intersection of postorbital-frontal suture and orbital rim | II |
| 9 | Intersection of postorbital, frontal, and parietal | I |
| 10 | Anterior extent of supratemporal fenestra | II |
| 11 | Intersection of squamosal-parietal suture with supratemporal fossa | II |
| 12 | Midline point on frontoparietal suture | III |
| 13 | Ventralmost point on premaxilla-maxilla suture | II |
| 14 | Anteroposterior midpoint of 4<sup>th</sup> maxillary alveolus | III |
| 15 | Anteriormost point on jugal | III |
| 16 | Posterior extent of lacrimal-jugal suture | II |
| 17 | Ventral extent of lateral postorbital foramen | III |
| 18 | Apex of facet for postorbital process of jugal | II |
| 19 | Dorsomedial corner of foramen at base of postorbital process of jugal | III |
| 20 | Posteroventral tip of quadratojugal process of postorbital | II |
| 21 | Posteriormost tip of jugal | II |
| 22 | Posteriormost tip of quadratojugal | II |
| 23 | Ventral tip of squamosal “hornlet” (dorsally bounding the external otic recess) | II |
| 24 | Dorsal extent of foramen aereum | III |
| 25 | Lateral extent of paroccipital process | II |
| 26 | Ventral extent of supraoccipital (contact with otoccipitals in mature individuals) | I/III |
| 27 | Midpoint of dorsal margin of occipital condyle | III |
| 28 | Lateral extent of foramen magnum | II |
| 29 | Midpoint of posterodorsal margin of foramen for jugular canal | III |
| 30 | Midpoint of posterior margin of foramen for carotid canal | III |
| 31 | Dorsolateral extent of foramen for CN XII-1 | III |
| 32 | Dorsolateral extent of foramen for CN XII-2 and CN XII-3 | III |
| 33 | Midpoint of posterior margin of median Eustachian tube ostium | III |
| 34 | Midpoint of posterior choanal ridge, or midpoint between ridges that do not contact about the midline | III |
| 35 | Lateral extent of posterior choanal ridge | II |
| 36 | Posterior extent of palatal process of pterygoid | II |
| 37 | Posterior extent of suborbital fenestra | III |
| 38 | Intersection of ectopterygoid-ptyergoid suture and suborbital fenestra | I |
| 39 | Anterior extent of maxillary process of ectopterygoid | I |
| 40 | Anterior extent of suborbital fenestra | III |
| 41 | Posterior extent of maxilla | II |
| 42 | Posterior extent of last maxillary alveolus | II |
| 43 | Posterior extent of palatine process of maxilla | II |
| 44 | Intersection of opposing pterygoids and palatines | I |
| 45 | Intersection of opposing palatines and maxillae | I |
| 46 | Intersection of opposing maxillae and premaxillae | I |
| 47 | Ventral tip of ectopterygoid | II |
| 48 | Ventral tip of pterygoid ala | II |
| 49 | Anterolateral corner of skull table | III |
| 50 | Posterolateral corner of skull table | III |
| 51 | Medial extent of quadrate condyles | II |
| 52 | Intersection of laterosphenoid and quadrate suture and trigeminal foramen | I |
| 53 | Ventral extent of pterygoid incisure | II |
| 54 | Medial extent of oral margin of premaxilla | II |

To assess the influence of taphonomic distortion, I deliberately included a specimen (AMNH R 173752; *Alligator sinensis*) with a distorted rostrum; it is unclear whether this is an anatomical variation or postmortem deformation, but in either case it simulates inclusion of distorted specimens in GM analyses. To assess the influence of incomplete sampling in the fossil record, I conducted multiple sensitivity analyses in which select individuals were deleted. One replicate included only one juvenile per taxon, one each with only juveniles of *A. mississippiensis* or *A. sinensis*, and one each with no adults of either taxon. All analyses included the single adult *C. crocodilus*.

Determination of morphospace clusters is inherently subjective, so to objectively determine whether the geometric morphometric analyses suggest the presence of distinct taxa, I subjected the PC scores for each taxon to agglomerative hierarchical clustering – a method that can search for clusters emergent within a multidimensional dataset, and report the optimal number of clusters within a dataset, and is therefore potentially useful for paleontological studies attempting to test the monospecificity of a fossil assemblage. Agglomerative hierarchical clustering was implemented via the R package ‘factoextra’ (Kassambara and Mundt 2020).

### Cladistic Analysis of Ontogeny

Because cladistic analysis of ontogeny (cladistic ontogeny hereafter) is primarily employed by a small group of researchers, it warrants brief explanation here. Brochu (1996) extensively surveyed the crocodilian postcranial skeleton and documented the sequential addition of characters throughout ontogeny. He scored these discrete characters in a phylogenetic matrix and analyzed it under parsimony-based phylogenetic methods to develop a staging scheme for determining the maturity of crocodilians that was, at least theoretically, independent of size (though given that size and maturity both increase during ontogeny, they are surely autocorrelated and are thus not truly independent). This method depends on an explicit analogy between ontogeny and evolution as proceeding from the sequential addition of characters over time. In cladistic ontogeny, shared ontogenetic characters that define successively mature growth stages are referred to as “synontomorphies”, rather than synapomorphies, with the equivalent of plesiomorphies usually simply called the “immature state”. The outgroup is an artificial operational taxonomic unit (OTU) that bears only the immature state (identified *a priori*) for each character, which is often termed an “artificial embryo”. The resulting branching diagram represents not a cladogram but an “ontogram”, and diminishingly inclusive groups correspond to progressively mature growth stages.

The method was first applied to document ontogenetic development in extinct animals by Carr and Williamson (2004), in their study of *Tyrannosaurus rex* ontogeny, and first used to address the ontogeny problem by Longrich and Field (2012), who scored individuals of both *Torosaurus* and *Triceratops* to establish that some individuals of the former were subadults at the time of death, and thus logically could not represent fully mature individuals of the latter.

Zietlow (2020) expanded the purview of cladistic ontogeny further in her study of three species of the mosasaurid squamate *Tylosaurus* from the Western Interior Seaway of North America.

This study includes an attempt to combine ontogenetic and phylogenetic data into a single analysis, and articulates explicit hypotheses regarding how an ontogram topology would reflect different scenarios. In the case that two species have morphologically similar juveniles, Zietlow (2020) proposed that the least mature specimens would resolve as successive outgroups to an ontogram “split”, after which each taxon would form its own “clade” due to divergence of ontogenetic trajectories. If juveniles were morphologically distinct or undersampled, Zietlow (2020) further predicted that the ontogram would split near the root, with no individuals plotting outside of clades corresponding to the different species. Finally, Zietlow (2020) predicted that the ontogram would show a single ladder-like topology if the two species broadly shared an ontogenetic trajectory, in which some characters would resolve as repeated “autontomorphies” of different individuals and thus provide evidence that these individuals pertained to one taxon. The same paper also proposes that these topological signals may also indicate sexual dimorphism.

These expected topological signals were applied to interpretation of Zietlow’s (2020) *Tylosaurus* data, but have not been validated empirically, and I sought to test them here.

I compiled a discrete character matrix describing cranial ontogeny and phylogenetic variation in the genus *Alligator*. Character state descriptions were based upon the four exemplar specimens subjected to full CT segmentation, which allowed the digital isolation and visualization of every bony element in the cranium, and permitted a detailed character search. To properly simulate a *de novo* search for ontogenetic characters in the fossil record, I avoided sourcing characters from the literature, though it is probable that I have independently identified characters incorporated into phylogenetic analyses of crocodylians. Character sampling was performed in the 3D modeling and animation program Blender. Each element was imported and rotated into a standardized orientation, scaled to the same size, and positioned so I could simultaneously compare all 4 exemplars of a particular element in orthographic projection. This approach completely eliminates distractions of size and parallax while conducting anatomical comparisons. First, I surveyed for differences between juveniles and adults of the two taxa, with the naïve assumption that they are of purely ontogenetic origin. I then compared the juveniles of the two species for character differences, and finally the two adults. After comparisons were made, I wrote character descriptions making sure to capture all states observed in the exemplar individuals. I followed the principles of contingent coding articulated by Brazeau (2011) to accommodate cases in which a structure appeared during ontogeny and required subsidiary characters to describe its morphology. A full character list is present in the Supplemental Information, and totals 189 characters.

Not all possible characters were constructed due to practical difficulty in scoring other specimens – for example, joint surfaces between crocodilian cranial bones tend to develop into highly interdigitate sutures during ontogeny, but external suture traces belie internal suture complexity and thus complexity cannot be fully assessed without a full CT segmentation. The *a priori* identified immature state for each character generally corresponded to that shown by juvenile *Alligator sinensis* (the smallest exemplar juvenile), except in cases where the juvenile *Alligator mississippiensis* lacked a structure present in the other juvenile. This approach mimics the paleontological implementation of cladistic ontogeny, in which juvenile character states are usually determined by comparison of multiple individuals and summarized as a composite “artificial embryo”. I further tested additional outgroup protocols to determine how they affected character polarity and the resultant ontogram topology. I ran further analyses using each juvenile exemplar as the outgroup, and a final analysis in which I used the adult *Caiman crocodilus* as an outgroup, mirroring its phylogenetic relationship to the remaining study animals. I scored 32 specimens, avoiding repetitive scoring of a large sample of juvenile *Alligator mississippiensis* with sequential specimen numbers that I presume were collected from the same nest, and yielded largely identical scores. I analyzed the cladistic ontogeny dataset in TNT v1.5 (Goloboff and Catalano 2016) under the following search parameters: multiple replicates of new technology searches until 20 hits at the shortest tree length were obtained, after which the best trees were subjected to a final iteration of tree bisection and reconnection branch swapping and zero-length branches were collapsed. Strict, 50% majority rule, and Adams consensus topologies were used to summarize the results.

### Survey for Ontogenetically Invariant Characters

The presence of ontogenetically invariant characters was first assessed via principal coordinates analysis (PCoA) of the discrete character matrix built for the prior step, based upon the pairwise Euclidean distances between specimens. Clear separation of specimens along taxonomic lines prompted manual scrutinization of the character matrix, noting any characters for which juveniles and adults of each taxon were scored identically and differently – making them ideal ontogenetically invariant diagnostic characters present in both juveniles and adults. I also noted characters that varied within taxa but did not show a clear ontogenetic signal, which I interpret as polymorphic traits that vary independently of ontogeny. Discovery of stable maxillary tooth counts prompted a targeted round of data collection from osteological specimens in the AMNH Herpetology collection, increasing sample size of *Alligator mississippiensis*, *Alligator sinensis*, *Caiman crocodilus*, and *Caiman yacare*. Skull size was measured as the width of the skull across the posterior extent of the quadratojugals, which is closely correlated with skull length in *Alligator* (Dodson 1975a).

## Results

### Geometric Morphometrics

Analysis of the full 3D landmark dataset finds that the first principal component, summarizing the majority (63.49%) of variation, is tightly correlated with centroid size, suggesting that this axis is dominated by allometric morphological changes associated with changes in skull size (Fig. 2a). Visual exploration of this morphospace confirms that this axis represents a trend towards rostral elongation, occipital and skull table flattening, and posterior growth of the quadrates and paroccipital processes that are well-documented in crocodylian ontogeny; accordingly, both *Alligator* species occupy the full range of PC1.

**Figure 2.**
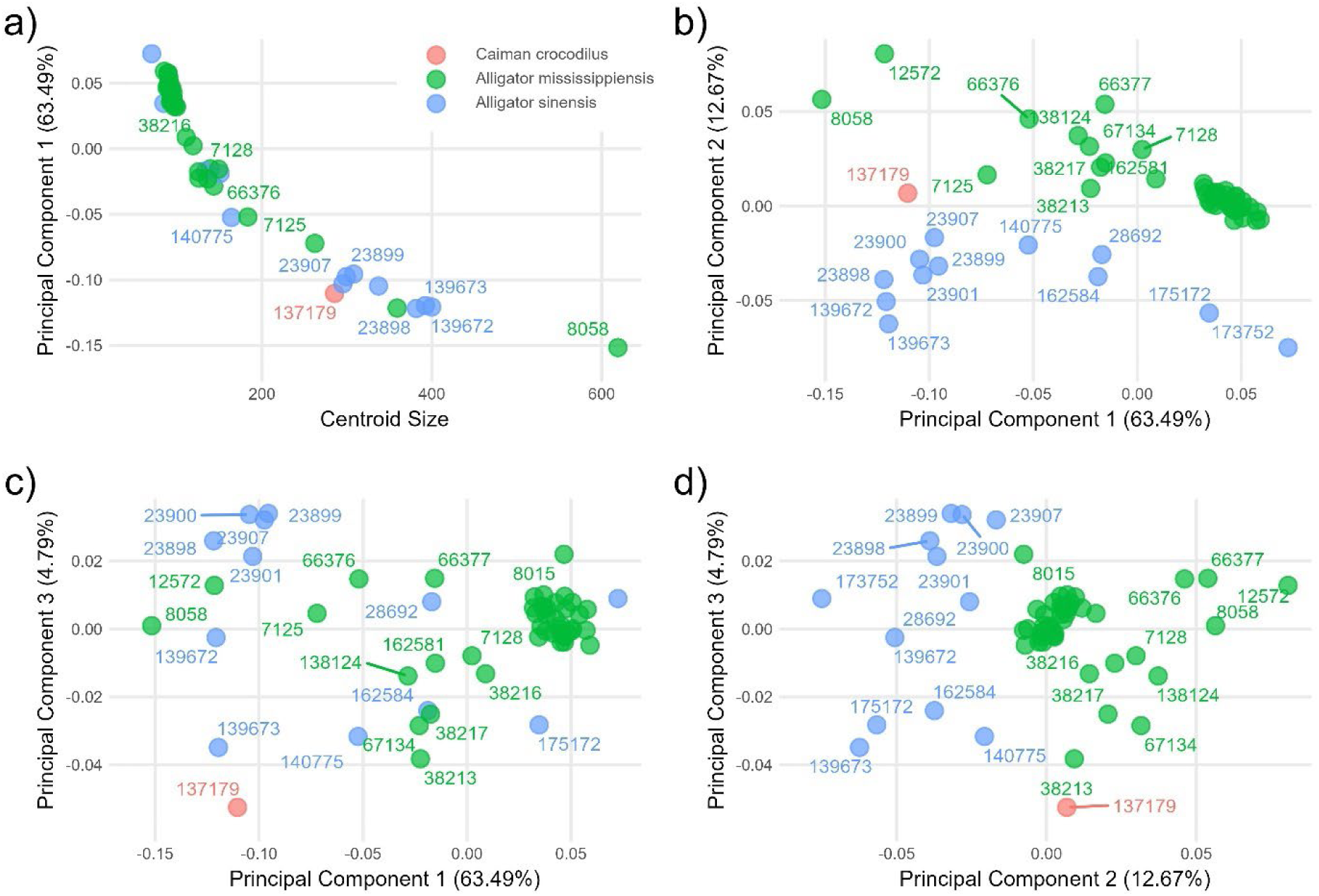
Results of geometric morphometrics analysis, including a) correlation of centroid size with principal component (PC) 1, b) PC1 vs. PC2, c) PC1 vs PC3, and d) PC2 vs PC3.

Principal component 2 describes ∼13% of shape variation, and generally separates the two *Alligator* species when plotted against PC1 (Fig 2b). PC2 primarily corresponds to the height of the posterior skull. Principal component 3 describes less than 5% of shape variation and shows wide variation within both species that is not correlated with ontogeny (Fig. 2c) or skull height (Fig 2d). *Caiman crocodilus* plots external to *Alligator* along PC3, but within *Alligator* on PC1 and PC2. Despite visually obvious separation of *Alligator mississippiensis* and *Alligator sinensis* along PC2, agglomerative hierarchical clustering fails to recover clusters corresponding to the taxa included (Fig. 3). Instead, this test suggests that the optimal solution is that there are 14 clusters within the dataset. The resultant cluster dendrogram suggests that most of these clusters correspond to an ontogenetic stage of a single taxon, but the deeper branching structure of the dendrogram does not separate species from one another, but instead groups their growth stages. The *Caiman* individual is positioned within a cluster of *Alligator sinensis* adults. Removing PC1 from the clustering analysis to remove the obviously overriding ontogenetic signal results in a cluster dendrogram (Fig. 4) that does separate the two *Alligator* species, but instead positions *Caiman crocodilus* within *Alligator mississippiensis*, and more importantly suggests that a single cluster is the optimal scenario – thus providing no statistical support for the distinction of the two *Alligator* species.

**Figure 3.**
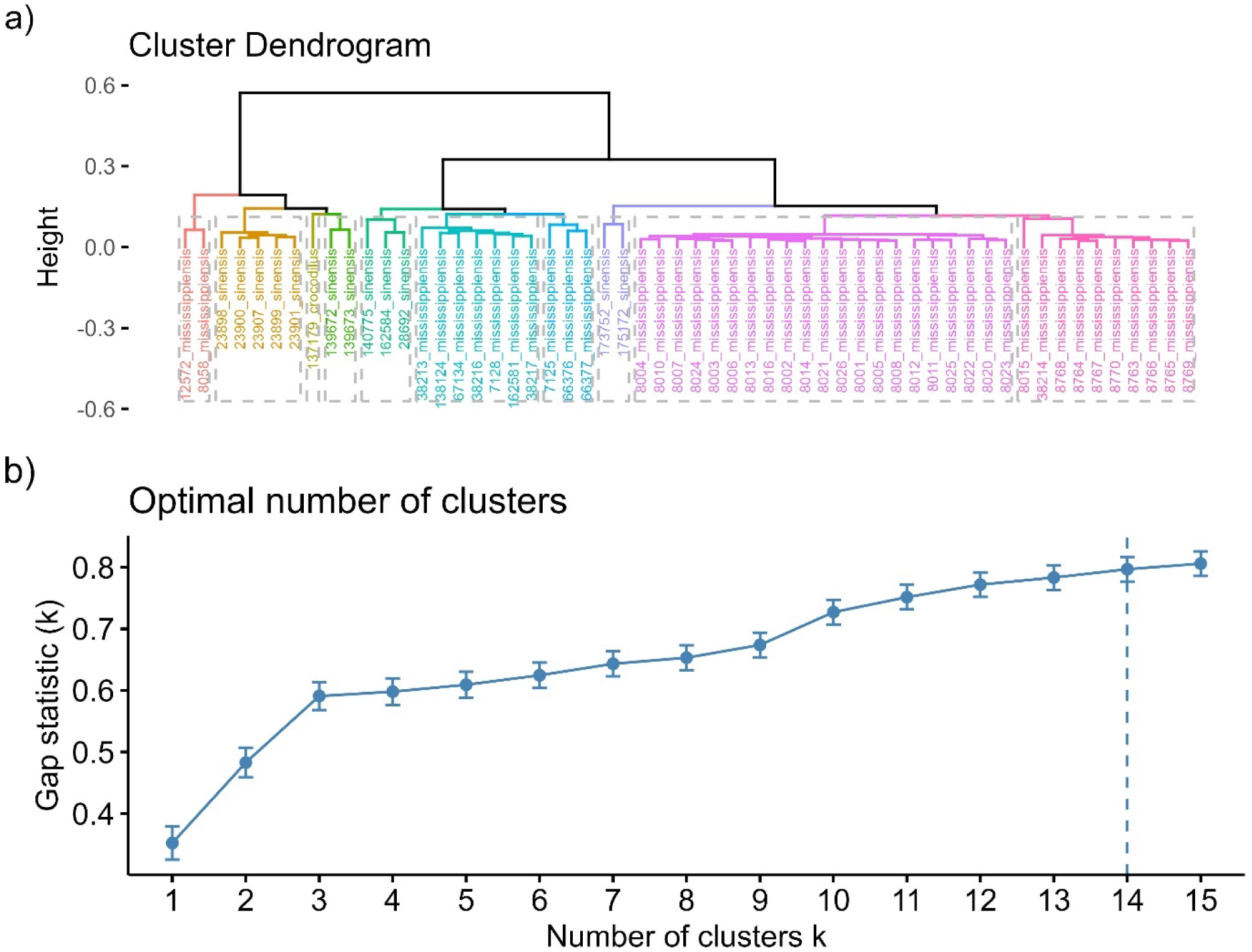
Results of agglomerative hierarchical clustering analysis on PCA scores derived from the geometric morphometrics analysis, showing a) the cluster dendrogram and b) gap statistic K indicating the optimal number of clusters in the dataset.

**Figure 4.**
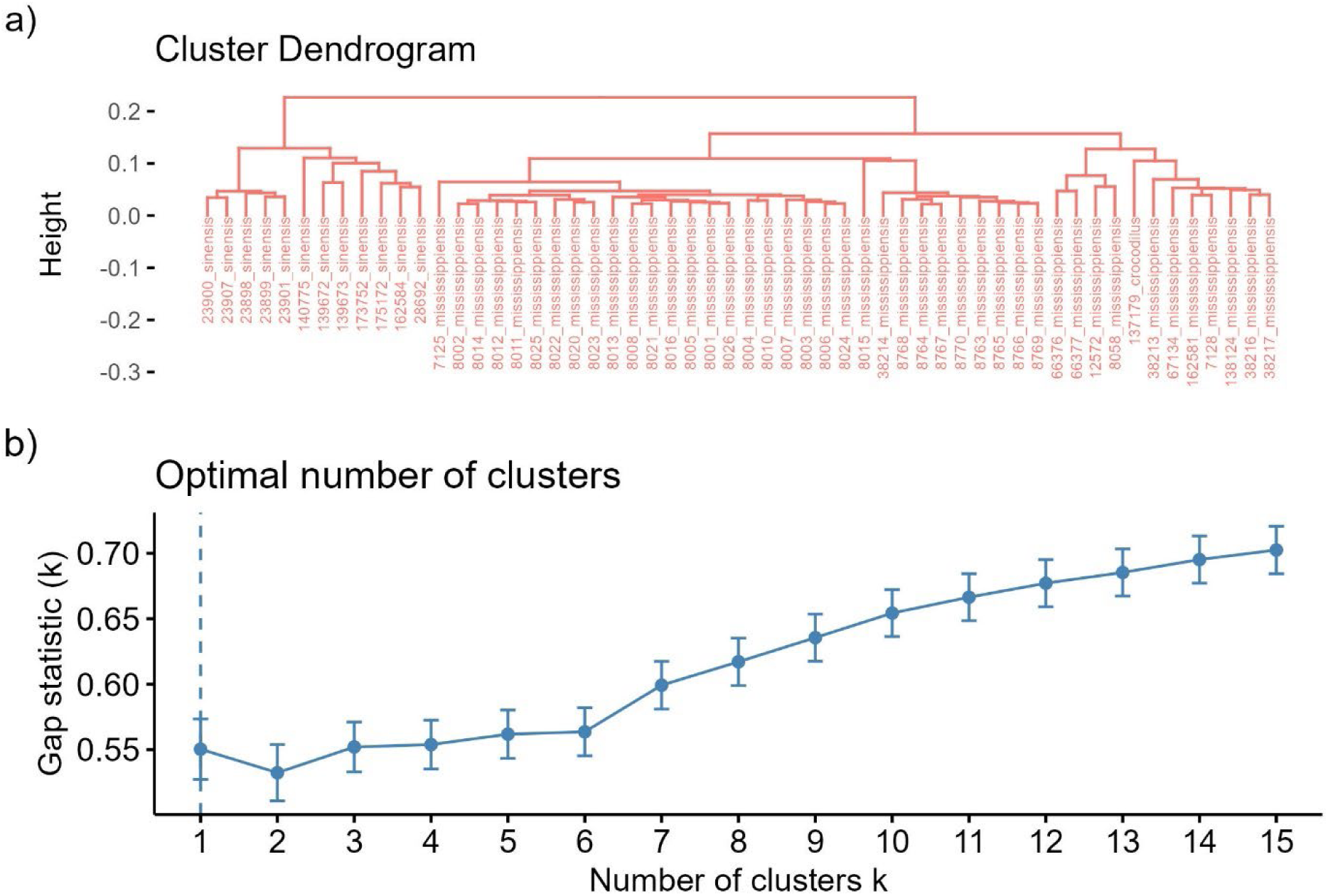
Results of agglomerative hierarchical clustering analysis on PCA scores derived from the geometric morphometrics analysis with PC1 (highly correlated with size), showing a) the cluster dendrogram and b) gap statistic K indicating the optimal number of clusters in the dataset.

Removing all juvenile individuals from the analysis most significantly affects the morphospace by nearly doubling the variation summarized by PC2, presumably because many specimens at one extreme of PC1 are omitted (see Supplemental Appendix II for results of sensitivity analyses). Agglomerative hierarchical clustering suggests a single cluster, interspersing individuals of both species throughout the dendrogram topology. When ontogenetic change is removed from consideration, the clustering analysis does recover a two-cluster scheme corresponding to the two *Alligator* species, with *Caiman crocodilus* placed within *Alligator mississippiensis*. Removing all *Alligator sinensis* juveniles while retaining *Alligator mississippiensis* juveniles does not significantly affect the results of the initial analysis.

Removing all *Alligator mississippiensis* juveniles, while retaining *Alligator sinensis* juveniles, results in a nine-cluster optimal scheme, and with PC1 omitted a two-cluster solution that separates the two *Alligator* species but places *Caiman crocodilus* within *Alligator sinensis*.

Removing adults of *Alligator mississippiensis* results in a 13-cluster arrangement, or a one-cluster arrangement with PC1 omitted, in which *Alligator sinensis* includes *Caiman crocodilus*. Finally, removing adults of *Alligator sinensis* yields a 13-cluster scheme, or a two-cluster scheme without PC1, again placing *Caiman crocodilus* within *Alligator sinensis*.

### Cladistic Analysis of Ontogeny

Survey for ontogenetically and phylogenetically variable characters within *Alligator* yielded 189 discrete morphological characters. Under no outgroup selection does cladistic analysis of ontogeny recover a correct taxonomy. With an artificial embryo as the outgroup OTU, the analysis recovered three most parsimonious trees (MPTs) of length 786 (Fig. 5a). All topologies, and thus the strict consensus, indicate an obvious ‘split’ like that predicted by Zietlow (2020) to suggest the presence of two species or sexual dimorphs of a single species. However, one of these “clades” includes only *Alligator mississippiensis* juveniles, while the other includes individuals of *Alligator sinensis*, *Caiman crocodilus*, and *Alligator mississippiensis* in a highly asymmetrical topology, in which size is correlated with an increasingly crownward position. The tree topology, therefore, clearly shows a mixture of phylogenetic and ontogenetic signal, but does not accurately reflect the known taxonomy of the sample. Importantly, the *Caiman* and the three most mature *Alligator mississippiensis* resolve in a small clade (hereafter referred to as “Clade A”) which is placed such that they appear to be the most mature *Alligator sinensis* individuals. The clade containing an apparent growth series of *Alligator sinensis* thus conveys both misleading phylogenetic signal (viz., it contains three different species) and a misleading ontogenetic signal (viz., inclusion of Clade A suggests ontogenetic changes that do not actually occur in *Alligator sinensis*).

**Figure 5.**
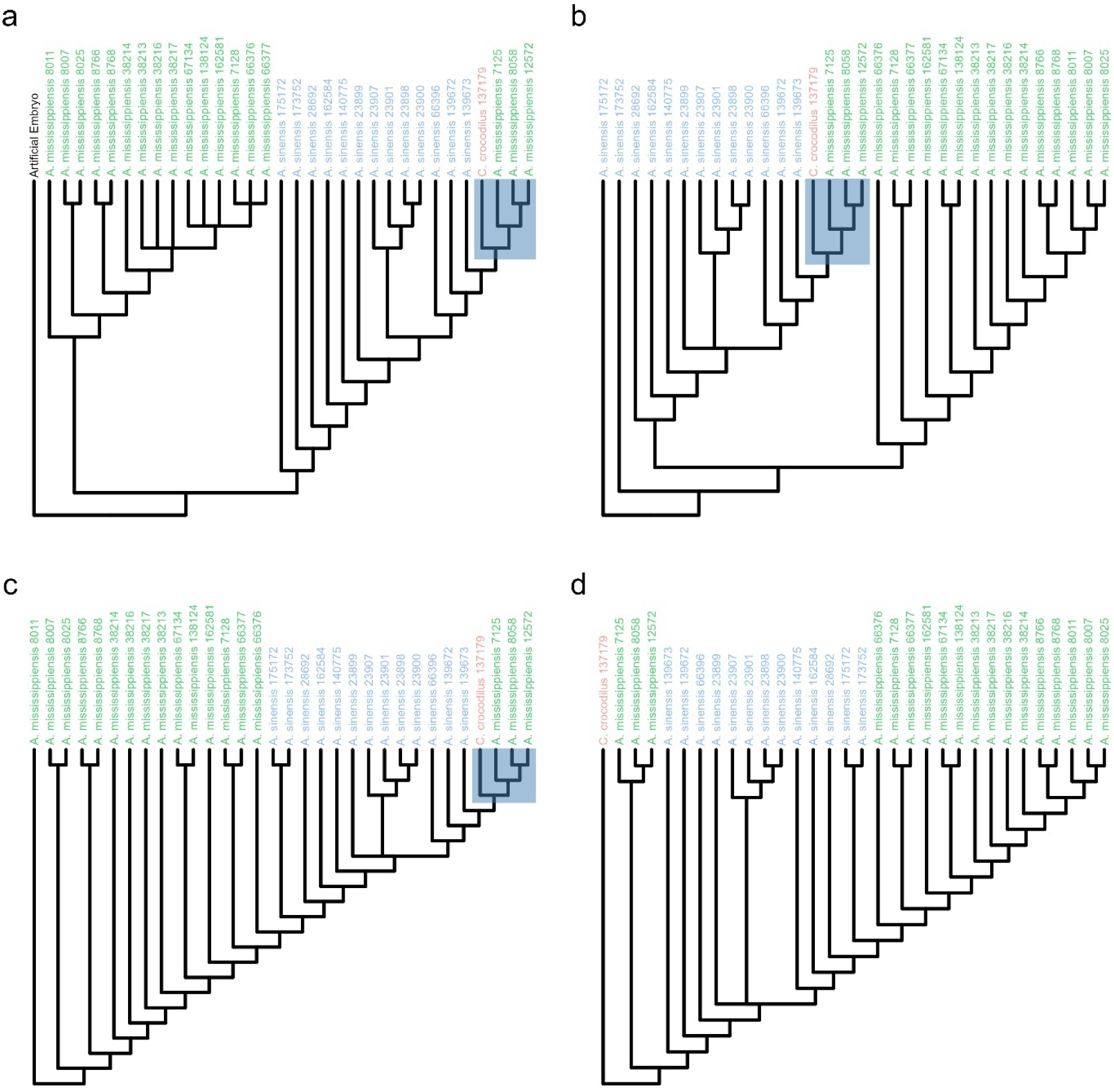
Results of cladistic analysis of ontogeny, using a) an artificial embryo outgroup, b) the juvenile exemplar *Alligator sinensis* as outgroup, c) the juvenile exemplar *Alligator mississippiensis* as outgroup, and d) the *Caiman crocodilus* adult as outgroup. Blue highlights denote Clade A.

When the analysis is polarized based on the juvenile exemplar of *Alligator sinensis*, it yields two MPTs of length 761 (Fig. 5b). These trees differ only in the position of a single individual (AMNH R 23899) of *Alligator sinensis*; results are otherwise comparable to those obtained with an artificial embryo outgroup, notably yielding Clade A as the most mature OTUs at the terminus of an *Alligator sinensis* growth series. AMNH R 23899 is also the only specimen to occupy more than one position when the juvenile exemplar *Alligator mississippiensis* is chosen as outgroup (Fig. 5c). This trial also yields two MPTs of length 761, but these form one highly asymmetrical topology that positions *Alligator sinensis* in the middle of an *Alligator mississippiensis* growth series. The only evidence of a “split” occurs near the tip of the tree, where individuals of *Alligator sinensis* occur on both sides of the split, with one side also including Clade A at its terminus. When the *Caiman crocodilus* individual is used as the outgroup, Clade A obviously cannot form, but again two trees of length 761 form, and AMNH R 23899 is the labile OTU (Fig. 5d). The notable feature of this topology is that successively small specimens are recovered at more “derived” positions; this is unsurprising, because the outgroup OTU is an adult, and therefore this analysis optimizes the mature state as plesiomorphic. Despite the outgroup being the true evolutionary outgroup to the two *Alligator* species, this analysis does not recover any “split” between them, instead placing them within one chimeric growth series.

*Ontogenetically Invariant Characters* The presence of ontogenetically invariant characters was established by applying principal coordinates analysis (PCoA), a distance-based multivariate ordination method that can accommodate discrete characters, to the discrete character matrix. The resultant morphospace (Fig. 6a) clearly separated the two species of *Alligator*, with *Caiman crocodilus* plotting as intermediate between them (Fig. 6a). Both taxa formed linear groups that appear to correspond to parallel ontogenetic trajectories, and Axis 1 shows a logarithmic correlation with increasing centroid size, suggesting that Axis 1 primarily describes an ontogenetic trajectory and that showing a deceleration of character change near the terminus of the sampled series for both species. Despite the visually obvious separation of the two species, hierarchical clustering analysis suggests that the dataset is optimally divided into four clusters (Fig. 7). The same specimens that formed Clade A in the above analysis clustered here within adult *Alligator sinensis*, and removing Axis 1 from the analysis again yields a single cluster (data not shown).

**Figure 6.**
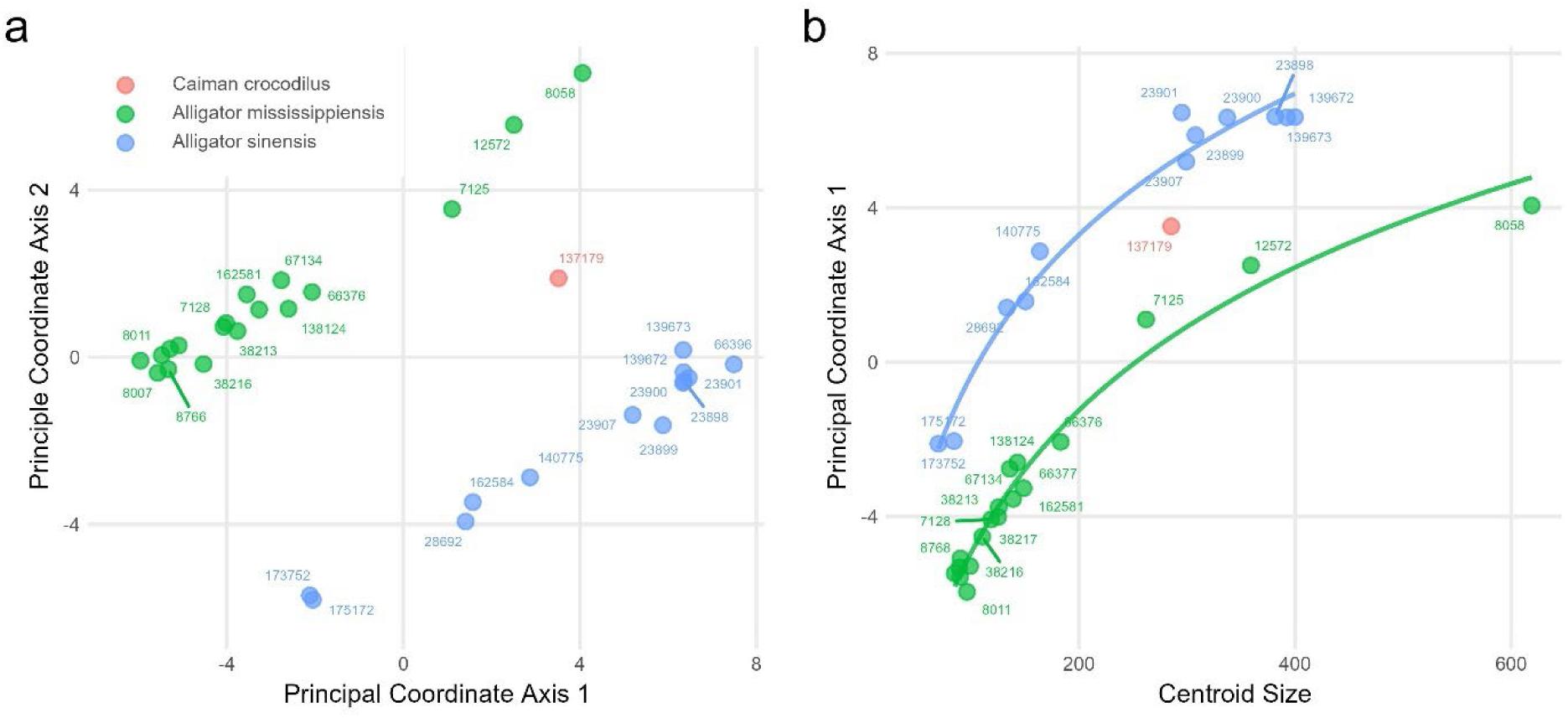
Results of PCoA ordination of discrete character data, including a) principal coordinate axes 1 and 2 and b) relationship between principal coordinate 1 score and centroid size from GM analyses.

**Figure 7.**
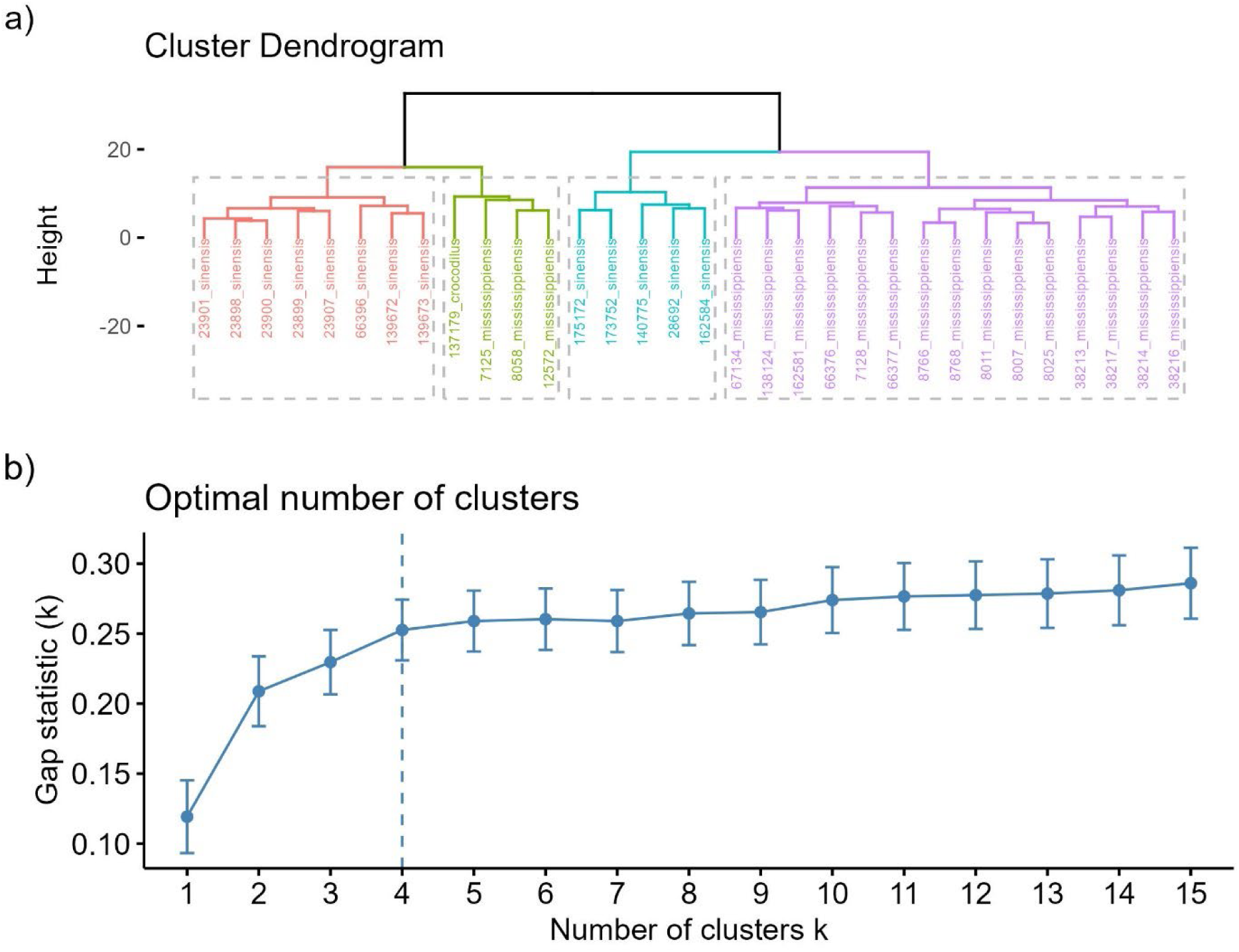
Results of agglomerative hierarchical clustering analysis on PCoA scores derived from the character matrix, showing a) the cluster dendrogram and b) gap statistic K indicating the optimal number of clusters in the dataset.

A manual search for ontogenetically invariant characters that were unique to a species of *Alligator* revealed that the following characters were both taxonomically informative and present throughout the ontogeny of each species, from the youngest to the oldest specimens studied (Fig. 8):

1. Composition of the subnarial foramen (C11). In all *Alligator sinensis*, the subnarial foramen is formed between the premaxilla and the maxilla (C11.0), but in *Alligator mississippiensis* this foramen is formed between the premaxilla and the nasal (which projects a strong subnarial process, which laps over the maxilla anteriorly to exclude it from the subnarial foramen; C11.1).
2. Communication of lateral and caviconchal pneumatic recesses (C33). In *Alligator sinensis*, the lateral pneumatic recess does not anastomose with the caviconchal recess within the maxilla (C33.0); this connection is present in *Alligator mississippiensis* (C33.1)
3. Position of internal foramen for the palatal vasculature (C37). The internal foramen for the palatal vasculature (which leads into a canal that exits on the palatal surface of the maxilla) is positioned in a vertical strut of bone ventral to the lateral recess in *Alligator sinensis* (C37.0), but within the lateral recess in *Alligator mississippiensis* (C37.1).
4. Presence of a descending process of the prefrontal (C58). The prefrontal lacks a descending process in *Alligator mississippiensis* (C58.1) but has one in *Alligator sinensis* (C58.0).
5. Position of lateral postorbital foramen (C84). The foramen opens anteriorly in *Alligator sinensis* (C84.0) but opens laterally in *Alligator mississippiensis* (C85.1).
6. Length of ventral process of ectopterygoid (C111). The ventral process of the ectopterygoid extends about two-thirds of the length of the pterygoid ala in *Alligator mississippiensis* (C111.1), but only extends half the length of the pterygoid ala in *Alligator sinensis* (C111.0).
7. Accessory foramen/notch dorsal to the foramen aereum of the quadrate (C130). This foramen is absent in all *Alligator mississippiensis* (C130.0) and present in *Alligator sinensis* (C130.1). The foramen is not always fully enclosed (and so may be present as a notch), but no indication of one is found in *Alligator mississippiensis*.
8. Ventral palatine foramen. *Alligator mississippiensis* lacks neurovascular foramina on the ventral surface of the palatine medial to the suborbital fenestra (C178.0), but two foramina are present here in *Alligator sinensis* (C178.1).
9. Shape of anterior ramus of vomer. The vomer has an smoothly convex dorsal margin of the anterior ramus in *Alligator mississippiensis* (C179.1), but is concave with a distinct ‘step’ in *Alligator sinensis* (C179.0)
10. Presence of a vomerine recess. A pneumatic recess is present in the vomer of all *Alligator mississippiensis* (C180.1) and is absent in all *Alligator sinensis (*C180.0). Another 10 characters consistently differed between the two species over ontogeny, but showed some degree of polymorphism (e.g., one or both species had at least one individual with the state typical of the other).
11. Length of palatal process of premaxilla (C13). *Alligator mississippiensis* always has a short palatal process of the premaxilla (C13.0), while *Alligator sinensis* has a long palatal process (C13.1) in all but one individual. This individual is a very small juvenile, providing potential evidence that the length of the palatal process establishes after hatching, but given its absence in other juveniles of this taxon I provisionally consider it polymorphic.
12. Length of incisive foramen of premaxilla (C17). *Alligator mississippiensis* generally has a long incisive foramen (C17.1), but two individuals resemble *Alligator sinensis* which has a short incisive foramen (C17.0).
13. Number of maxillary teeth (C23). All individuals of *Alligator sinensis* have either 13 (C23.0) or 14 (C23.1) maxillary teeth, while *Alligator mississippiensis* generally has 15 (C23.2) or 16 (23.3) teeth. A single individual of the latter taxon has 14 (C23.1) teeth on one side, and 15 (C23.2) on the other.
14. Size of lateral fenestra of maxilla (C32). In all *Alligator sinensis*, the lateral fenestra is small (C32.0), while in all but two *Alligator mississippiensis*, the lateral fenestra is large (C32.1).
15. Position of the ostium of the nasolacrimal canal (C45). In all *Alligator sinensis*, the ostium of the nasolacrimal canal is distinctly posterior (C45.1 or 45.2) to the posterior level of the orbital rim of the lacrimal; the ostium is anterior to this level in all but one *Alligator mississippiensis* (C45.0).
16. Presence of a subcristal process of the prefrontal (C57). The subcristal process is absent (C57.0) in *Alligator sinensis*, and present in all but three *Alligator mississippiensis* (C57.1).
17. Presence of an accessory medial process of the ectopterygoid (C112). This process is absent in most *Alligator mississippiensis* (C112.0) but present in all *Alligator sinensis* (C112.1).
18. Composition of foramen for supraorbital branch of trigeminal nerve (C144). This foramen is formed entirely by the laterosphenoid (C144.1) in *Alligator mississippiensis* (variable left-to-right on a single specimen, but otherwise invariant), but is formed between the laterosphenoid and quadrate in *Alligator sinensis* (C144.0).
19. Presence of a vertical ridge on the supraoccipital (C154). All *Alligator mississippiensis* lack a ridge on the supraoccipital (C154.0), and all but one *Alligator sinensis* exhibit one (C154.1).
20. Size of lateral choanal ridges of pterygoid (C168). The lateral choanal ridges are large in most *Alligator sinensis* (C168.0) but small in most *Alligator mississippiensis* (C168.1) such that they do not produce a fossa posterior to them.
21. Presence of neurovascular foramina lateral to the choana (C169). These foramina are absent from the pterygoid of *Alligator sinensis* (C169.0) and present in all but one *Alligator mississippiensis* (C169.1). In sum, 21 discrete characters showed no evidence of ontogenetic variation, and either absent or limited polymorphism within each species. These characters, therefore, do not fit widely accepted conceptual models in which diagnostic apomorphies are acquired throughout postnatal ontogeny, and if included in a phylogenetic analysis would be robust to stemward slippage. However, still other characters did show evidence of ontogenetic variability, but culminated in divergent adult morphologies that make them useful for identifying only more mature individuals (Fig. 9):
22. Palatal diverticulum of caviconchal recess (C34). This diverticulum is always absent (C34.0) in *Alligator sinensis*, and is absent in young *Alligator mississippiensis* but present (C34.1) in mature *Alligator mississippiensis*.
23. Prefrontal recess (C53). This diverticulum is always absent (C53.0) in *Alligator sinensis*, and is absent in the youngest *Alligator mississippiensis* but present (C53.1) in most *Alligator mississippiensis*.
24. Palatine bulla (C177). The palatine bulla is absent in *Alligator mississippiensis* (C177.0) and young *Alligator sinensis*, but manifests in more mature *Alligator sinensis* as an inflated region of the palatines (C177.1).

**Figure 8.**
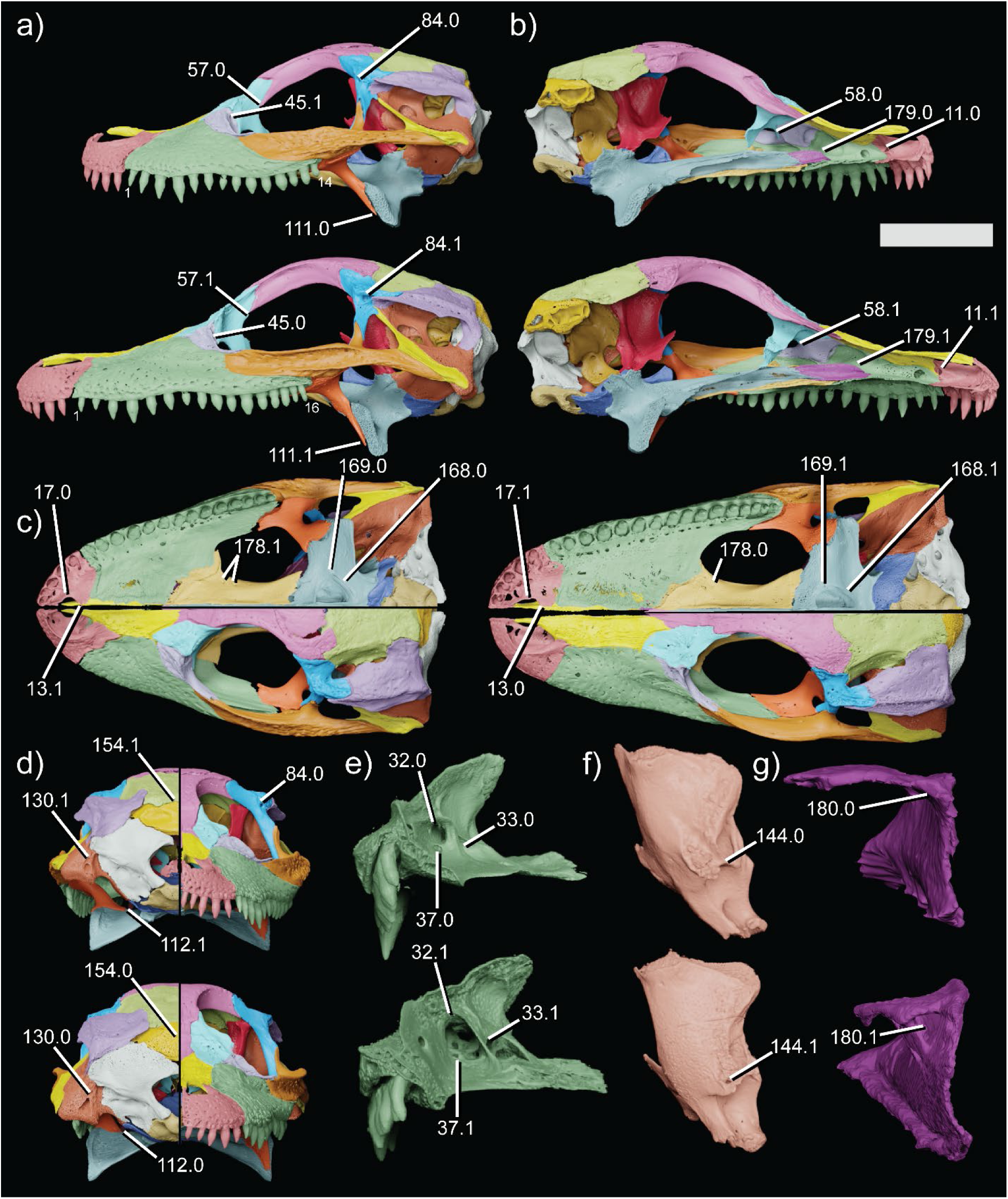
Ontogenetically invariant diagnostic characters and character states showing a) left lateral view, b) medial view, c) ventral (above) and dorsal (below) views, d) posterior (left) and anterior (right) views, e) posterior view of left maxilla, f) posterior view of left laterosphenoid, and g) posterior view of left vomer. For all panels except c), *Alligator sinensis* (AMNH R 175172) is above and *Alligator mississippiensis* (AMNH R 8011) is below; in panel c), *Alligator sinensis* is to the left. Character numbers denote those described in the Discussion and listed in Supplemental Appendix I; states denote states observed in these specimens. Small text numbers denote maxillary tooth count. Scale = 10mm.

**Figure 9.**
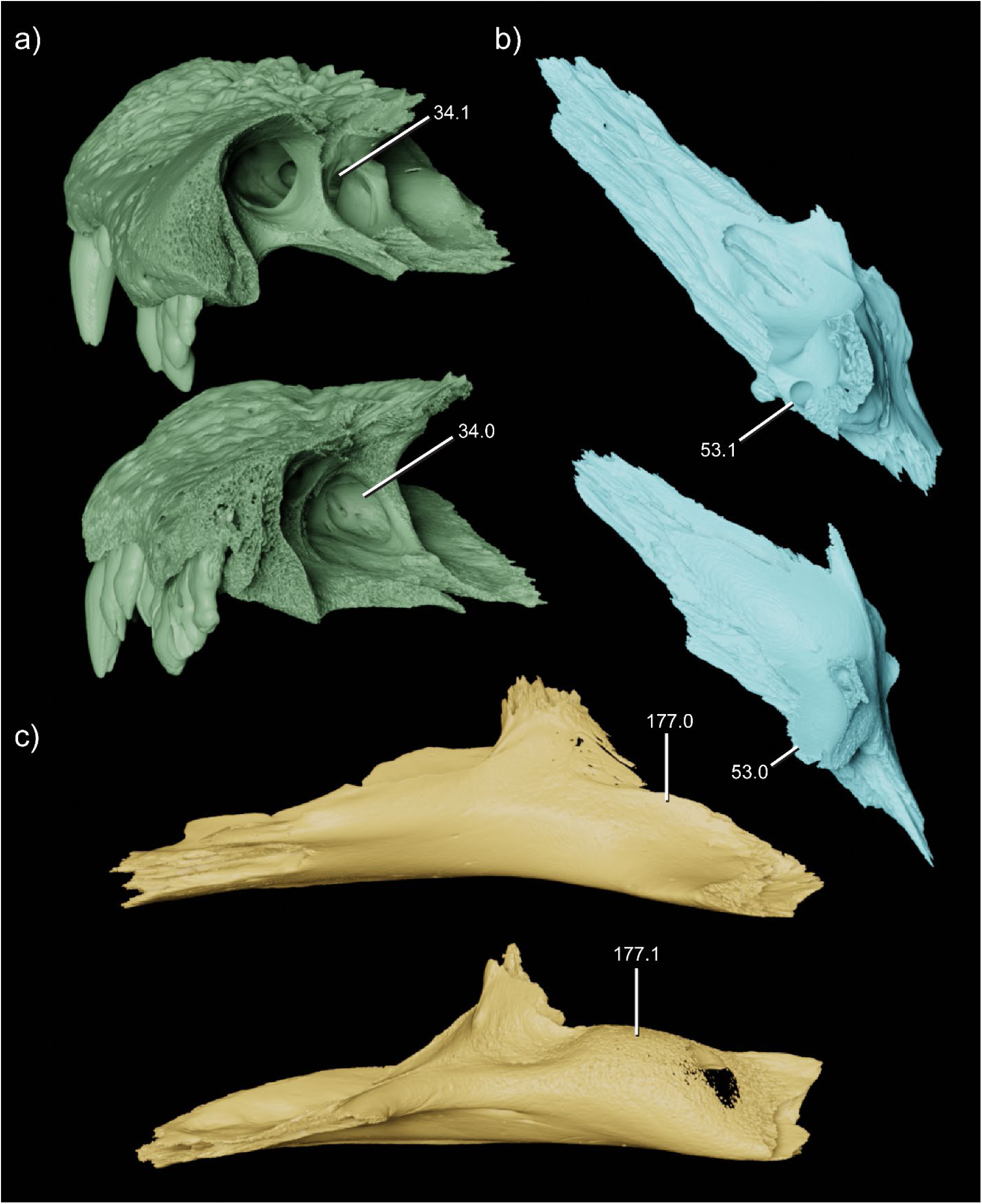
Ontogenetically variable diagnostic characters of *Alligator* species, showing a) left maxilla in oblique posterodorsal view, b) left prefrontal in oblique ventrolateral view, and c) palatine in left lateral view. In all panels, *Alligator mississippiensis* is on top and *Alligator sinensis* is below. Bones are scaled to the same size for clarity.

## Discussion

### Geometric Morphometrics

The results of my geometric morphometrics analyses do not support the use of the method for quantitative tests of taxonomic hypotheses. When the identities of the specimen are known *a priori*, it is trivial to visualize the two ontogenetic trajectories in PC space (Fig. 2), and to recognize that the taxa share occupy fully nonoverlapping shape spaces.

However, agglomerative hierarchical clustering (a method that does not consider *a priori* classifications, only the data themselves) fails to recover these morphospaces as “clusters” (Fig. 3), indicating that the species morphospaces are not statistically emergent (i.e., the data do not form discoverable clusters). With PC1 included, the ontogenetic signal overwhelms any phylogenetic signal; while it is unlikely that the clusters would be interpreted as pertaining to many distinct taxa, due to their obvious differences in size and maturity, it would be impossible to quantitatively determine which individuals belonged to which taxon, and how many taxa were present. Excluding PC1 from the clustering analysis (Fig. 4) effectively factors ontogeny out, but typically overcorrects to a single cluster solution because the two *Alligator* species are, overall, quite similar – they are separated along PC2, but there is more variation within each taxon along that axis than there is range separating them. The only instances in which the clustering algorithm came close to finding the correct taxonomy were those in which either the adult *Alligator sinensis* or the juvenile *Alligator mississippiensis* were excluded from the analyses.

This is unsurprising, because these are the groups that come closest to each other along PC2, which is the only axis that separates the two taxa. Elimination of one group thus inflates the apparent difference between the morphospaces occupied by each *Alligator* species. In sum, geometric morphometrics fails to empirically recover the known taxonomy of the trial dataset except in trials excluding a significant amount of the data.

Furthermore, in all iterations of the GM analysis, the *Caiman crocodilus* individual clustered within a species of *Alligator*. The *Alligator* ontogenetic series dominate the variation in the sample, and thus also dominate the signal inherent in the PC space. In a dataset like this, the singleton taxon would need to be radically different from all other specimens to fall clearly outside the morphospaces of the other taxa – but in this case, it seems unlikely that workers would mistakenly include it in the ontogenetic series in the first place. Despite the numerous character differences between *Alligator* and *Caiman*, nothing in the morphospace plots would suggest that the *Caiman* individual was not an alligator. Even large samples of both *Alligator* species were not statistically emergent, and this problem is only compounded when acknowledging the possibility that some specimens may pertain to an unexpected third taxon.

We should expect that extinct ecosystems hosted multiple closely related species, just as modern ecosystems do today even at high trophic levels, and therefore this phenomenon is troubling. It is difficult to envision how a GM analysis of a fossil dataset could adequately account for the possibility that unsuspected new taxa are included, especially if only one or a paucity of the individuals sampled pertain to that new taxon. Given that the two *Alligator* species do clearly separate in PC space, it may appear that visual examination of morphospace plots is the most effective way to interpret the results of GM analyses. However, this practice is not repeatable or empirical and does little more than add a quantitative veneer to a fundamentally qualitative taxonomic assessment. If mathematics is to guide taxonomy, the data must speak for themselves.

The failure of the data to do so in the present analysis is concerning. While the two *Alligator* species are overall quite similar, they are morphologically distinguishable and have never been suggested to be synonymous. The failure of morphometric methods to recover obviously distinct groups in morphospace suggests that geometric morphometrics cannot effectively discriminate between anatomically similar species, especially when faced with ontogenetic and polymorphic variation within both. Geometric morphometrics is, then, highly susceptible to type II error (false negatives, or failures to recognize species that are present in a dataset), and does not offer a practical solution to the ontogeny problem. Furthermore, sampling biases also introduce significant potential for type I error (false positives, or erroneous recognition of new taxa). No morphospace is free of gaps – individuals within populations vary independent of growth stage, sometimes significantly. If intermediate individuals are unknown, it is very possible that individuals from two extremes of a taxon’s morphospace would be mistaken for members of separate species, with the presence of a morphological gap between them taken as evidence in favor of separation. For example, several recent studies have suggested that one specimen of *Velociraptor mongoliensis* (Mongolian Academy of Sciences [MPC-D] 100/982) pertains to a new species due to an apparent gap in morphospace between it and other *Velociraptor* specimens (Powers et al. 2020, Powers et al. 2022). While this is certainly plausible, with only four specimens included in the analysis, it remains possible that we are simply missing morphologically intermediate *Velociraptor mongoliensis* individuals, and that this “gap” is a sampling artefact attributable solely to the low sample size. While geometric morphometrics is a powerful tool for addressing many questions in evolutionary biology, and may offer some valuable perspective on which specimens are truly the most dissimilar, I do not consider it methodologically or epistemologically sufficient for testing taxonomic hypotheses for the reasons enumerated above.

### Cladistic Analysis of Ontogeny

All four of my attempts at cladistic analysis of ontogeny fail to yield consensus topologies that indicate the correct phylogenetic and ontogenetic status of the specimens in my dataset (Fig. 5). None of my analyses corroborated the prediction by Zietlow (2020) that cladistic analysis of ontogeny would recover a clean divergence between the sampled species. While two outgroup protocols did recover strict consensus trees with basal divergences, these divergences did not sort OTUs along clearly phylogenetic or ontogenetic lines. Instead, one clade comprised only juvenile *Alligator mississippiensis*, and the other an apparent growth series of *Alligator sinensis*, with *Caiman crocodilus* and adult *Alligator mississippiensis* placed as its most mature members (forming Clade A). Thus, these topologies have the dubious distinction of conveying both misleading phylogenetic signal and misleading ontogenetic signal. In a paleontological study where species identity is unknown (or the question of focus), it would be impossible to determine how many species are represented within the ontogram based on its topology alone. The two other protocols found a highly asymmetrical strict consensus topology without a basal divergence, and in each *Alligator sinensis* is placed within the *Alligator mississippiensis* growth series. Zietlow (2020) suggested that such a topology could also be consistent with multiple species, if terminal branches showed “homologous sets of intraspecific variation”. This tree would, presumably, represent one in which the ontogenetic signal is dominant (perhaps due to similar ontogenetic sequences in both taxa), forcing traits specific to one species to optimize as convergences at different points along the ontogenetic trajectory. However, this pattern does not manifest in the present analysis.

Rather, putative ontogenetic characters (which are, actually, the phylogenetic characters that distinguish the two species) optimize with two state changes – one from the condition in *Alligator mississippiensis* to that in *Alligator sinensis*, followed by a reversal to the plesiomorphic state in adults of the former species. This is important to note, because apparent reversals of large suites of ontogenetic characters have been found in prior studies employing cladistic ontogeny – most notably, Carr (2020) reported many reversals to putatively juvenile states among adult *Tyrannosaurus rex*. My results suggest that these patterns may warrant further scrutiny. In summary, cladistic analysis of ontogeny did not recover a correct taxonomy, nor did it accurately describe the sequence of character changes that occur during ontogeny in either taxon.

These analyses fail to validate the proposal that cladistic ontogeny can effectively sort ontogenetic and phylogenetic variation. Beyond my own empirical test, however, it is worth exploring the method at a deeper level due to its apparent success when applied to fossil datasets. It is possible that my trial dataset differs in some fundamental way from those assembled for fossil datasets. As in exemplar uses of cladistic ontogeny such as those of Carr (2020) and Zietlow (2020), I compared specimens over a range of body sizes and presumed ontogenetic stages and strove to codify every morphological character difference that was anatomically unambiguous and repeatably scoreable. My analytical protocol, however, does differ from these in three key respects. First, my dataset has a relative paucity of missing data compared to paleontological datasets, but having more data should, if anything, improve my analyses rather than hindering them. Second, Zietlow (2020) did not treat multistate characters as ordered. This is clearly problematic when considering ontogenetic characters, which we may expect to change gradually and unidirectionally – for example, several in Zietlow’s (2020) dataset describe the absolute size of certain elements, and treating these characters as unordered does not penalize ontogram topologies that infer growth stages where bones grow quite suddenly without passing through intermediate character states, nor topologies in which a bone may actually *shrink* before growing again. Carr (2020) does, however, use ordered characters in his analysis, so my use of ordered characters is not at odds with all prior implementations of this method. More problematically, Carr (2020) reports an analytical protocol in which taxa were iteratively added to the ontogram search, to determine which specimens induced the recovery of multiple MPTs; these specimens were considered wildcards and removed from the analysis *a priori*. Treatment of wildcard specimens in phylogenetic analyses is a topic worthy of discussion in its own right; some OTUs may be so incomplete that they result in a net loss of information from the tree search, but this is an extreme case, and I concur with perspectives generally favoring inclusion of labile OTUs in the tree search, but removing them *a posteriori* to report reduced strict consensus tree topologies. In any case, identifying any OTU with multiple most parsimonious positions as a “wildcard” seems like an exaggeration, and the result is an analysis that purposefully eliminates alternative most parsimonious topologies, and does not correctly communicate what ambiguity may exist within the dataset. My use of conventional tree search approaches and consensus trees, therefore, may not precisely mirror Carr (2020), but may be seen as indicating the kinds of alternative topologies that his analytical protocol eliminates from consideration.

A further consideration is the possibility that *Alligator mississippiensis* and *Alligator sinensis* are more dissimilar from one another than any undetected fossil species would be, and thus that *Alligator* is an unsuitable model system for assessing the efficacy of cladistic ontogeny. It is true that the two taxa differ in many characters, including many that are ontogenetically invariant and others that appear during growth (see below). However, the presence of a “split” manifesting in an ontogram containing multiple taxa, as predicted by Zietlow (2020), would logically be more likely to occur in situations where the two taxa are differentiated by a large number of characters. The two species of *Alligator* diverged in the Eocene, and have had about 50 million years of time to acquire new autapomorphies (Wu 2003). If cladistic ontogeny cannot reliably detect the difference between two species that diverged shortly after the extinction of the non-avian dinosaurs, it is difficult to envision how it could possibly be a reliable means of delimiting shallowly diverged, possibly sympatric species that are known only from an incomplete fossil record.

Beyond my empirical failure to validate the method, there are several epistemological considerations that make cladistic analysis of ontogeny an unsuitable method for testing taxonomic hypotheses The use of an “artificial embryo” as the outgroup OTU is fraught with difficulty. The artificial embryo OTU is, in practice, a collation of the hypothesized immature states for each character, based upon the least mature specimens for which each character is preserved. This runs the obvious risk of inferring an “embryonic” condition that is actually informed by individuals that have already progressed significantly far along their ontogenetic trajectory. The most extreme case was a recent cladistic ontogeny analysis of growth among alioramin tyrannosaurids, in which the approximately nine-year-old holotype specimen of *Alioramus altai* was used as the sole source of “immature” characters, and was scored identically to the artificial embryo (Foster et al. 2022). Determination of which specimens are “least mature” is typically made based on size, rather than osteohistological data – and indeed, several analyses include absolute size among their ontogenetic characters (Ezcurra and Butler 2015, Carr et al. 2017, Carr 2020, Zietlow 2020). This process results in the outgroup OTU representing the character states observed in the smallest specimens to display each character, meaning that it is a composite that may not represent the true combination of characters at any stage of development, and forcing the analysis to interpret these states as “immature” and enforcing the ontogenetic trajectory that had previously been inferred based on size alone. Therefore, despite claims that cladistic ontogeny is advantageous in offering a size-independent means of maturity estimation (Brochu 1996, Carr 2020, Zietlow 2020), these analyses are not size-independent in practice. It is perhaps unsurprising, then, that published analyses employing cladistic ontogeny typically find a robust correlation between size and maturity. If the ontogenetic character matrix includes multiple species, the artificial embryo will represent a combination of ontogenetic and phylogenetic information, making it very difficult to interpret the resultant ontogram topology. Any node may be supported by a mixture of ontogenetic and phylogenetic characters, making it difficult to determine which “clades” represent growth stages, which represent species, and which represent some combination of both (e.g., a valid species and immature members of a larger sister taxon). A similar problem has emerged in invertebrate paleontology through the use of “semaphoront coding” (Lamsdell and Selden 2013), which scores immature and adult members of species as separate OTUs, and thereby generates “phylogenetic trees” that do not actually contain a purely phylogenetic signal (Sharma et al. 2017). Both semaphoront coding and cladistic ontogeny, then, produce “tree-shaped objects” (Wheeler and Pickett 2007) from mixed data sources that cannot be interpreted readily. Even if it were simple to distinguish between these data sources on an ontogram topology, published expectations for how different scenarios should manifest in an ontogram lack explicit criteria and include topologies both with and without deep divergences (Zietlow 2020), essentially meaning that almost any conceivable topology could be interpreted in different ways by different workers. Most fundamentally, progressive ontogenetic change occurs within individual organisms and, as such, do not meet the assumption of hierarchical organization that is inherent to Hennig’s model of phylogenetic systematics (Sharma et al. 2017), despite claims to the contrary (Brochu 1996, Zietlow 2020). “Adult” is not a subgroup of “juvenile”, in the way that species are subgroups of larger clades. In summary, a host of philosophical and practical challenges make cladistic ontogeny unsuitable for testing taxonomic hypotheses in the fossil record, and I suggest that the results of prior studies using the method be interpreted with healthy skepticism until they can be confirmed through other, more robust analytical protocols.

### Ontogenetically Invariant Characters

Multivariate ordination of my discrete character matrix recovers two obvious linear clusters (Fig. 6a), which correspond to the two species of *Alligator*. Axis 1 correlates closely with centroid size (Fig. 6b) and can be interpreted as the primary ontogenetic trajectory; the two species have approximately parallel trajectories, suggesting broadly similar ontogenetic trends (as we may expect among closely related taxa). Axis 2, then, can be interpreted as the primary phylogenetic axis. Notably, the spectacled caiman individual plots between the two species of *Alligator*, and is approximately equally spaced from both along Axis 2 and in the region of similarly-sized *Alligator* individuals of both species along Axis 1. Despite the visually obvious separation of these ontogenetic trajectories, hierarchical clustering analysis does not elucidate a correct taxonomy (Fig. 7); instead, the dendrogram clearly shows a primarily ontogenetic signal, in which juveniles diverge from adults, after which each growth stage separates along taxonomic lines (excepting *Caiman crocodilus*, which clusters with *Alligator mississippiensis*). Removing PCoA Axis 1 from the clustering analysis results in a single cluster scenario, so removing the influence of ontogenetic variation does not clarify the taxonomic composition of the sample.

In sum, 23 of 189 characters (∼13%) were highly phylogenetically significant, and of these 20 showed no signs of ontogenetic variation. Thus, we can affirm that closely related living species can and do possess ontogenetically invariant characters that allow confident determination of their identity from apomorphic evidence alone. Therefore, the repeated failure of existing multivariate methods might be of less concern than it would outwardly appear. While ontogenetic variation is predominant in my dataset, and some diagnostic characters do occur in adults only, it is also clear that individuals are born with characters that do not change during ontogeny. Importantly, these characters need not be simply general characters shared by every species in their clade, but can instead be highly diagnostic traits that distinguish close relatives and obviate the need for multivariate statistical approaches. Even very young *Alligator* display a suite of ontogenetically invariant characters that show no evidence of polymorphism, and while the two species do become increasingly distinct during growth this is encouraging evidence that fossils of young animals may similarly already possess important diagnostic characters. The question now becomes whether or not it is possible to predict which characters are ontogenetically invariant in extinct taxa – and therefore, which characters are “good” characters for testing taxonomic hypotheses.

Reviewing the above list of characters, it should be immediately evident that ontogenetically invariant characters are overwhelmingly associated with neurovascular foramina, pneumatic recesses, and the sutural topology of the skull. Characters associated with muscle scars, the shapes of bone margins, ornamental features, and overall proportions show far less tendency to be ontogenetically invariant, and are in fact among the most ontogenetically variable character types in my dataset. The asymmetric distribution of ontogenetically invariant characters suggests that their invariance is not some evolutionary quirk unique to *Alligator* or a similarly exclusive clade; rather, I propose that these characters are invariant due to broadly-conserved aspects of the vertebrate developmental program, and are thus generally applicable to a range of study systems.

Despite the common lay perspective that bones are the “scaffold” or “foundation” for the soft tissues of the body, bones appear fairly late in development, and their patterning and ultimate morphology are shaped by the soft tissues that precede them. Of the most informative character types identified above, two (cranial foramina and pneumatic recesses) relate to soft tissue systems that are laid down prior to the onset of cranial skeletogenesis. In human embryology, cranial nerve development occurs quite early – most cranial nerves appear between developmental days 26-38 (Carnegie Stages 12-15), and by day 53 (end of Carnegie Stage 20) all nerves are both present and have reached their target organs (Smit et al. 2022). The cranial arteries also appear early in development (Takahashi et al. 2012), and assume an adult branching pattern by approximately ∼52 days of development (Paget 1948). The onset of cranial bone development is far later – for example, in humans the sphenoid complex begins to form at 16 weeks (∼112 days) of development (Mano et al. 2021). Cranial foramina form not as a result of developing blood vessels and nerves ‘tunneling’ through the bone to reach their targets, but because the developing bones envelop the pre-existing neurovasculature as they condense from mesenchyme. The neurovascular structures are not passive participants in this process – in chicken embryos, chondrogenesis of the occipital bones is interrupted in the immediate vicinity of primordial jugular, carotid, and hypoglossal foramina, which presumably protects the contents of these foramina from being stenosed or severed by encroachment of the surrounding bone (Akbareian et al. 2015). It has been suggested that the nerves and blood vessels themselves actually *initiate* the slowdown in chondrogenesis that leads to the development of a foramen to transmit them (Akbareian et al. 2015). Cranial nerve foramina form during embryonic development, in response to the position of nerves and blood vessels that are laid down very early in development, long prior to mesenchymal condensation and the later onset of chondro- or osteogenesis. Furthermore, dermal bone osteogenesis follows expansion of capillary networks within mesenchymal condensations, suggesting that these capillaries are a source of signals that promote ossification and that alterations to the vascular pattern can induce changes in skeletal morphology (Percival and Richtsmeier 2013).

Pneumatic recesses in cranial bones are enigmatic structures of unclear function that show a high degree of variation. The epithelial sinuses housed in these recesses have a documented tendency to expand progressively during life, which has been noted in systems as diverse as *Alligator* (Witmer 1995, Dufeau and Witmer 2015) and humans (Scuderi et al. 1993). The growth of cranial sinuses (and therefore, the recesses they leave in the cranial skeleton) is often described as ‘opportunistic’, given the apparent tendency of these sinuses to expand as far as they can. Witmer (1997) proposed the ‘epithelial hypothesis’ of sinus growth, positing that sinus growth is driven by an intrinsic capacity of the epithelia to invade and excavate bone, and is constrained only by the biomechanical integrity of the skull. This hypothesis explains much of the variability that is obvious in cranial sinuses, such as asymmetry and development of pathologic fistulae. Given the abundant observations of how ontogenetically and individually variable sinuses are, I and many of my colleagues have historically shied away from their use as taxonomic characters. And yet, my data for *Alligator* suggest that the basic pattern of pneumatic invasion of the cranial skeleton is highly diagnostic. While it is true that both *Alligator* species show progressive expansion of the cranial sinuses, the simple presence or absence of a pneumatic recess was consistent, as were large-scale patterns of pneumatization (such as confluence of the lateral and caviconchal recesses of the maxilla). Perhaps unsurprisingly, the precursors of cranial sinuses also appear early in development as nascent outpocketings of the respiratory cavity, which are later enclosed by bone during condensation (Scuderi et al. 1993, Witmer 1995). Unlike cranial nerves and blood vessels, not all cranial sinuses are present *in embryo*, but those that are not do appear quite early in postnatal ontogeny. In *Alligator mississippiensis*, all sinuses are present within a few months of hatching (Witmer 1995), and humans pneumatize the frontal sinus (the last sinus to form) by the second year of life (Scuderi et al. 1993). The consistency of which bones are pneumatized within species suggests that there is greater regulation of sinus development than proposed by the ‘epithelial hypothesis’ – while I do not doubt that sinuses do grow progressively and opportunistically, there seems to be additional governance that controls where these sinuses are able to *start* growing, and their ultimate morphology (e.g., why the palatine inflates into a bulla in *Alligator sinensis* but not *Alligator mississippiensis*). The similarity in cranial anatomy between the two *Alligator* species precludes explanation of their divergent pneumatization patterns as reflecting differences in mechanical constraints on their skulls; I consider it more likely that there is a fundamental molecular regulatory program that dictates where the sinuses are “allowed” expand. In humans as well, despite a high degree of variation in sinus size and extent, the basic pattern of pneumatization is highly constrained – humans develop sinuses in the maxilla, ethmoid, sphenoid, and frontal. The human palatines and vomers, despite being in direct apposition with the respiratory tract, do not pneumatize – and the presence of a pneumatic recess in the extremely thin vomer of *Alligator* suggests that this is not simply a consequence of their delicate nature. Further research is required to resolve the dichotomy between sinus distribution (which appears tightly regulated) and sinus expansion (which does appear to be fundamentally opportunistic and occasionally leads to pathology). One further point of relevance is that there is no evidence of any cranial sinus being *lost* during ontogeny by overgrowth of bone to constrict its ostium. Dufeau and Witmer (2015) suggest that the laterosphenoid recess is lost during crocodilian ontogeny, but the pneumatic space within the bone persists in all specimens studied herein; what is lost is the ostium of this recess, which passes through the sutural surface of the laterosphenoid and prootic, and closes as their joint surface becomes increasingly interdigitate through ontogeny. Putative juvenile *Tyrannosaurus rex* individuals have a pneumatic recess in the quadratojugal that is absent in adults (Witmer and Ridgely 2010, Carr 2020), but this pattern is unknown in other species; another large theropod, *Allosaurus*, appears to have had a the same complement of pneumatic sinuses throughout posthatching ontogeny (Rauhut and Fechner 2005). In sum, we can conclude that the topological distribution of pneumatic recesses is highly species-specific, and expect that only very young members of a species may lack pneumatic recesses present in the adults. The size and morphology of these recesses may readily change during growth, but growth appears to be unidirectional – we should not expect ontogenetic transformations in which pneumatic recesses shrink or close.

My data on *Alligator* also suggest that sutural topology of the cranial skeleton is ontogenetically invariant and shows little polymorphism. By this, I mean not the form of the joint surfaces between bones (which show an obvious ontogenetic trend towards interdigitation), but in *how* the bones contact – which bones form joints with which others, the discrete processes, rami, or laminae of each bone, and how the bones relate to cranial openings. In the present dataset, presence of a subnarial process of the nasal is itself sufficient to identify a specimen as *Alligator mississippiensis*, and presence of a descending process of the prefrontal is likewise sufficient to identify a specimen as *Alligator sinensis*. The ossification of skull bones follows patterns laid down very early in development, which are dictated by gene expression in the embryonic brain and both sensory and ectodermal epithelia (Noden 1988, Hanken and Thorogood 1993, Hu et al. 2003, Wada et al. 2005, Brugmann et al. 2007, Bhullar et al. 2015).

While current research has so far allowed confident identification of the mechanisms responsible for mostly “coarse” morphological changes, such as the fusion of the avian premaxillae into a beak (Bhullar et al. 2015), there is no reason to think “finer” details such as the presence or absence of a bony process, or the position of a bony contact relative to some landmark, are not governed by similar signaling cascades. Indeed, embryos of different galloanseran bird species show distinct developmental trajectories from the first appearance of facial landmarks (Smith et al. 2015), leading the authors to propose that morphological differences among adults of closely related species can and do arise from modification to the early stages of development. The development of the vertebrate skull is complex, and represents the influence of multiple soft-tissue systems in addition to pleiotropic interactions between skull bones themselves (Percival et al. 2018), and while we certainly lack a general predictive model for skull morphology available evidence makes it clear that the topological architecture of the skull is governed by early developmental events, and that we therefore should not expect architectural changes to the skull during postnatal ontogeny.

I also found no evidence for ontogenetic loss of maxillary tooth positions in modern crocodylians. Expanded data collection from alligatoroid specimens in the AMNH Herpetology collection suggest that variation, if present, is exclusively on the order of ±1 positions, and shows no consistent relationship with body size (Fig. 10) – therefore, I interpret this character as polymorphic, but not ontogenetic. Indeed, Brown et al. (2015) found a similar pattern in both crocodilians and the Komodo dragon, suggesting that macropredatory reptiles may be generally characterized by ontogenetically stable maxillary tooth counts; however, the presence of an ontogenetic *increase* in tooth count common to many lizards, basal archosauriforms, basal sauropodomorphs, ornithischian, and perhaps basal theropod dinosaurs (Rauhut and Fechner 2005, Brown et al. 2015, Ezcurra and Butler 2015, Chapelle et al. 2019) indicates that ontogenetically invariant tooth counts are not a general feature of tetrapods. Maxillary tooth count is interesting in that it provides direct rebuttal of proposed tooth loss in theropod ontogeny along two complementary lines of evidence. First, Carr (1999) partially justified a loss of several maxillary tooth positions between juvenile *T. rex*/*Nanotyrannus* and adult *T. rex* by analogy to crocodylians, citing Mook (1921), Wermuth (1953), and Iordansky (1973). I am unable to read German, so I cannot verify what Wermuth (1953) actually wrote; however, Carr (1999) did not accurately reflect what Mook (1921) and Iordansky (1973) described in their papers. Mook (1921) identified the number of maxillary teeth *below the orbit* as undergoing an ontogenetic reduction in crocodylians; this observation is correct, but does not involve loss of maxillary teeth, but simply positive allometry of the snout that draws the entire toothrow forward, causing fewer teeth to lie ventral to the orbit. Iordansky (1973) stated that the 2^nd^ premaxillary tooth of some species is lost by juveniles, but does not describe maxillary tooth count as decreasing during ontogeny. The loss of a premaxillary tooth during life is common in crocodylians, but occurs due to pathology (overgrowth of a dentary tooth can obliterate the tooth position and leave a diastema in its place), rather than as a normal feature of the posthatching developmental program. In any case, Carr (1999) was mistaken in citing these papers as describing the ontogenetic loss of maxillary tooth positions, and as described above this pattern is not documented in either my data or other more recent studies of crocodylian anatomy. Second, this character serves as a cautionary tale. While compiling my data matrix, I did think that ontogenetic maxillary tooth loss would prove to be a trend in crocodylians, because both of my juvenile exemplars had one more maxillary tooth than their adult counterparts. My full data matrix revealed that this was entirely a chance occurrence, driven by a low sample size and rectified with the addition of more data. It seems that Carr (1999) may have been affected by a similar issue. Carr (1999) proposed an ontogenetic reduction from 15 maxillary teeth in “stage 1” juveniles to 13 in “stage 3” adults of the tyrannosaurid *Gorgosaurus libratus*, based on a sample of only 10 specimens and without any quantitative analysis. Currie (2003a) found no statistical evidence for ontogenetic reduction in tooth count in *Gorgosaurus*, and recently described individuals smaller than those studied by Carr (1999) both have 14 maxillary teeth (Voris et al. 2022), which is the most common dental count for this genus. Furthermore, the youngest known individual of *Tarbosaurus bataar* has only 13 maxillary teeth, like most *Tarbosaurus* adults (Tsuihiji et al. 2011). Carr et al. (2017) proposed that another tyrannosaurid, *Daspletosaurus torosus*, gained and then lost teeth as it grew, but this observation is based upon a longstanding misidentification of a juvenile specimen, which is more convincingly interpreted as a young *Gorgosaurus libratus* (Voris et al. 2019). Without this specimen, all *Daspletosaurus torosus* individuals have between 14 and 16 maxillary tooth positions, and thus are consistent with the range of variation seen in crocodylians and other tyrannosaurs. Therefore, the available evidence favors the interpretation that tyrannosaurids, like modern crocodylians, did not lose maxillary tooth positions during growth, and instead showed limited variation about a modal tooth count that was particular to each species. Further research is required to determine where transitions between ontogenetic increase vs. stability of tooth counts occur, and what anatomical, developmental, or ecological factors correlate with these alternative patterns.

**Figure 10.**
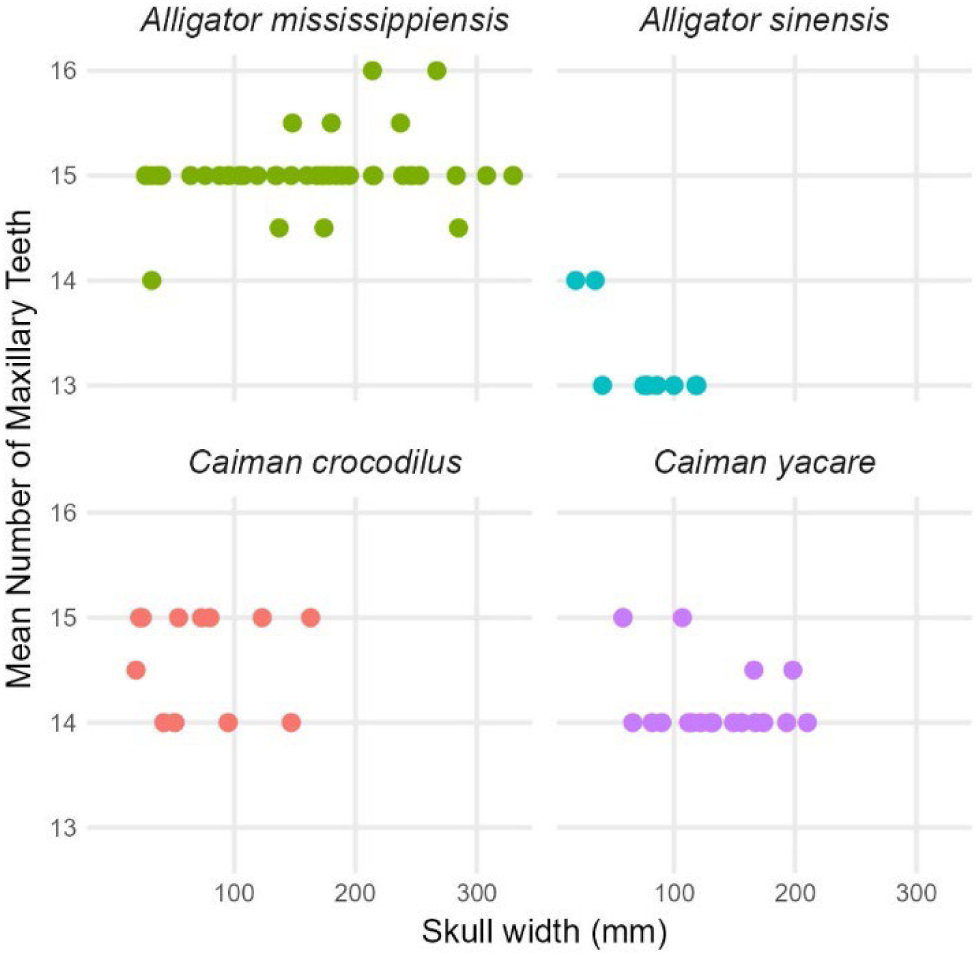
Maxillary tooth count in select extant alligatoroids, illustrating typical variation of ±1 maxillary tooth positions. Tooth counts are averaged from left to right sides, sometimes yielding fractional counts in individuals displaying asymmetrical variation.

The ontogenetic invariance of dental count is of interest beyond Tyrannosauridae, however, for two main reasons. First, tooth count is a widely applicable and presumably homologous character among most tetrapods, allowing a rare example of directly comparable character states among a wide array of taxa. Secondly, the generation of teeth is yet another process mediated by epithelial-mesenchymal interaction during development. The first teeth in *Alligator* appear at approximately the same time as the cranial nerves and arteries, though the ultimate patterning of the tooth row and determination of the final number of functional tooth positions continues to hatching (Westergaard and Ferguson 1990). While the first embryonic tooth generation is formed from cells derived from odontogenic placodes, an epithelial cell population termed the dental lamina replaces it and persists as a source of odontogenic stem cells throughout life (Tsai et al. 2016). The persistence of a dental lamina permits *Alligator* (and toothed amniotes generally) to undergo continuous tooth replacement (polyphyodonty), and this cell population plays a key role in patterning the toothrow, such as determining the number of tooth positions (Westergaard and Ferguson 1990). Given the above-discussed tendency of epithelial tissues to maintain and create space for themselves, it seems plausible to hypothesize that the persistence of a dental lamina precludes ontogenetic loss of tooth positions. Only one reptile, the early-diverging theropod *Limusaurus inextricabilis*, is documented to lose tooth positions ontogenetically, and this appears to result from a complete cessation of tooth replacement that is consistent with degeneration of the dental lamina (Wang et al. 2017).

However, in *Limusaurus* void spaces persist in the maxilla and dentary in the former positions of the dental alveoli, suggesting that the dental lamina alternative may persist but become inactive. In any case, we can observe that like the other character types described above, tooth patterning is a process that occurs early in development (long prior to the condensation of the eventual tooth-bearing bones) and shows no ontogenetic variation in my model system. This result may be broadly applicable, given that extant macropredatory reptiles also show no sign of ontogenetically variable tooth counts, but the phylogenetic distribution of ontogenetically stable dental counts requires further study (a goal of future work by myself and others).

In summary, characters that derive from patterns and topology laid down in early stages of embryonic development appear to be fixed throughout later ontogeny, irrespective of how the skeleton itself is later modified. The ‘architectural plan’, so to speak, is fixed from early embryology, and certainly by birth or hatching. While we lack a cohesive understanding of how increasingly granular cranial characters develop, known mechanisms make it clear that the architecture of the cranial skeleton is controlled by the soft tissues that are laid down long prior to the initiation of osteogenesis, which are in fact active signaling centers governing osteogenesis itself. These developmental mechanisms are broadly conserved among vertebrates, so while I welcome further studies in other extant study systems, I am confident that this general principle will hold in species that have comparable developmental programs (e.g., metamorphic amphibians may not be subject to the same constraints). It is perhaps of note that the single consistent cranial character differentiating lions from tigers (a textbook example of osteologically similar living species) is an architectural character – the posterior level of the nasofrontal suture, relative to the maxillofrontal suture (Williams et al. 2015). Similarly, Voris et al. (2022) found that an overwhelming majority of apparently ontogenetically invariant characters among derived tyrannosaurid dinosaurs related to the sutural relations of skull bones. In contrast, my results show that non-architectural characters such as the shape of bone contours, the gross proportions of the skull, dermal bone ornamentation, and muscle scar appearance are characterized by high degrees of ontogenetic variation. These traits have been identified as ontogenetic by generations of anatomists before me; I consider the significant result of the present work identifying not which characters do vary during ontogeny, but rather the identification of those that do not.

I conclude that the answer to the final question posed in the introduction to this manuscript is “yes” – ontogenetically invariant characters both exist and occur following basic principles of embryonic development. The presence of differences in skull architecture, therefore, allows us to falsify the hypothesis that the character differences between two specimens are attributable to ontogenetic variation. I hope these results will prove relevant to continued investigations of the proposed “extreme ontogenetic changes” experienced by many extinct clades, most famously including non-avian dinosaurs, as well as taxonomic assessment of new discoveries and historical collections alike. It is certainly tempting to declare that many transformations involving ‘extreme’ ontogenetic changes are invalid, and indeed some prior research has been criticized on developmental grounds – for example, Longrich and Field (2012) criticized the proposal that *Torosaurus* is the adult form of *Triceratops* partially on the grounds that such a hypothesis would require late ontogenetic addition of osteoderms, which in modern taxa obey patterning rules laid down during embryonic development. This principle would also cast doubt on the synonymy of Late Cretaceous pachycephalosaur genera, which is partially predicated on rearrangement of osteoderm ornaments (Horner and Goodwin 2009, Goodwin and Evans 2016). However, we cannot be sure that extinct species with anatomy unlike any modern animal did not have concomitant modifications to the developmental program without corroborating evidence from the fossil record itself. Here, ‘Rosetta Stone’ specimens of potentially intermediate ontogenetic stage and exceptional preservational quality will have renewed importance, for they will allow us to test how applicable models derived from extant study systems are to extinct clades of interest. While theropod specialists may note with interest that many of the widely accepted ontogenetic character transitions in *Tyrannosaurus rex* (e.g., enclosure of the subnarial foramen by the maxilla, loss of a quadratojugal pneumatic recess, development of long medial nasofrontal processes, and loss of several maxillary tooth positions) are identified by this work as unlikely to vary during ontogeny, key specimens relevant to the *Nanotyrannus*/juvenile *Tyrannosaurus* debate remain under study by myself and others, and without the data they offer any definitive statement is premature. It is clear, however, that the results of the present study contrast sharply with prevailing interpretations of ontogeny as a “black box” that can be invoked to explain almost any character difference between specimens, and I predict that if ‘extreme ontogeny’ proves to be real, we will find that it does not involve architectural modification to the skull and its soft tissues, but rather exaggeration of ontogenetic trajectories in proportion, shape, and ornamentation that exist in all amniotes.

The above discussion warrants a final word of caution. Establishing that a character is not *ontogenetically* variable is not equivalent to establishing that it is not *intraspecifically* variable. The overwhelming focus on ontogeny belies the numerous sources of intraspecific variation that may exist superimposed upon each other. Trait polymorphism is perhaps the most obvious, and as seen above can allow even ontogenetically invariant characters to vary within species. For example, the hypoglossal canal in the human skull is partly or fully divided by a bony septum in ∼20% of individuals (Guarna et al. 2023, Kalthur et al. 2023). This occurs due to abnormal development of the neurovasculature traversing the canal and is therefore almost surely not ontogenetic variation, but normal polymorphism within *Homo sapiens*. Indeed, evolution cannot occur in the absence of intraspecific variation, so we should expect that every phylogenetically informative character was, at some point in evolutionary history, polymorphic within a single species. For this reason, I urge circumspection in applying my results to the fossil record. The presence of a putatively ontogenetically invariant difference between two specimens need not mean that the two specimens belong to different species – this difference could always represent polymorphism, especially if there is only one such difference. That said, the present dataset does suggest that polymorphism is uncommon in ontogenetically invariant characters. This is probably not coincidental. Modifying architectural traits necessarily involves alteration to early steps in the developmental program, which increases the risk that they render the embryo nonviable – therefore, we may expect that these changes are developmentally “harder” and would appear less frequently, making it most likely that such character changes are associated with deeper lineage divergences, and thus that they are strong evidence of species distinction. If this model holds, these characters would also be those that are least likely to be homoplastic, and therefore they may be the strongest characters for both taxonomic hypothesis testing and phylogenetic analyses. The long-recognized importance of cranial arterial and venous anatomy in mammalian systematics (e.g., Wible 1986, Wible 1987, MacPhee et al. 2023, Crowell et al. 2024) corroborates this expectation. Recent research has demonstrated that among the appendicular skeleton, later-developing distal limb bones show greater variation that earlier- developing proximal elements, and evolve more rapidly (Stepanova and Womack 2020, Rothier et al. 2023), lending further credence to this interpretation. At present, it seems safe to say that early-developing architectural characters are likely to both be ontogenetically invariant and show limited polymorphism, but further research on polymorphism itself is sure to modify how paleontologists may accommodate it when interpreting the fossil record.

While polymorphism remains a potentially problematic unknown, the present results do allow me to articulate guidelines for vetting the possibility of fossil specimens representing an ontogenetic series:

1. Characters relating to the sutural topology of the skull, which is determined by patterns of gene expression in epithelia from early stages of embryonic development, are unlikely to change in posthatching ontogeny. This includes characters such as “bone X contacts bone Y”, “bone X has a distinct process overlapping part of bone Y”, and “bone X reaches the posterior level of landmark Z”, and “process A of bone X is longer than process B of bone Y”, among potential others.
2. Characters relating to the pathway of major cranial blood vessels and nerves, which are laid down prior to and regulate osteogenesis, are unlikely to change during posthatching ontogeny. These include, for example, “foramen bounded by bones X and Y, between bones X and Z, or entirely within bones X, Y, or Z”, “foramen in bone X is single or divided”, “foramen in bone X is on the lateral or anterior surface of bone X”, or “foramen for neurovascular bundle A is present or not present”.
3. Characters related to the distribution of pneumatic recesses (osteological correlates of air sinuses, which begin to form prior to osteogenesis) are unlikely to vary during posthatching ontogeny, with the possible exception of the very earliest stages of posthatching life. Such characters may include “pneumatic recess present in bone X” and “pneumatic recesses A and B internally communicate”, potentially among others. While the ultimate adult morphology and size of pneumatic recesses may vary, we may also predict that such change will be unidirectional (and result in larger overall size), so shrinking or secondary closure of pneumatic recesses are also unlikely.
4. Tooth counts show clade-specific patterns in ontogenetic variability, but such tendencies either involve ontogenetic addition of tooth positions, or ontogenetically stable tooth positions. In the latter case, at least, tooth count usually varies by ±1 from a modal value, and the modal value is much more common than the others. No clade shows a tendency towards a loss of tooth positions, despite published claims to the contrary.
5. Ontogenetically invariant characters overwhelmingly represent characters that arise early in embryonic development. As such, they are also less likely to be polymorphic among species, though it is impossible to fully exclude the possibility that any one character varies among a sample due to polymorphic intraspecific variation.
6. Ontogenetically variable characters, such as bone surface ornamentation, suture fusion, muscle scar development, and proportional and shape changes, tend to show a unidirectional and predictable trend, allowing apparent trend reversals to be independent indicators of species distinction among sufficiently large datasets. E.g., “bone X has an ornamental structure” may be on ontogenetically variable character state, but is unlikely to be the *juvenile* character state; conversely, “bone X lacks a muscle scar” is unlikely to represent the *adult* state.

Both critical appraisal of these predictions in a range of study systems, and through finer resolution of molecular signaling pathways that ultimately generate skeletal morphology, clearly represent important topics for future research.

## Conclusions

Despite benefitting from decades of methodological development and an increasingly rich body of fossil evidence, vertebrate paleontologists have encountered persistent difficulty with the ontogeny problem – which states, fundamentally, that ontogenetic character change may obscure the species identity of a specimen. In many cases, paleontologists have argued back and forth over the same suite of characters, because the ontogeny problem is fundamentally one of interpretation. Perhaps in response to this longstanding challenge, paleontologists have adopted a range of epistemological assumptions and analytical approaches that (implicitly or explicitly) attempt to avoid type I taxonomic error, at the expense of a very high risk of type II error. In our effort to avoid recognizing species based on growth stages of one taxon, we have put ourselves at risk of not recognizing many species that are truly represented in the known fossil record. In general, only glaring differences between putative species are accepted as evidence that they are distinct (and, given recent proposals of “extreme ontogeny”, even major anatomical differences are not always interpreted as phylogenetic in origin). Regardless, such an approach ignores the reality that closely related and osteologically similar species coexist in modern faunas, a pattern that holds over a range of clades, trophic levels, and biomes. I have heard colleagues suggest in conversation that a paleontological species should be recognized as something fundamentally different than an extant species – a broader unit, perhaps more equivalent to a modern genus, or even a more inclusive group – to account for the fact that so many closely related modern species are difficult to tell apart from skeletal evidence alone. I disagree. Species are the operational unit of evolutionary biology, and if paleontologists are to sit at the metaphorical “high table” (Maynard Smith 1984) and bring their data to bear on major questions about the evolution of life on Earth, we must at least *try* to bring our species concept in line with the modern. Though we cannot apply any species concept predicated on either behavioral or genomic data, we can use those species concepts to develop explicit expectations for how members of a species should vary from one another due to ontogeny and other sources of individual variation. In short, we should test taxonomic hypotheses by quantifying not how different two specimens are, but by how those two specimens are different.

My study of ontogeny, polymorphism, and phylogenetic variation among *Alligator* reveals that even young juvenile animals can be confidently identified based upon a suite of ontogenetically invariant traits, many of which are not polymorphic. These invariant characters overwhelmingly reflect early events in embryonic development, potentially explaining both why they are so infrequently polymorphic and why they do not change later in ontogeny – the architecture, so to speak, of the skull is set early, with later ontogenetic modifications to shape, proportions, and bone surface features superimposed upon, but not altering, the pre-existing plan of the tissues of the cranium. Ontogeny is perhaps best considered as analogous to remodeling a home, a process which may significantly alter the aesthetics and function of a space while preserving architectural features such as support beams, plumbing, and electrical wiring, rather than wholesale renovation. While a full survey of ontogenetic and polymorphism in living amniotes would require a field-wide effort, the sequence of the amniote developmental program is conservative, so I do consider my results applicable to the ontogeny problem writ large.

Indeed, anecdotal data from other systems do suggest that architectural differences, when present, indicate that different individuals belong to different species, if not more inclusive clades.

Morphology results from development, and in my opinion any anatomical trait must be considered in the context of how it develops. Evolutionary developmental biology has facilitated major breakthroughs in our understanding of vertebrate evolution because it allows us to interpret the fossil record from a mechanistic perspective that is otherwise unattainable; however, explicit integration of these observations into more strictly “paleontological” research remain rare. Taxonomy is fundamental to all else in evolutionary biology, and taxonomic hypotheses, despite being derided as “boring” or “unimportant” by some contingents, have fundamental importance to virtually all other scientific questions – perhaps explaining why species delimitation has been, and continues to be, an active area of research. Integrating mechanistic insights derived from evolutionary developmental biology and documented ontogenetic variation among living species of known identity into the paleontological toolkit offers tremendous potential to resolve the “ontogeny problem” and to, at long last, put persistent taxonomic debates to rest.

## Supporting information

Character Matrix

## Acknowledgements

This work represents a large portion of my PhD thesis, written at the Richard Gilder Graduate School of the American Museum of Natural History. I am indebted to many, both at the AMNH and other institutions, who facilitated my development as a scientist by supporting me, challenging me, and teaching me so much I did not know when I started as a student. These individuals include M. Norell, J. Flynn, J. Meng, M. Hopkins, W. Harcourt-Smith, A. Turner, B-A. Bhullar, F. Burbrink, and A. Watanabe. My doctoral committee members in particular (M. Norell, J. Flynn, J. Meng, M. Hopkins, W. Harcourt-Smith, and A. Turner) deserve special thanks for reading and revising multiple prior versions of this manuscript, which significantly improved its quality. I further thank L. Zanno and the North Carolina Museum of Natural Sciences Paleontology research group, under whose guidance and in whose company both I and this manuscript matured. This work would not have been possible without the support of AMNH collections and facilities staff including M. Chase, A. Smith (AMNH MIF), C. Mehling, C. Merril, S. Goldberg (AMNH Paleontology), and D. Kizirian and L. Vonnahme (AMNH Herpetology). Science is honed by discussion and debate, and my friends and colleagues E. Tschopp, K. Chapelle, S. Johnston, M. Fabbri, A. Ruebenstahl, and D. Meyer have all played an important role in the refinement of the ideas presented herein. Of all my compatriots, however, none deserves more thanks than A. Zietlow, who was a constant source of support and friendship, and a frequent sparring partner in debates that greatly improved the quality of the research described in this manuscript – remaining supportive even when it became clear that my results opposed her own previously published work, as all scientists imagine they will, but few do. I further thank her for coining the wonderful turn of phrase “sympatry and similarity do not equal synonymy”, which I have used in this manuscript with permission. Finally, I thank K. Williams, who has been with me on my scientific journey since the day we met as prospective graduate students, and my family, for their lifetime of love, support, and encouragement. I could not have completed this work without them.

## Footnotes

1 Perhaps due to the abundance of osteological maturity indicators in mammals (e.g., epiphyseal fusion, cranial suture fusion, dental eruption sequence), this does not seem to have become a topic of significant interest in mammalian paleontology. Thus, this manuscript focuses mostly on reptiles, with an emphasis on non-avian dinosaurs (where the questions addressed herein have been most problematic).

