## Supplemental Appendix II for "Resolving the “Ontogeny Problem” in Vertebrate Paleontology"

Supplemental Appendix II - Resolving the “Ontogeny Problem” in Vertebrate Paleontology

James G. Napoli

Results of Geometric Morphometrics Sensitivity Analyses
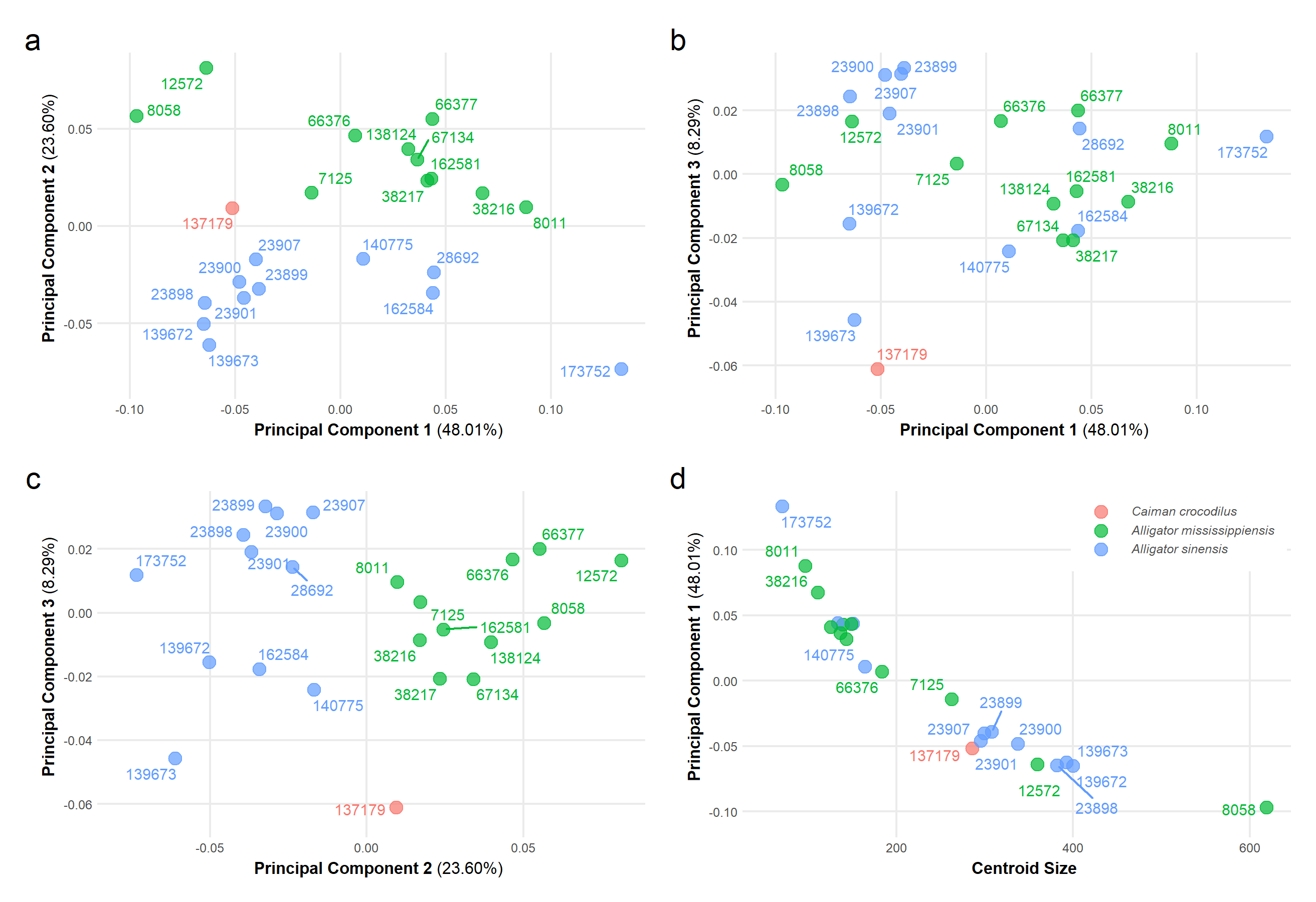

**Figure S1.** Results of geometric morphometrics analysis of 3D landmark dataset excluding all juveniles except the exemplar specimens, showing a) principal components one and two; b) principal components one and three; c) principal components two and three; d) centroid size versus principal component one.

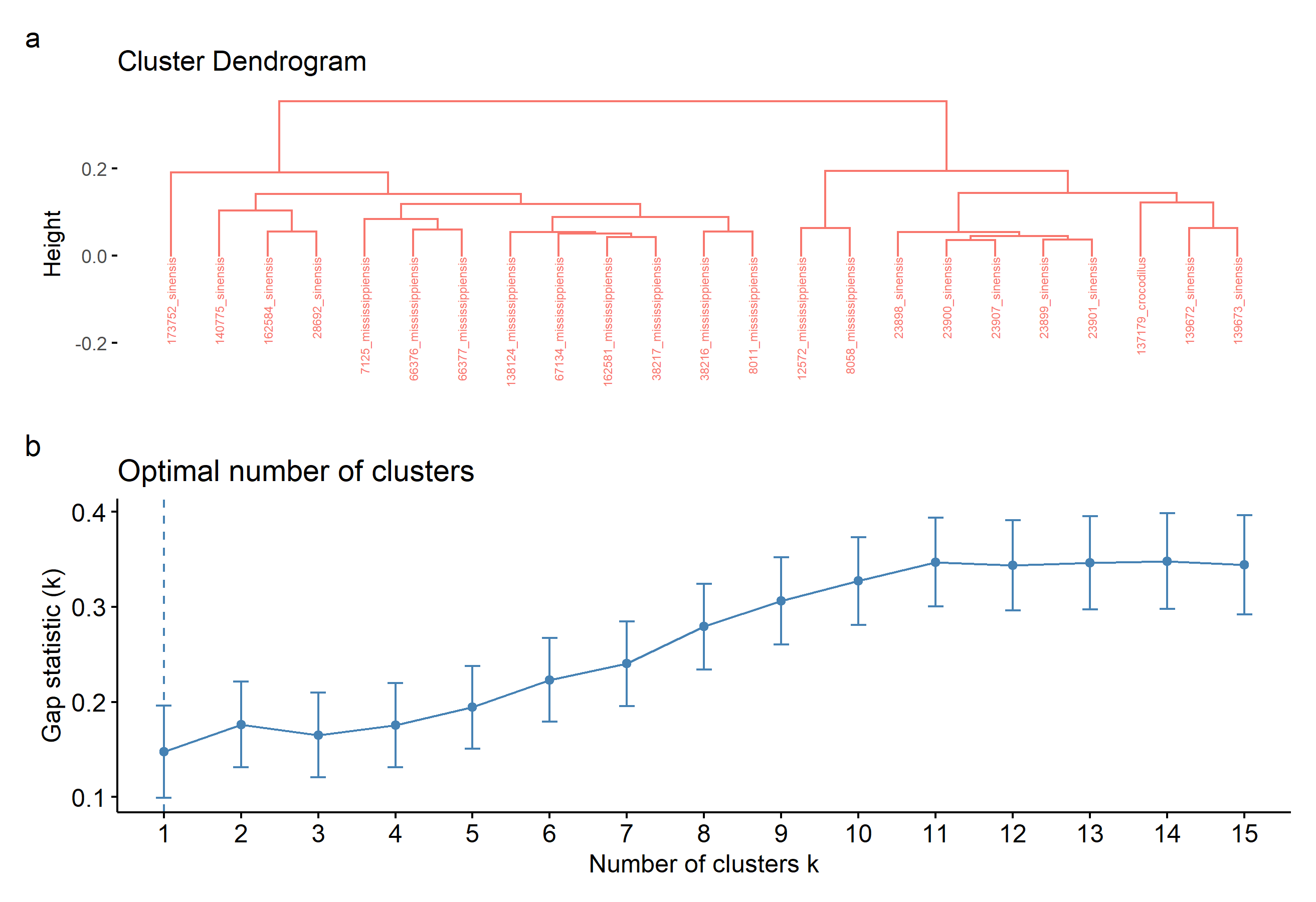

**Figure S2.** Results of agglomerative hierarchical clustering analysis on PC scores of individual specimens with all juveniles except exemplars excluded, including a) a cluster dendrogram showing the relative similarity of the included specimens and b) gap statistic plot showing the optimal number of clusters describing the dataset. A single cluster best describes the data.

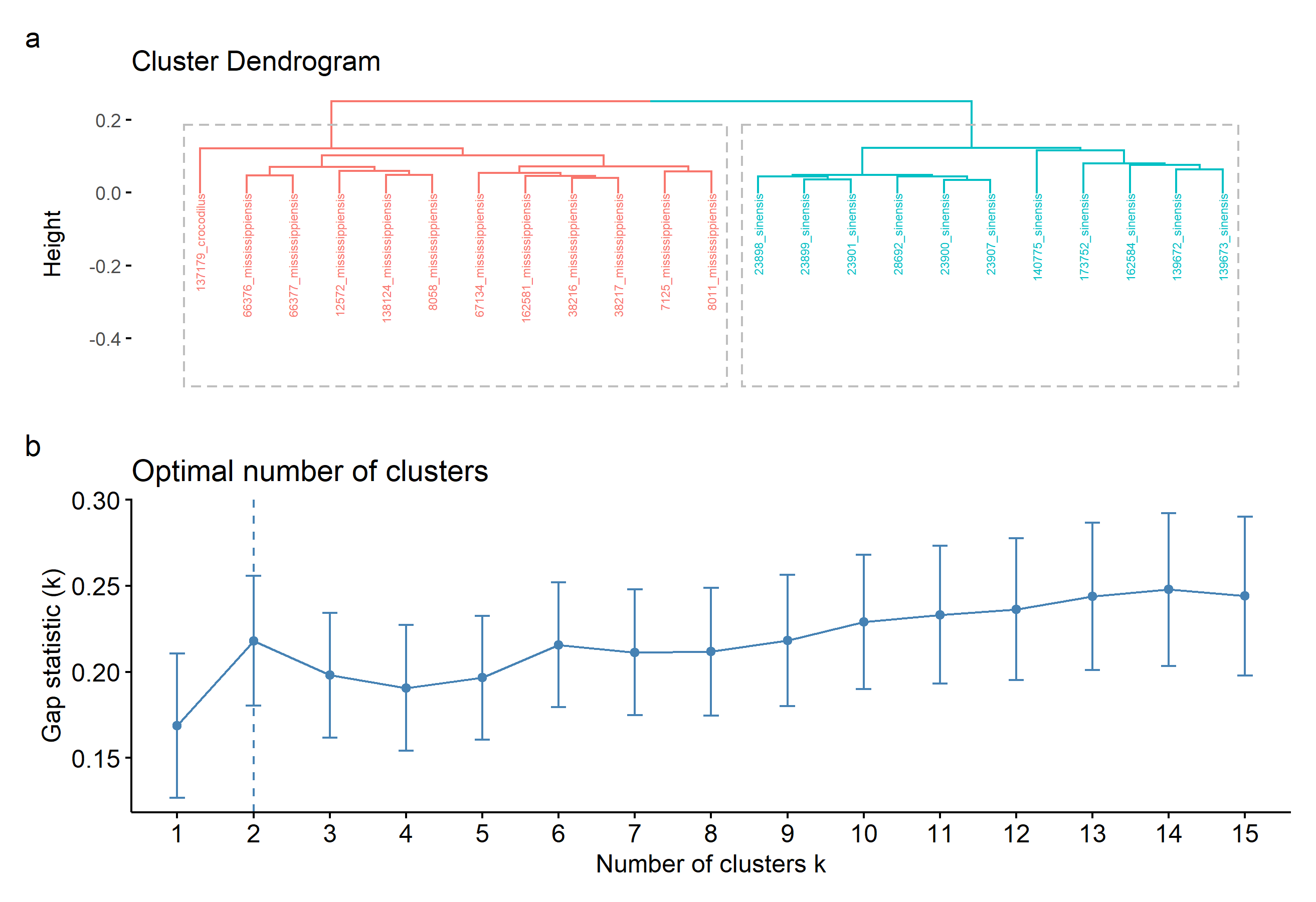

**Figure S3.** Results of agglomerative hierarchical clustering analysis on PC scores of individual specimens with PC1 (highly correlated with size) and all juveniles except exemplars excluded, including a) a cluster dendrogram showing the relative similarity of the included specimens and b) gap statistic plot showing the optimal number of clusters describing the dataset (two).

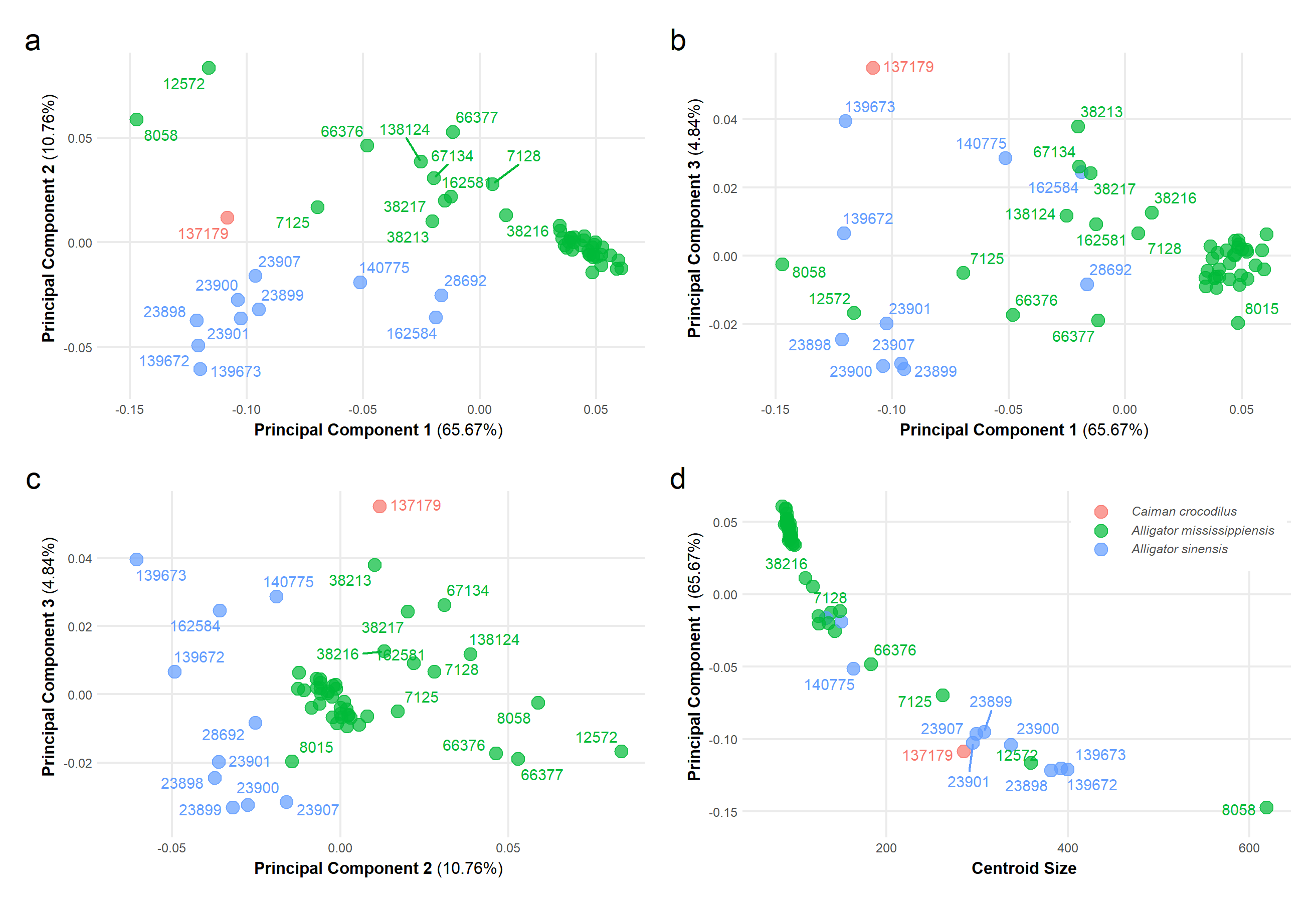

**Figure S4**. Results of geometric morphometrics analysis of 3D landmark dataset with juvenile *Alligator sinensis* excluded, showing a) principal components one and two; b) principal components one and three; c) principal components two and three; d) centroid size versus principal component one.

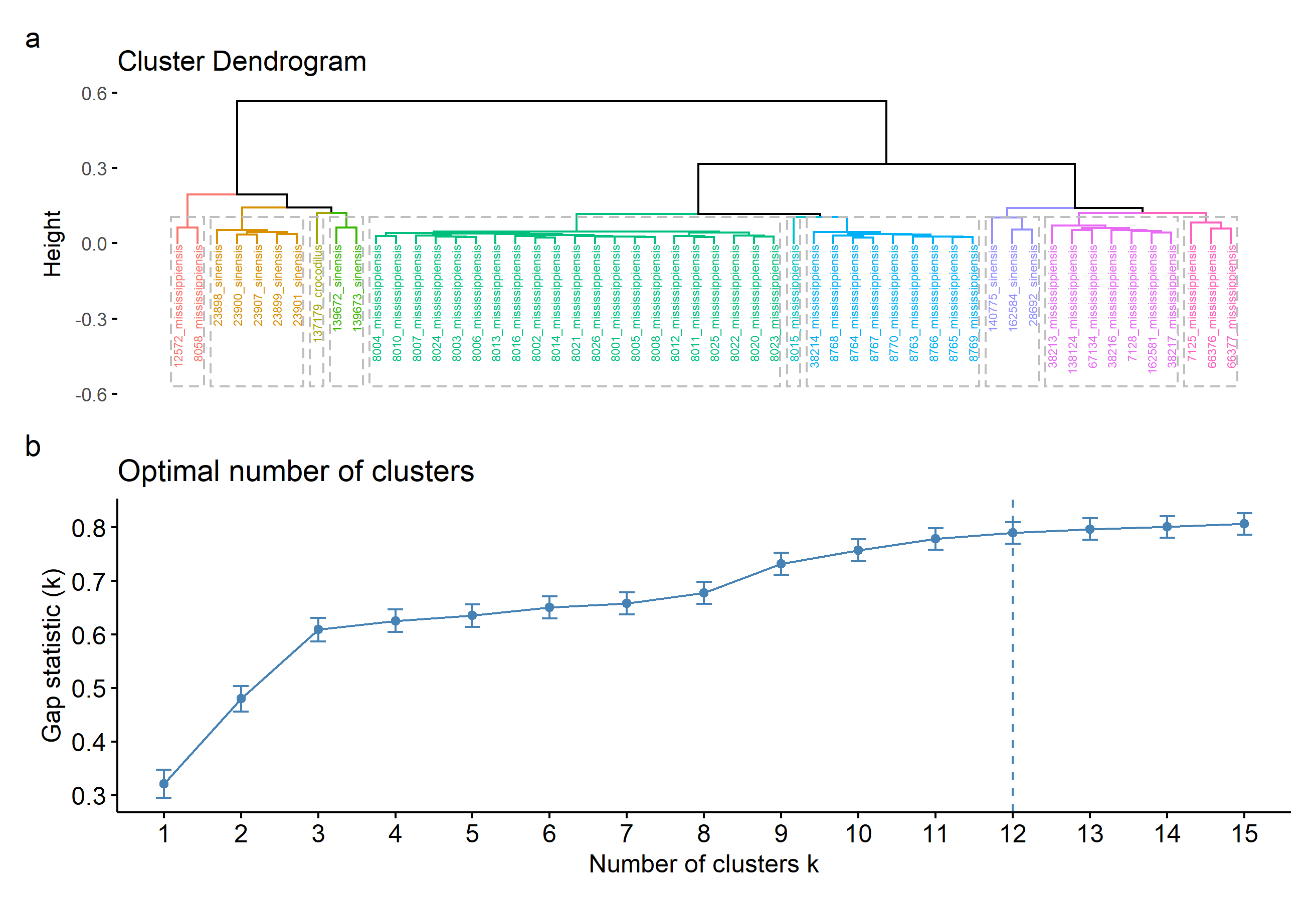

**Figure S5.** Results of agglomerative hierarchical clustering analysis on PC scores of individual specimens with juvenile *Alligator sinensis* excluded, including a) a cluster dendrogram showing the relative similarity of the included specimens and b) gap statistic plot showing the optimal number of clusters describing the dataset. Branches in panel a) are colored by cluster, with dotted lines surrounding the labels of each.

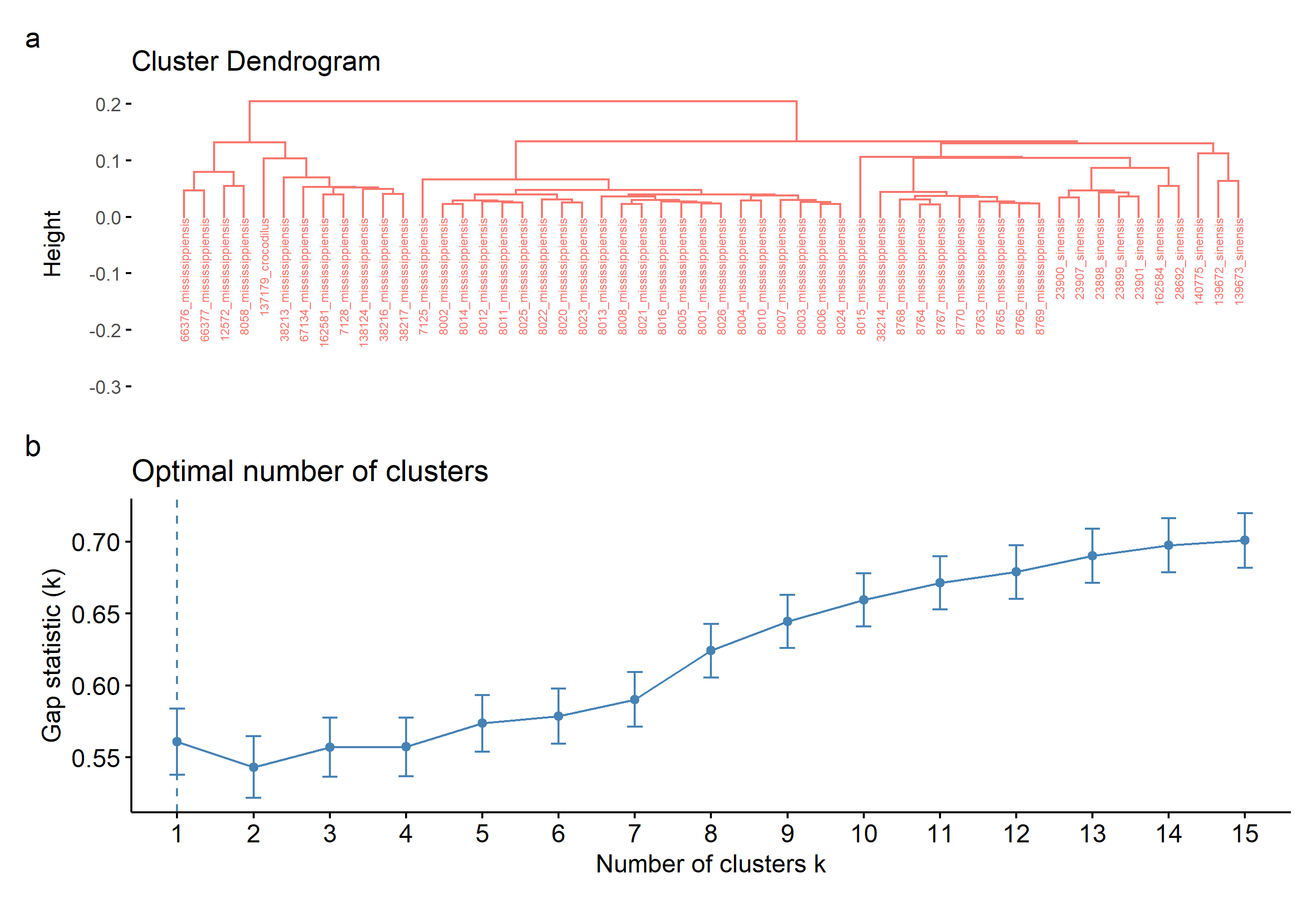

**Figure S6.** Results of agglomerative hierarchical clustering analysis on PC scores of in­dividual specimens with PC1 (highly correlated with size) and juvenile *Alligator sinensis* excluded, including a) a cluster dendrogram showing the relative similarity of the includ­ed specimens and b) gap statistic plot showing the optimal number of clusters describing the dataset. The dataset is best described by a single cluster.

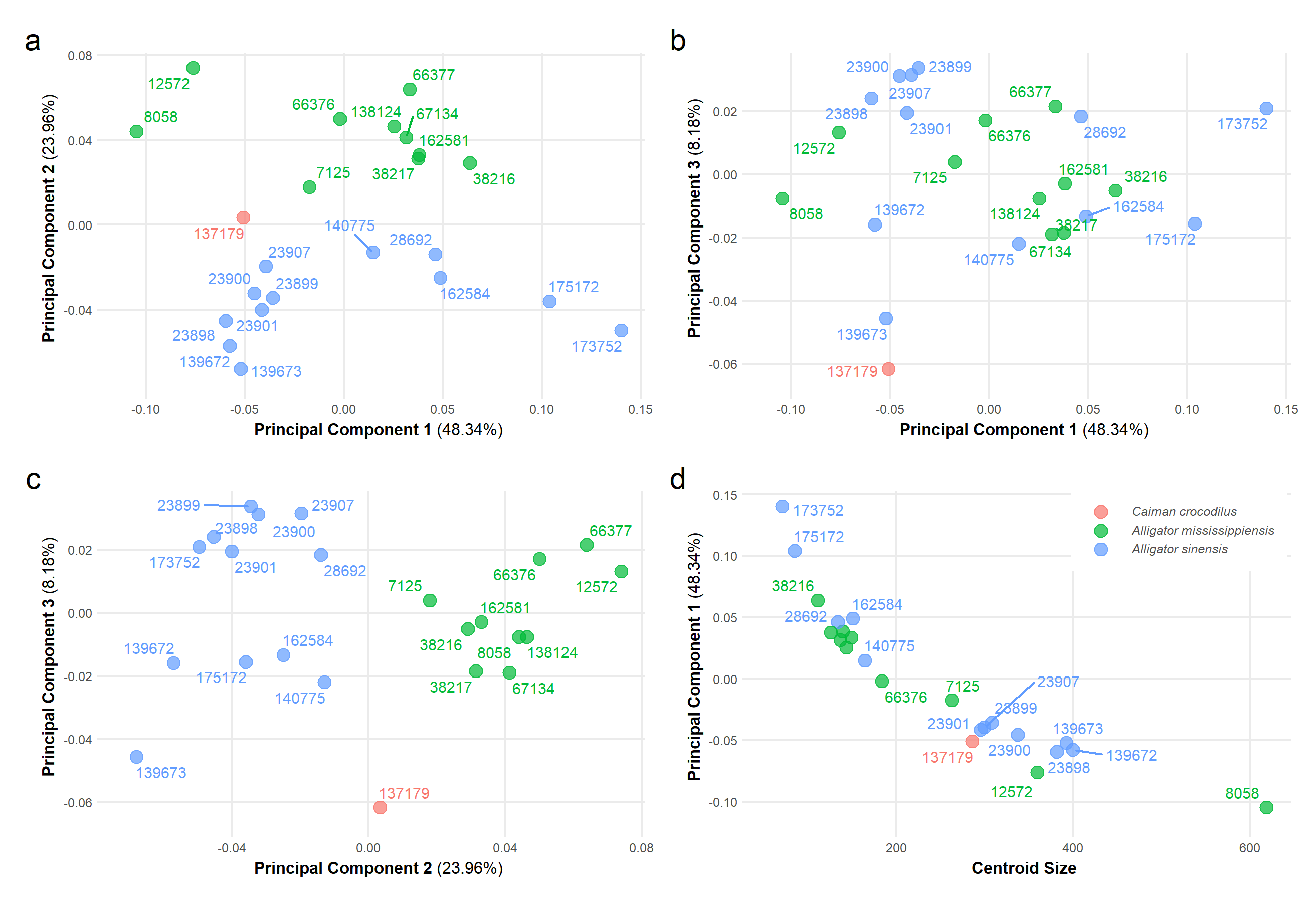

**Figure S7.** Results of geometric morphometrics analysis of 3D landmark dataset with juvenile *Alligator mississippiensis* excluded, showing a) principal components one and two; b) principal components one and three; c) principal components two and three; d) centroid size versus principal component one.

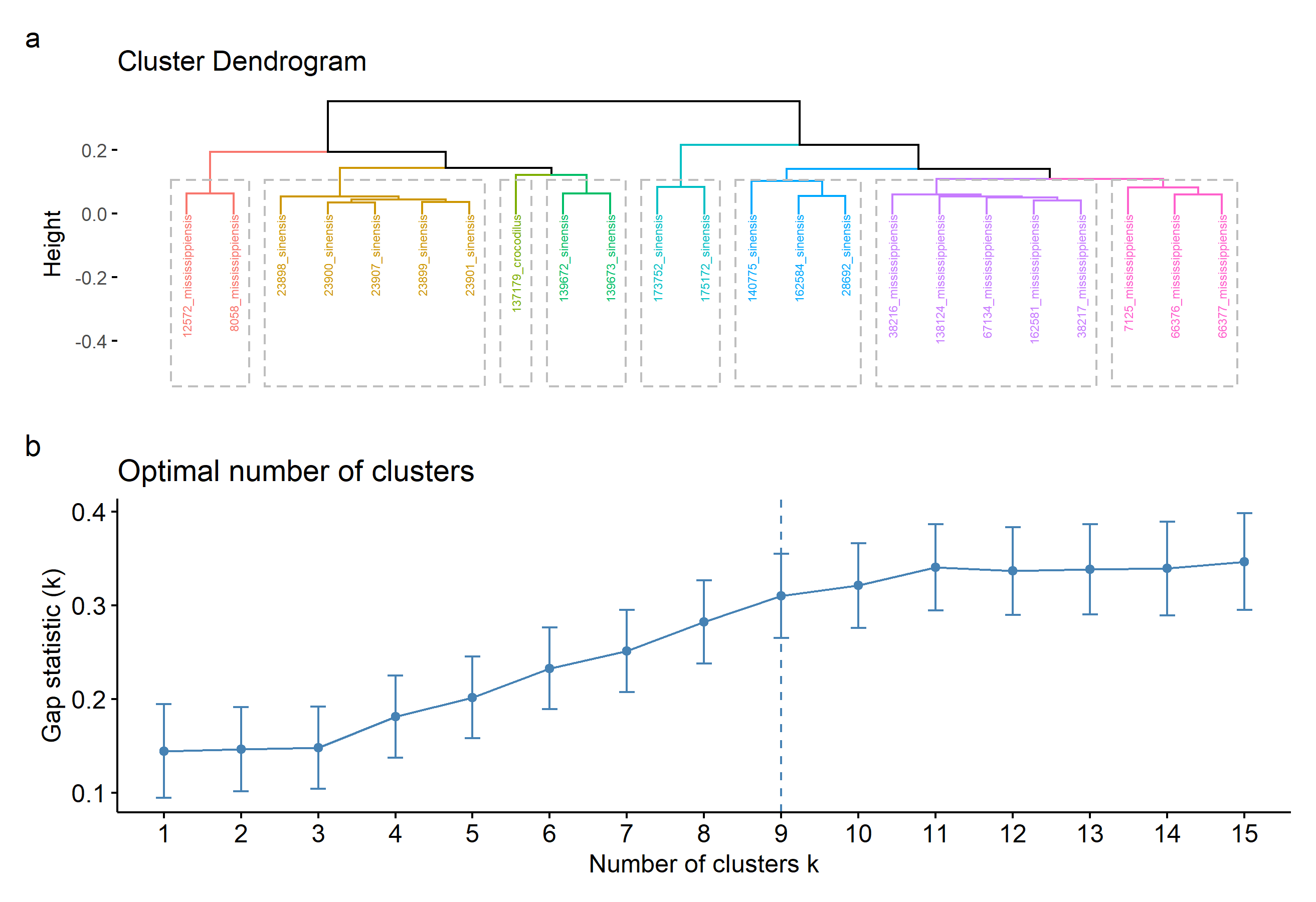

**Figure S8.** Results of agglomerative hierarchical clustering analysis on PC scores of individual specimens with juvenile *Alligator mississippiensis* excluded, including a) a cluster dendrogram showing the relative similarity of the included specimens and b) gap statistic plot showing the optimal number of clusters describing the dataset. Branches in panel a) are colored by cluster, with dotted lines surrounding the labels of each.

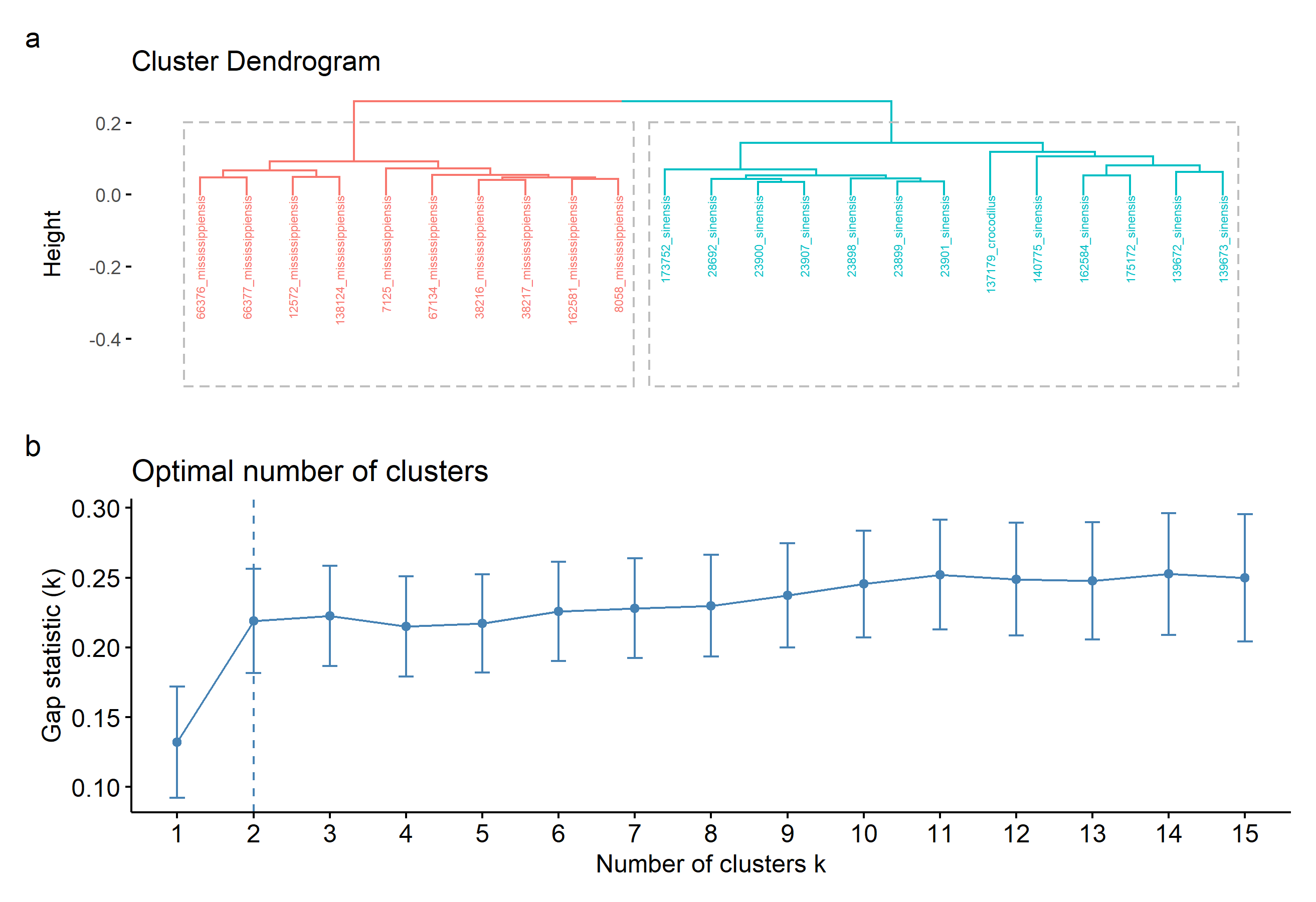

**Figure S9.** Results of agglomerative hierarchical clustering analysis on PC scores of in­dividual specimens with PC1 (highly correlated with size) and juvenile *Alligator missis­sippiensis* excluded, including a) a cluster dendrogram showing the relative similarity of the included specimens and b) gap statistic plot showing the optimal number of clusters describing the dataset (two).

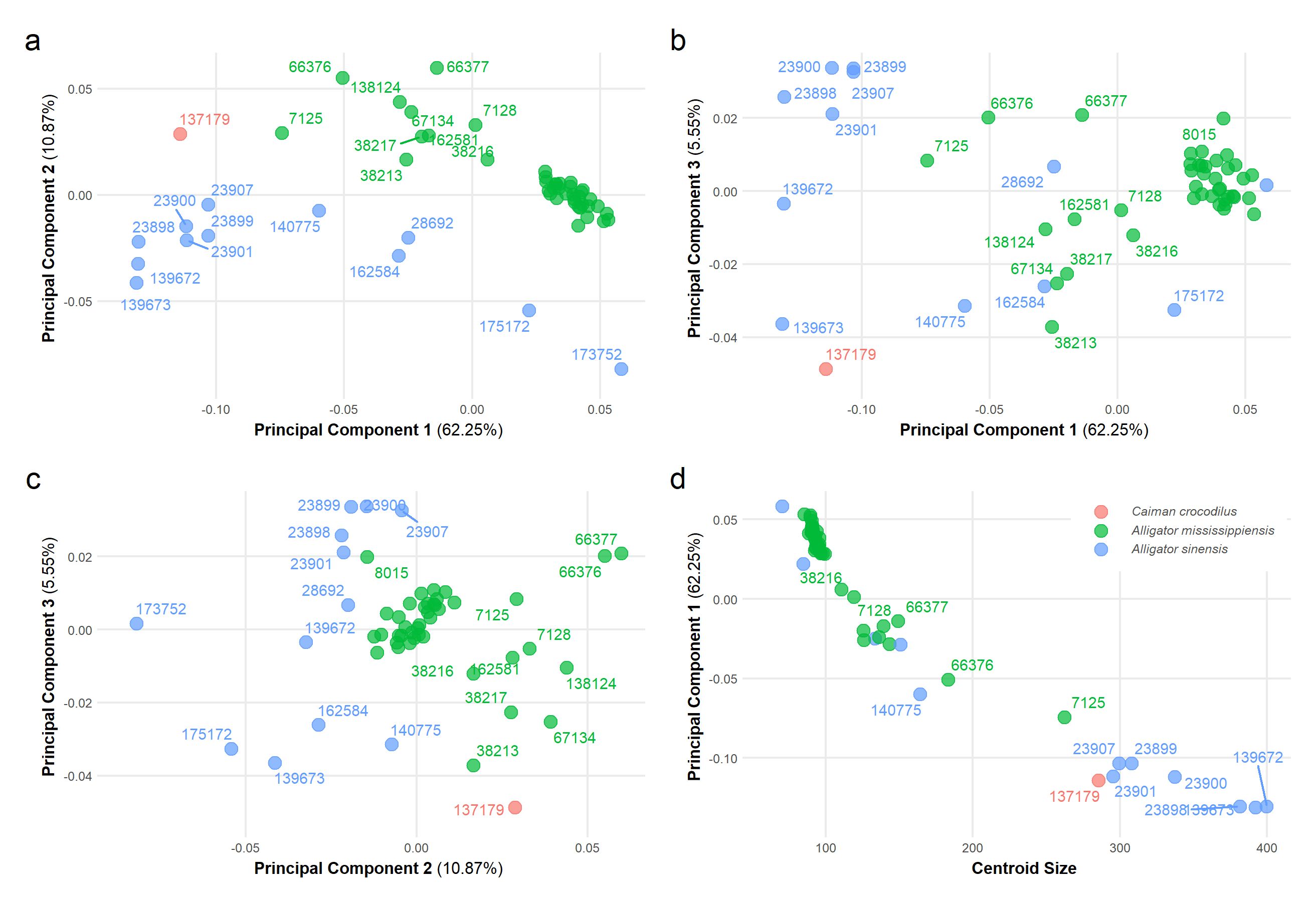

**Figure S10.** Results of geometric morphometrics analysis of 3D landmark dataset with adult *Alligator mississippiensis* excluded, showing a) principal components one and two; b) principal components one and three; c) principal components two and three; d) cen­troid size versus principal component one.

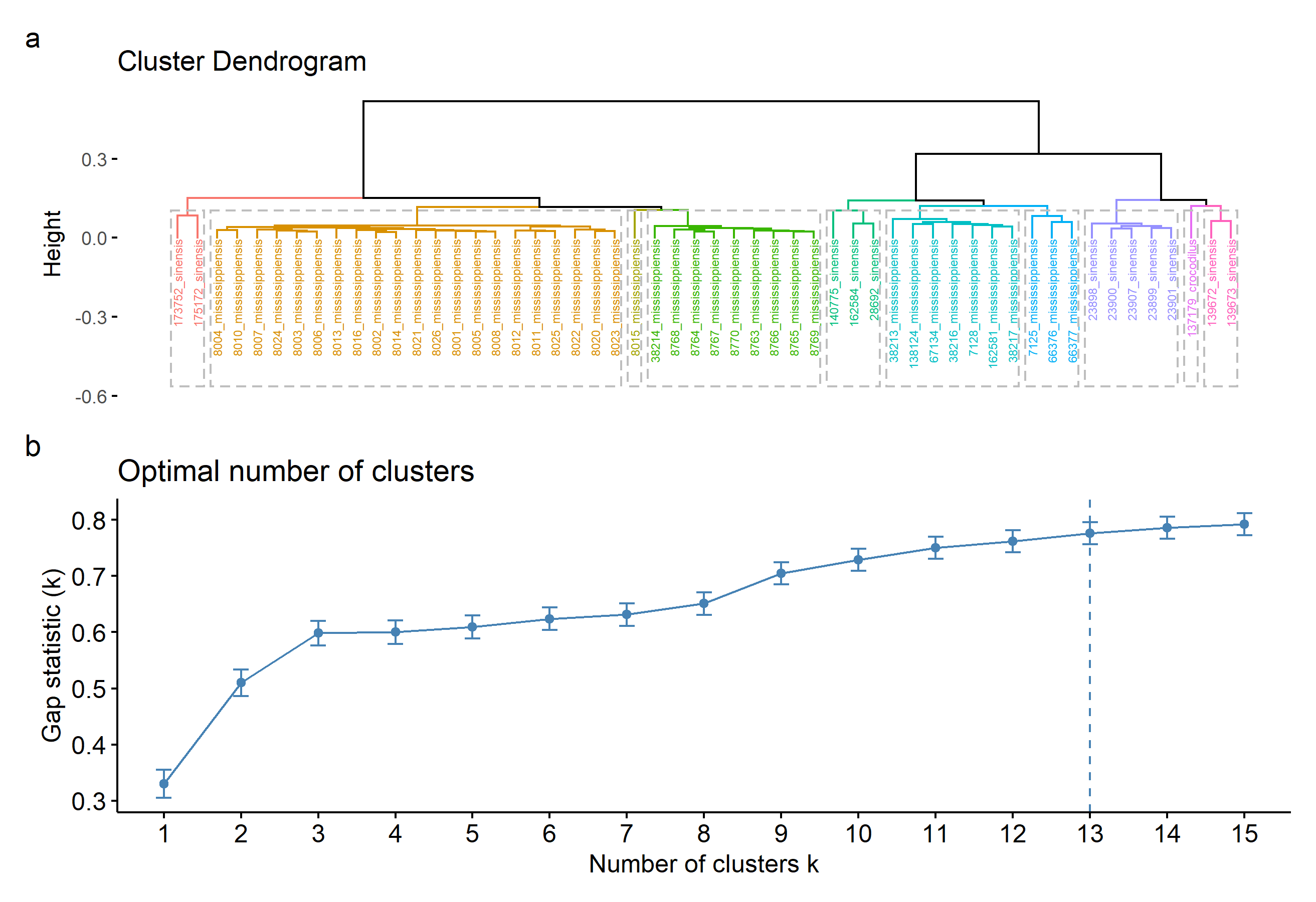

**Figure S11.** Results of agglomerative hierarchical clustering analysis on PC scores of in­dividual specimens with adult *Alligator mississippiensis* excluded, including a) a cluster dendrogram showing the relative similarity of the included specimens and b) gap statistic plot showing the optimal number of clusters describing the dataset. Branches in panel a) are colored by cluster, with dotted lines surrounding the labels of each.

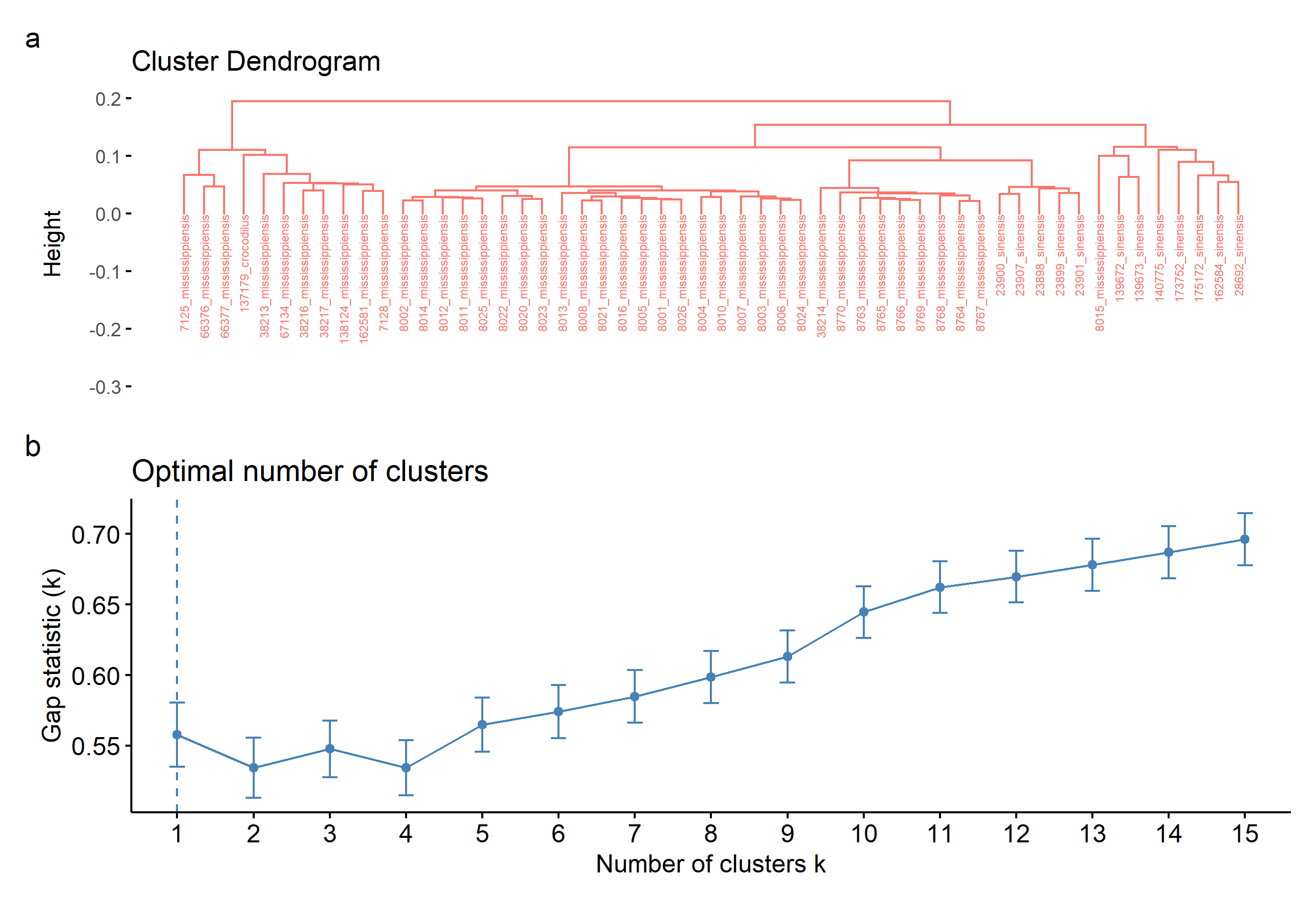

**Figure S12.** Results of agglomerative hierarchical clustering analysis on PC scores of individual specimens with PC1 (highly correlated with size) and adult *Alligator missis­sippiensis* excluded, including a) a cluster dendrogram showing the relative similarity of the included specimens and b) gap statistic plot showing the optimal number of clusters describing the dataset.

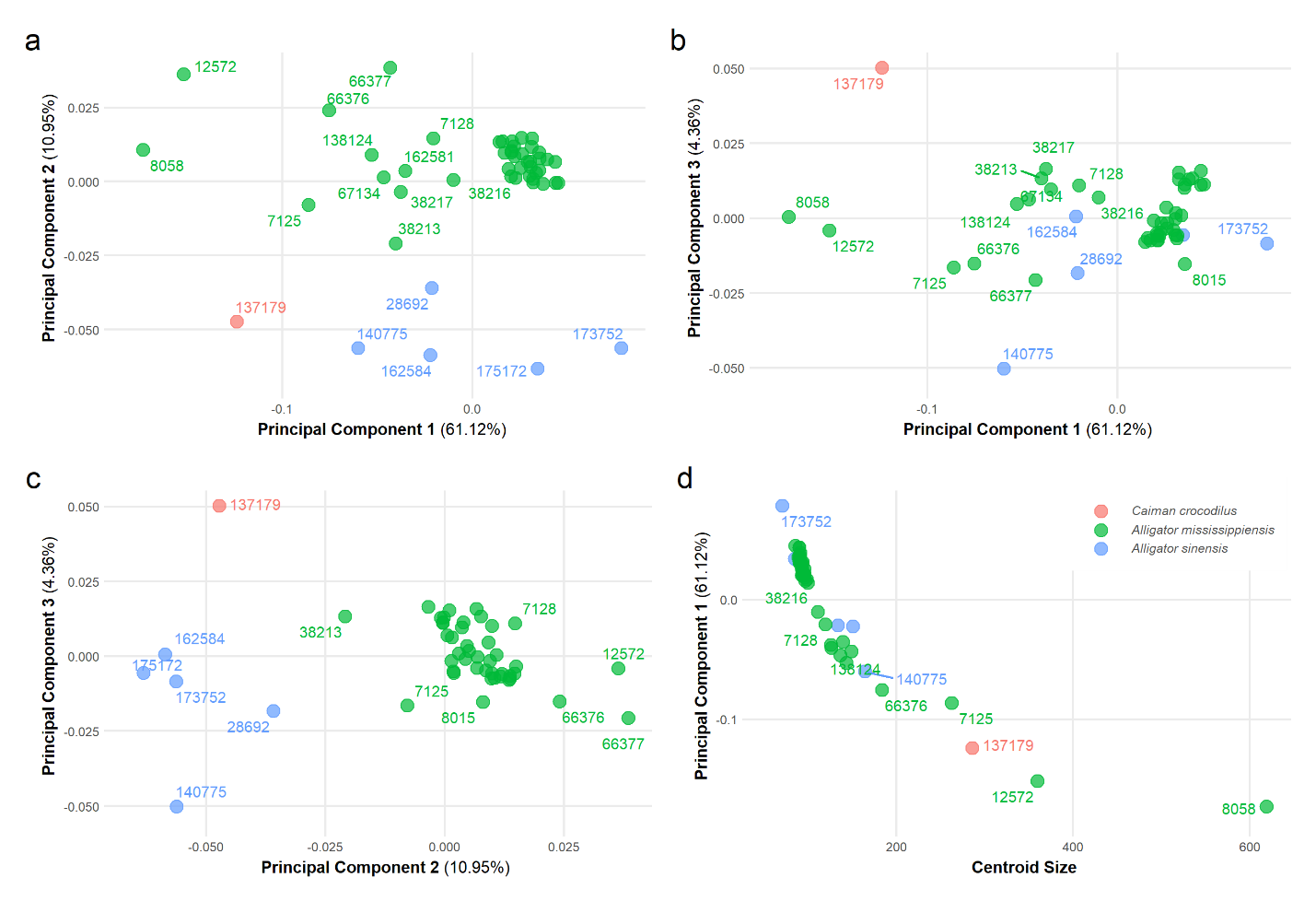

**Figure S13.** Results of geometric morphometrics analysis of 3D landmark dataset with adult *Alligator sinensis* excluded, showing a) principal components one and two; b) prin­cipal components one and three; c) principal components two and three; d) centroid size versus principal component one.

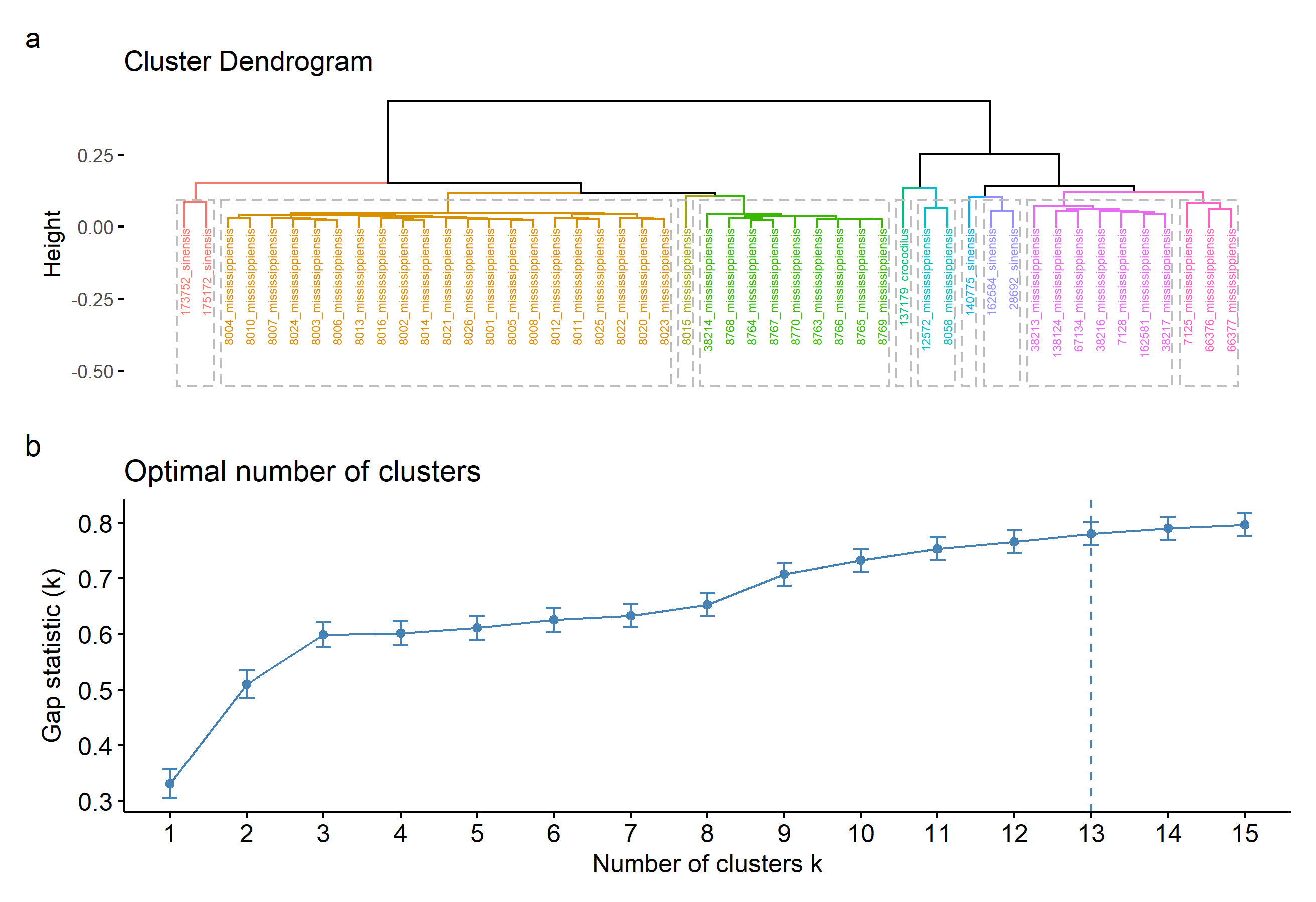

**Figure S14.** Results of agglomerative hierarchical clustering analysis on PC scores of individual specimens with adult *Alligator sinensis* excluded, including a) a cluster den­drogram showing the relative similarity of the included specimens and b) gap statistic plot showing the optimal number of clusters describing the dataset. Branches in panel a) are colored by cluster, with dotted lines surrounding the labels of each.

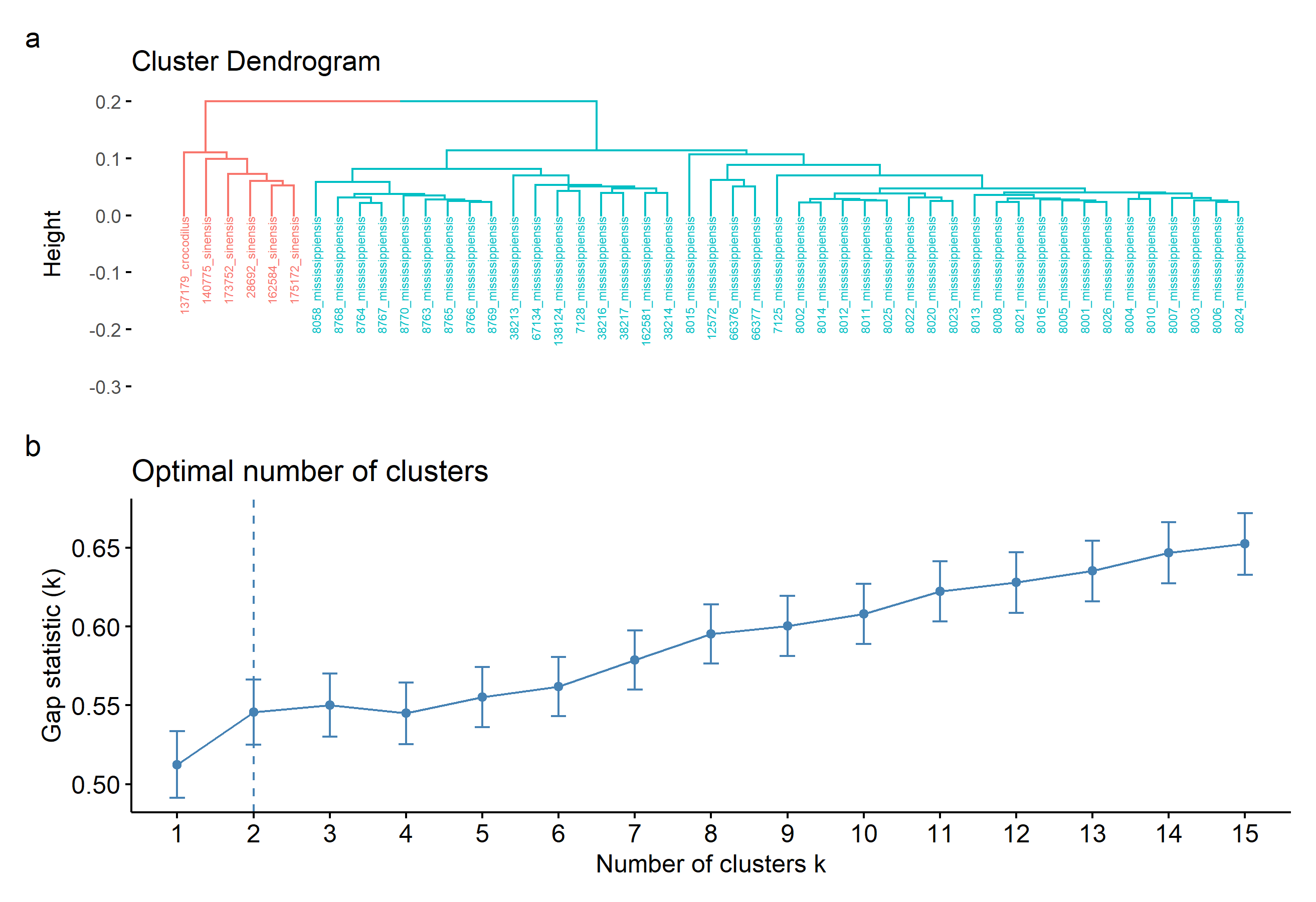
**Figure S15.** Results of agglomerative hierarchical clustering analysis on PC scores of individual specimens with PC1 (highly correlated with size) and adult *Alligator sinensis* excluded, including a) a cluster dendrogram showing the relative similarity of the includ­ed specimens and b) gap statistic plot showing the optimal number of clusters describing the dataset.
