## Supplemental Appendix I for "Resolving the “Ontogeny Problem” in Vertebrate Paleontology"

Supplemental Appendix I - Resolving the “Ontogeny Problem” in Vertebrate Paleontology

James G. Napoli

Ontogenetic and phylogenetic character list

The following character list enumerates characters and states observed in comparison of juvenile and adult members of the two extant species of *Alligator*. Characters treated as ordered in my cladistic analysis of ontogeny are indicated as such.

1. Premaxilla, proportions: greater than 70% as wide as long (0); less than or equal to 70% as wide as long (1).
2. Premaxilla, incisive foramen, mediolateral width: narrow, such that the nasal process does not extend medial to the incisive foramen (0); wide, such that the nasal process does extend medial to the incisive foramen (1).
3. Premaxilla, nasal process, shape and orientation: straight, projecting posterodorsally (0); curved, projecting posterodorsally at its base and posteriorly at its tip (1).
4. Premaxilla, external naris, anterior margin, orientation: anteromedial (0); strictly mediolateral (1).
5. Premaxilla, external naris, lateral rim, form: sharp lamina (0); inflated convexity (1).
6. Premaxilla, premaxillary body, surface of bone ventral to posterior extent of lateral rim, shape: smoothly convex, in line with remainder of external surface of premaxilla (0); strongly excavated below level of rest of bone and overhung by lateral rim of external naris (1).
7. Premaxilla, premaxillary body, sculpturing of bone laterally adjacent to external naris: absent (0); present (1).
8. Premaxilla, posterior process, orientation: posterodorsal (0); strictly posterior (1).
9. Premaxilla, posterior process, orientation of external surface: dorsolateral (0); dorsal (1).
10. Premaxilla, narial fossa, posterior region, small neurovascular foramen: present (0); absent (1).
11. Subnarial foramen, composition: formed by premaxilla and maxilla (0); formed by premaxilla and nasal (1).
12. Premaxilla, premaxillary teeth, orientation: procumbent (0); decumbent (1).
13. Premaxilla, palatal process, anteroposterior length posterior to incisive foramen: shorter than incisive foramen (0); longer than incisive foramen (1).
14. Premaxilla, dentigerous margin, shape in dorsal view: straight (0); strongly curved (1).
15. Premaxilla, anterior extent of joint surface for maxilla: joint surface extends anterior to anteroposterior midpoint of the premaxilla (0); joint surface terminates posterior to anteroposterior midpoint of the premaxilla (1).
16. Premaxilla, nasal process, length: extending to 33% or less of anteroposterior length of premaxilla (0); extending more than 33% of anteroposterior length of the premaxilla (1).
17. Premaxilla, incisive foramen, anteroposterior length: less than 33% of the length of the premaxilla (0); greater than 33% of the length of the premaxilla (1).
18. Premaxilla, posterior process, size: long, with a prominent exposure between nasal and maxilla (0); short, with a restricted exposure between nasal and maxilla (1).
19. Premaxilla, premaxillary teeth, orientation of first two teeth: strictly mediolateral to one another (0); second tooth is posteriorly displaced relative to first tooth (1).
20. Maxilla, joint surface for lacrimal, position of notch relative to total length of maxilla: in the anterior half of the maxilla (0); approximately at anteroposterior midpoint of maxilla (1); in the posterior half of the maxilla (2). ORDERED.
21. Maxilla, external surface, sculpturing: absent (0); moderate (1); heavy (2); extreme (3). ORDERED.
22. Maxilla, dental row, orientation: straight (0); medially deflected at anterior end, creating convex profile over fourth maxillary tooth (1).
23. Maxilla, maxillary toothrow, number of teeth: 13 (0); 14 (1); 15 (2); 16 (3). ORDERED.
24. Maxilla, jugal process, length posterior to last maxillary tooth: short (0), long, with a distinct region of post-dental bone (1).
25. Maxilla, palatine process, presence: absent (0); present (1).
26. Maxilla, palatine process, size: small (0); large (1).
27. Maxilla, palatal process, ventral view, shape of contact with palatine: straight to variably convex and directed mostly anteromedially (0); strongly squared, with a distinct “corner” marking a transition from anteroposterior to mediolateral orientation (1).
28. Maxilla, palatal process, foramina, number: two (0); one (1).
29. Maxilla, palatal process, foramina, separation: narrow (0); wide (1).
30. Maxilla, postvestibular recess, foramen, shape: circular (0); oval (1).
31. Maxilla, interfenestral strut, mediolateral thickness: narrow (0), broad (1).
32. Maxilla, lateral fenestra, size: small (0); large (1).
33. Maxilla, lateral recess, communication with caviconchal sinus: absent (0); present (1).
34. Maxilla, caviconchal recess, accessory diverticulum invading palatal process: absent (0); present (1).
35. Maxilla, jugal process, palatal shelf, articulation for ectopterygoid, form: palatal shelf of jugal process forms a socket to receive the anterior process of the ectopterygoid (0); palatal shelf does not form socket for ectopterygoid, bones articulate along a flat contact (1).
36. Maxilla, supra-alveolar canal, foramen, position: far ventral to dorsal margin of jugal process (0); nearly abutting the dorsal margin of the jugal process (1).
37. Maxilla, foramen for palatal vasculature, position: ventral to lateral fenestra (0); in the floor of the lateral recess (1).
38. Nasal, proportions: greater than 25% as wide as long (0); between 20% and 30% as wide as long (1); less than 20% as wide as long (2). ORDERED.
39. Nasal, external surface: smooth (0); lightly sculptured (1); heavily sculptured (2). ORDERED.
40. Lacrimal, body, shape: roughly square in lateral view (0); anteroposteriorly elongate in lateral view (1).
41. Lacrimal, posterodorsal process, size: barely developed (0); distinct but very short (1); about half length of jugal process (2); extending posteriorly further than the laterally exposed portion of the jugal process (3). ORDERED.
42. Lacrimal, orbital fossa, visibility in lateral view: absent (0); present (1).
43. Lacrimal, ridge bounding orbital fossa, shape: thin and sharp (0); inflated (1).
44. Lacrimal, external surface, sculpturing: absent (0); present (1).
45. Lacrimal, posterior ostium of nasolacrimal canal, position: anterior or level to posterior extent of orbital rim (0); slightly posterior to posterior extent of orbital rim (1); far posterior to posterior extent of orbital rim (2). ORDERED.
46. Lacrimal, ridge bounding orbital fossa, contact with jugal process: absent (0); present (1).
47. Lacrimal, neurovascular foramen ventral to posterior ostium of nasolacrimal duct: present (0); absent (1).
48. Prefrontal, anterior process, length relative to posterodorsal process: approximately equal (0); markedly longer (1).
49. Prefrontal, external surface, sculpturing: moderately sculptured (0); extensively sculptured (1).
50. Prefrontal, preorbital crest: absent (0); present (1).
    a. Note: all Alligator have some eminence on the dorsal surface of the prefrontal, which is likely homologous with the distinct preorbital crest found in *Caiman crocodilus* and *Alligator sinensis*. For purposes of this matrix, the crest is considered absent if it is not distinguishable from the orbital margin of the bone, a condition found in many adult *Alligator*. It should be noted that the crest does not resorb during ontogeny, but merges with an adjacent eminence such that a distinct crest is no longer present.
51. Prefrontal, preorbital crest: barely defined (0); distinct but unenlarged (1); well-developed with anteromedial excavation (2).
52. Prefrontal, prefrontal pillar, small neurovascular foramen ventral to frontal contact: absent (0); present (1).
53. Prefrontal, prefrontal recess: absent (0); present (1).
54. Prefrontal, prefrontal recess, development: incipient (0); extensive (1).
55. Prefrontal, prefrontal pillar, anterior surface, concavity: gentle (0); deeply excavated (1).
56. Prefrontal, prefrontal pillar, anterior surface, transverse ridge: absent (0); present (1).
57. Prefrontal, posterodorsal process, subcristal process: absent (0); present (1).
58. Prefrontal, descending process, presence: present (0); absent (1).
59. Prefrontal, posterodorsal process, orientation: in line with rest of bone (0); medially deflected (1).
60. Frontal, orbital margin, form and texture: thin and smooth (0); inflated and rugose (1).
61. Frontal, orbital rims, dorsoventral position: in line with skull roof between orbits (0); raised distinctly above skull roof between orbits (1).
62. Frontal, orbital rims, dorsoventral position: raised less than total thickness of combined cristae cranii and interorbital portion of the frontal (0); raised equal to or more the total thickness of the remainder of the frontal (1).
63. Frontal, dorsal surface, sculpturing: smooth (0); sculptured (1).
64. Frontal, posterior margin, shape: W-shaped (0); V-shaped to transverse, most posterior point occurs along midline (1).
65. Frontal, orbital region, length compared to total length of frontal: orbit occupies more than half the anteroposterior length of the frontal (0); orbit occupies less than half the anteroposterior length of the frontal.
66. Frontal, anterior incisure, presence: absent (0), present, such that the most anterior point on the frontal occurs lateral to the midline (1).
67. Parietal, fusion: incomplete (0); complete (1).
68. Parietal, dorsolateral margin, shape: straight (0); slightly embayed by supratemporal fossa (1); deeply embayed by supratemporal fossa, such that the median portion of the parietal is hourglass-shaped in appearance (2). ORDERED.
69. Parietal, lateral process, development: present as an articular facet that is not laterally projected from the parietal (0); present as a lateral projection that does not reach the lateral level of the rest of the parietal (1); present as a large lateral projection that extends to the same lateral level of the rest of the parietal (2). ORDERED.
70. Parietal, external surface, sculpturing: absent (0); present (1).
71. Parietal, parietal recess, medial fenestrae: separate (0); conjoined (1).
72. Parietal, parietal recess, anterior fenestrae: absent (0); present (1).
73. Parietal, supraoccipital joint surface, length compared to whole parietal: supraoccipital joint surface is less than 1/3rd the length of the parietal (0); supraoccipital joint surface is more than 1/3rd the length of the parietal (1).
74. Parietal, anterior margin, V-shaped notch in ventral lamina: absent (0); present (1).
75. Parietal, parietal recess, medial fenestrae, length: occupy more than half of the supraoccipital joint surface (0); occupy less than half of the supraoccipital joint surface (1).
76. Postorbital, dorsal surface, sculpturing: restricted to lateral edge (0); present over entire surface (1).
77. Postorbital, squamosal process, robustness: thin, less than breadth of frontal process (0); thick, equivalently broad as frontal process (1).
78. Postorbital, medial postorbital foramen, roof: thin (0); inflated (1).
79. Postorbital, border of skull table, shape: thin and sharp (0); inflated (1).
80. Postorbital, jugal process, jugal facet, orientation of apex: anterodorsal (0); posterodorsal (1).
81. Postorbital, jugal process, jugal facet, position of apex: anteriorly displaced on jugal process (0); approximately centered on jugal process (1).
82. Postorbital, quadratojugal process, length: short (0); long, approaching posterior level of squamosal process (1).
83. Postorbital, medial postorbital foramen, exposure in dorsal view: exposed (0); unexposed, completely roofed by bony lamina (1).
84. Postorbital, lateral postorbital foramen, position: anterior, anterior to apex of jugal facet (0); lateral, posterior to apex of jugal facet (1).
85. Postorbital, squamosal process, channel emanating from lateral postorbital foramen: absent (0); present (1).
86. Squamosal, dorsal surface, sculpturing: absent (0); present (1).
87. Squamosal, lateral margin of skull table: thin and sharp (0); inflated (1).
88. Squamosal, dorsal lamina roofing posterior end of supratemporal fossa: absent (0); present (1).
89. Squamosal, orientation of posterolateral boundary of supratemporal fossa: anterolateral (0); more mediolateral than anteroposterior (1).
90. Squamosal, supratemporal fossa, lateral to temporoorbital canal, neurovascular foramina, number: two (0); one (1).
91. Squamosal, medial process bounding temporoorbital canal: absent (0); present (1).
92. Squamosal, medial process bounding temporoorbital canal, size: small (0); large (1).
93. Squamosal, lateral lamina, overhang by lateral rim of skull table: limited to absent (0); marked, creating a channel on the lateral surface of the postorbital process of the squamosal but allowing lateral lamina to be seen in dorsal view (1).
94. Squamosal, lateral lamina, ventral process: indistinct, represented as a broad convexity (0); distinct, represented as a sharp triangle (1).
95. Squamosal, lateral lamina, ventral surface anterior to ventral process: sharp, no flat surface developed (0); marked by an expanded flat surface (1).
96. Squamosal, parietal process, position: markedly dorsal to rest of dorsal surface of squamosal (0); at approximately the level of the rest of the squamosal (1).
97. Squamosal, paroccipital process: ventrally directed (0); posteroventrally directed, far exceeding the posterior level of the skull table (1).
98. Jugal, anterior process, shape in dorsal view: mediolaterally bowed (0); straight (1).
99. Jugal, orbital rim of anterior process, shape: strongly laterally deflected, making the lateral surface of the bone deeply concave (0); weakly laterally deflected, making the lateral surface of the bone weakly concave (1); not laterally deflected, making the lateral surface of the bone flat (2). ORDERED.
100. Jugal, lateral surface, orientation: faces laterally (0); faces dorsolaterally (1).
101. Jugal, anterior process, depth: not markedly deeper than posterior process (0); much deeper than posterior process (1).
102. Jugal, anterior margin, shape of ventral extent: strongly squared (0); rounded (1); sloping smoothly posteroventrally (2). ORDERED.
103. Jugal, anterior margin, anterodorsal prong: absent (0); present (1).
104. Jugal, external surface, sculpturing: absent (0); present (1).
105. Jugal, medial surface, neurovascular foramina anterior to postorbital process, number: two (0); three (1).
106. Jugal, medial surface, neurovascular foramina anterior to postorbital process, size: most anterior foramen markedly smaller than others (0); all equal in size (1).
107. Jugal, joint surface for quadratojugal, separation from dorsal margin of posterior process: absent (0); present (1).
108. Jugal, lateral surface of postorbital process, large neurovascular foramen: absent (0); present (1).
109. Jugal, posterior process, shape: points mostly posteriorly, process not strongly arched (0); points posteroventrally, process is strongly arched (1).
110. Ectopterygoid, posterior process, orientation: points posteriorly (0); medially inflected (1).
111. Ectopterygoid, ventral process, length: extends halfway down pterygoid ala (0); extends about two-thirds the length of the pterygoid ala (1).
112. Ectopterygoid, ventral process, accessory medial process lapping posterior face of pterygoid: absent (0); present (1).
113. Quadratojugal, ascending process, anterior end, shape: straight (0); laterally deflected (1).
114. Quadratojugal, external surface, sculpturing: absent (0); present (1).
115. Quadratojugal, external surface, sculpturing, location: present at posterior end (0); present along entire length of element (1).
116. Quadratojugal, ascending process, quadratojugal spine: completely absent (0); present as a small convexity (1).
117. Quadrate, condyles, anteroposterior position: anteriorly positioned, located mostly or entirely anterior the posterior margin of the external otic recess (0); posteriorly positioned, located posterior to the external otic recess but not posterior to the quadrate portion of the paroccipital process (1); far posteriorly positioned, anteriormost point on quadrate condyles is at or posterior to the level of the quadrate portion of the paroccipital process (2).
118. Quadrate, condyles, mediolateral position: medially positioned, medial condyle is medial to lateral margin of the skull table (0); laterally positioned, medial condyle is at or past lateral margin of skull table in dorsal view (1).
119. Quadrate, siphonal foramen, size: large, nearly half the size of the external otic recess (0); small, approximately one-quarter the size of the external otic recess (1).
120. Quadrate, shaft, dorsal margin, shape in lateral view: flat to subtly convex (0); concave (1).
121. Quadrate, shaft, ventral surface, crest A: absent (0); present (1).
122. Quadrate, shaft, ventral surface, crest A’: absent (0); present (1).
123. Quadrate, shaft, ventral surface, rugosity at intersection of crest A and A’ (or terminus of one of these crests, if the other has not developed): absent (0); present (1).
124. Quadrate, ala, ventral surface, crest B: absent (0); present (1).
125. Quadrate, ala, ventral surface, crest B, extent: continuous (0); interrupted (1).
126. Quadrate, ala, ventral surface, crest C: absent (0); present (1).
127. Quadrate, shaft, otic buttress, size: subtle convexity (0); tall projection (1).
128. Quadrate, shaft, posterior process, shape of ventral margin: straight (0); with a discrete step (1).
129. Quadrate, shaft, external otic recess, discrete rim: absent (0); present (1).
130. Quadrate, shaft, foramen dorsal to foramen aereum: absent (0); present (1).
131. Laterosphenoid, anterior process: incompletely formed (0); complete (1).
132. Laterosphenoid, antotic crest: absent (0); present (1).
133. Laterosphenoid, capitate process, position: at same mediolateral level as body (0); far lateral to body of laterosphenoid (1).
134. Laterosphenoid, slender process, position: far separated from basisphenoid rostrum (0); approaching or contacting the basisphenoid rostrum (1).
135. Laterosphenoid, lateral bridge, orientation of long axis in lateral view: posteroventral (0); vertical (1).
136. Laterosphenoid, lateral bridge, anteroposterior width: narrow (0); wide, obscuring body of laterosphenoid from lateral view (1).
137. Laterosphenoid, lateral bridge, ventral contact with laterosphenoid body: absent (0); present, processes contact to fully enclose a channel (1).
138. Laterosphenoid, laterosphenoid recess, prootic ostium: absent (0); present (1).
139. Laterosphenoid, laterosphenoid recess, prootic ostium: large (0); small (1).
140. Laterosphenoid, laterosphenoid recess, extent: limited, does not invade body ventral to prootic ostium (0); extensive, does invade body ventral to prootic ostium (1).
141. Laterosphenoid, laterosphenoid recess, ventral ostium: absent (0); present (1).
142. Laterosphenoid, trochlear foramen, position: far posterior to optic notch in lateral view (0); adjacent to optic notch in lateral view (1).
143. Laterosphenoid, anterior process, dorsal foramina, number: one (0); two (1).
144. Laterosphenoid, foramen for supraorbital branch of trigeminal, composition: formed equally by laterosphenoid and quadrate (0); formed entirely by laterosphenoid (1).
145. Otoccipital, lateral exposure: broad (0), covered by quadrate and squamosal, exposed only at the lateral tip of the paroccicpital process (1).
146. Otoccipital, paroccipital process, length: short (0); long (1).
147. Otoccipital, anteroventral process, ventral lamina: absent or poorly developed (0); present as a large triangular lamina (1).
148. Otoccipital, joint surface for basioccipital, orientation of long axis: anteroventral (0); anteroposterior (1).
149. Otoccipital, occipital surface, shape: convex (0); flat (1).
150. Otoccipital, anteroventral process, lateral lamina, height: as tall as paroccipital process (0); distinctly shorter than paroccipital process (1).
151. Otoccipital, paroccipital process, lateralmost extent, position: dorsal half of paroccipital process (0); ventral half of paroccipital process (1).
152. Supraoccipital, ventral margin, shape: horizontal (0); triangular (1).
153. Supraoccipital, midbrain and hindbrain fossae, size: broad, occupying approximately 1/3rd the width of the supraoccipital (0); constricted, occupying approximately ¼ or less the width of the supraoccipital (1).
154. Supraoccipital, posterior surface, oval fossae that create an apparent vertical ridge on the midline: absent (0); present (1).
155. Supraoccipital, posttemporal fenestrae, shape of ventral margin: supraoccipital not raised under posttemporal fenestra (0); supraoccipital raised under posttemporal fenestra (1).
156. Basioccipital, height of ventral surface relative to occipital condyle: short, ventral surface does not lie below the level of the occipital condyle (0); tall, lies below occipital condyle but not to a level equivalent to the depth of the occipital condyle itself (1); very tall, lies one or more occipital condyle height below the level of the occipital condyle (2). ORDERED.
157. Basioccipital, median keel: absent (0); present (1).
158. Basioccipital, ventral foramen, position: just anterior to articular surface of occipital condyle (0); ventrally displaced, adjacent to dorsal extent of median keel (1).
159. Basioccipital, braincase floor, lateral wall of hindbrain region, orientation: strictly anterolateral (0); posteriorly strictly anterior, anteriorly anterolateral (1); entirely straight and separated by a discrete corner from the midbrain region of the braincase floor (2). ORDERED.
160. Basioccipital, anterior margin, shape: anteriorly pointed at midline (0); transverse (1).
161. Basioccipital, occipital condyle, surface, form: smoothly convex (0); marked by a shallow indentation (1).
162. Basisphenoid, ventral surface, excavation for Eustachian canal: absent or limited, basioccipital ventral surface is convex (0); extensive, widely visible in ventral view (1).
163. Basisphenoid, ventral exposure: extensive (0); limited by posterior growth of pterygoid (1).
164. Pterygoid, secondary choana, anteroposterior length: occupies more than half of anteroposterior length of palatal surface of pterygoid (0); occupies less than half of anteroposterior length of palatal surface of pterygoid (1).
165. Pterygoid, secondary choana, lateral choanal ridges, length: long, approaching or exceeding anterior level of secondary choanal aperture (0); moderate, extending about half length of secondary choana (1); short, extending about one quarter the length of the secondary choana (2). ORDERED.
166. Pterygoid, posterior projection of palatal lamina, contact with osseus interchoanal septum: absent (0); present, forming a pseudosuture (1).
167. Pterygoid, nasal passage, transverse ridges: absent (0), present (1).
168. Pterygoid, lateral choanal ridges, size: large, framing a fossa with the pterygoid alae (0); small, projecting slightly form palatal surface and not forming fossae with the pterygoid alae (1).
169. Pterygoid, palatal surface, neurovascular foramina at anterior level of secondary choana: absent (0); present (1).
170. Pterygoid, posterior margin, shape: straight (0); embayed (1).
171. Palatine, prefrontal process, position: in anterior half of palatine (0); at approximate anteroposterior midlevel of palatine (1); in posterior half of palatine (2). ORDERED.
172. Palatine, prefrontal process, neurovascular foramen in lateral side: unenclosed, present as a notch (0); fully enclosed by bone (1).
173. Palatine, anterolateral process, orientation: extends mediolaterally (0); extends anterolaterally (1).
174. Palatine, dorsal surface, anterior to prefrontal pillar: not pneumatized (0); pneumatized recesses present but not contained within their own large recess (1); pneumatized recesses present and extensive, and themselves housed within a large pneumatic recess (2). ORDERED.
175. Palatine, medial surface, posterior to prefrontal pillar: not pneumatized (0); pneumatized (1).
176. Palatine, prefrontal pillar, medial surface, pneumatization: absent (0); present (1).
177. Palatine, palatine bulla: absent (0); present (1).
178. Palatine, ventral surface, medial to suborbital fenestra, neurovascular foramina: absent (0), two foramina present (1).
179. Vomer, anterior prong, dorsal margin: marked by a concave step (0); straight to convex (1).
180. Vomer, posterior prong, pneumatic recess: absent (0); present (1).
181. Skull, shape, lateral view: domed over the orbit (0); flattened, orbital region approximately in line with skull table (1).
182. Skull, preorbital region, length: shorter than postorbital region (0); approximately same length as postorbital region (1); longer than postorbital region (2). ORDERED.
183. Skull, dermal bones, external surface, sculpturing: smooth (0); sculptured (1); extensively sculptured, with no area of smooth bone remaining (2). ORDERED.
184. Skull, basicranium, height: in-line with occipital condyle (0); extending far below occipital condyle, basicranium heavily verticalized (1).
185. Skull, quadrate condyles, posterior level: anterior to occipital surface (0); level with occipital surface (1), posterior to occipital surface (2). ORDERED.
186. Skull, supratemporal fenestra, shape: slitlike (0); oval (1); circular (2). ORDERED.
187. Jugal, anterior extension over maxilla: long, overlying 7 maxillary teeth (0); moderate, overlapping 6 maxillary teeth (1); short, overlying 5 or fewer maxillary teeth (2). ORDERED.
188. Skull, orbit, exposure in lateral view: wide (0); constricted (1); absent, completely obscured by jugal (2). ORDERED.
189. Mandible, anterior end, orientation of teeth: vertical (0); laterally splayed (1).
